# Rho GTPase signaling and mDia facilitate endocytosis via presynaptic actin

**DOI:** 10.1101/2023.10.12.561997

**Authors:** Kristine Oevel, Svea Hohensee, Atul Kumar, Irving Rosas-Brugada, Francesca Bartolini, Tolga Soykan, Volker Haucke

## Abstract

Neurotransmission at synapses is mediated by the fusion and subsequent endocytosis of synaptic vesicle membranes. Actin has been suggested to be required for presynaptic endocytosis but the mechanisms that control actin polymerization and its mode of action within presynaptic nerve terminals remain poorly understood. We combine optical recordings of presynaptic membrane dynamics and ultrastructural analysis with genetic and pharmacological manipulations to demonstrate that presynaptic endocytosis is controlled by actin regulatory diaphanous-related formins mDia1/3 and Rho family GTPase signaling. We show that impaired presynaptic actin assembly in the near absence of mDia1/3 and reduced RhoA activity is partly compensated by hyperactivation of Rac1. Inhibition of Rac1 signaling further aggravates impaired presynaptic endocytosis elicited by loss of mDia1/3. Our data suggest that interdependent mDia1/3-Rho and Rac1 signaling pathways cooperatively act to facilitate synaptic vesicle endocytosis by controlling presynaptic F-actin.

## INTRODUCTION

Synaptic transmission relies on the release of neurotransmitters by calcium-triggered exocytic fusion of synaptic vesicles (SVs) at specialized active zone release sites within presynaptic nerve terminals. Following fusion, compensatory endocytosis retrieves SV proteins and lipids from the presynaptic plasma membrane and SVs are reformed (Chanaday *et al*, 2019; Gan & Watanabe, 2018; Kononenko & Haucke, 2015; Saheki & De Camilli, 2012; Soykan *et al*, 2016), e.g. by clathrin-mediated vesicle budding (Kononenko *et al*, 2014; Watanabe *et al*, 2014) (but see (Wu *et al*, 2014) for evidence in favor of clathrin-independent SV reformation). Endocytosis relies on mechanical forces to enable membrane deformation and eventually fission (Engqvist-Goldstein & Drubin, 2003). In many biological systems, actin and actin-associated myosin motors appear to provide such mechanical force to facilitate vesicle formation (Anes *et al*, 2003; Marston *et al*, 2003). For example, the function of actin for endocytosis is well-established in yeast, whereas differential requirements for actin in distinct forms of endocytosis have been described in various types of mammalian cells including neurons (Boulant *et al*, 2011; Merrifield *et al*, 2005; Saffarian *et al*, 2009; Saheki & De Camilli, 2012). Pharmacological inhibition of actin assembly by latrunculin has been shown to cause the accumulation of endocytic intermediates at lamprey giant synapses (Shupliakov *et al*, 2002) and interfere with ultrafast endocytosis in response to single optical action potentials (APs) at hippocampal synapses (Watanabe *et al*, 2013; Watanabe *et al*., 2014; Wu & Chan, 2022) and with fast endocytosis at cerebellar mossy fiber boutons (Delvendahl *et al*, 2016). The same treatment does not hamper endocytosis of SV proteins at hippocampal synapses stimulated with trains of APs (Sankaranarayanan *et al*, 2003; Soykan *et al*, 2017). Genetic interference with actin function by conditional knockout (KO) of β- or γ-actin genes has suggested a crucial role for actin in all kinetically distinguishable forms of endocytosis (Wu *et al*, 2016) at hippocampal synapses and the calyx of Held, a fast giant synapse in the auditory brain stem (Borst & Soria van Hoeve, 2012). However, in the same preparations actin loss also affected SV exocytosis (Wu *et al*., 2016), suggesting a more general requirement of actin for presynaptic function. Consistently, actin has been shown to surround clusters of reserve pool SVs in lamprey (Bloom *et al*, 2003), to facilitate the replenishment of fast releasing vesicles in various models (Sakaba *et al*, 2013; Sakaba & Neher, 2003), to be involved in Tau pathology (Zhou *et al*, 2017), to induce spine growth and plasticity, (Cingolani & Goda, 2008) and to steer neuronal migration (Shinohara *et al*, 2012) among various other roles.

We have previously demonstrated that pharmacological interference with the function of formins, a group of Rho family-associated proteins that nucleate linear filamentous (F) actin, kinetically delays SV endocytosis at hippocampal boutons and blocks compensatory endocytosis at the calyx of Held in pre-hearing rats (Soykan *et al*., 2017). Capacitance measurements at the calyx of Held in post-hearing rodents suggest a less stringent requirement for formin-mediated actin assembly for endocytosis (Hori *et al*, 2022). The latter observation may reflect the operation of compensatory pathways for actin assembly, e.g. via Rac1 or Cdc42 Rho-family GTPases that promote branched actin networks (Eisenmann *et al*, 2005; Goode & Eck, 2007; Hodge & Ridley, 2016).

In contrast to the vast body of literature regarding the role of actin and actin-associated proteins at the postsynapse [(Cingolani & Goda, 2008; Colgan & Yasuda, 2014) and references therein], most notably the actin-rich mesh of the postsynaptic density of glutamatergic synapses, comparably little is known about the signaling pathways (e.g. via guanine nucleotide exchange factors and GTPase activating proteins for Rho family small GTPases (Muller *et al*, 2020)) that mediate actin assembly at the presynapse. Recent work has suggested that presynaptic Rac1 negatively regulates synaptic strength and release probability by altering SV priming and replenishment at central synapses (O’Neil *et al*, 2021) including the calyx of Held (Keine *et al*, 2022). These data suggest a negative regulatory role for Rac1 in SV exocytosis. The abundance of postsynaptic actin (Chen *et al*, 2020; Cingolani & Goda, 2008; Colgan & Yasuda, 2014) has hampered the analysis of the nanoscale localization of actin at presynaptic nerve terminals and of the actin-regulatory proteins that control its dynamics. KO of Dynamin, the main enzyme for endocytic membrane fission (Ferguson *et al*, 2007; Imoto *et al*, 2022; Raimondi *et al*, 2011), has been shown to cause the accumulation of F-actin at and around stalled endocytic intermediates in non-neuronal cells (Ferguson *et al*, 2009) but this phenotype seems less overt at hippocampal synapses (Raimondi *et al*., 2011). Hence, actin in addition to a possible local role at endocytic invaginations might serve a more general function in endocytosis at synapses, for example by modulating plasma membrane tension or contractility, by providing a restrictive membrane scaffold that promotes membrane fission by keeping endocytic invaginations under longitudinal tension (Boulant *et al*., 2011; Roux *et al*, 2006; Wu & Chan, 2022), or via formation of an F-actin ring around the active zone that mechanically couples exocytic membrane compression to endocytic pit formation (Ogunmowo *et al*, 2023).

Here we combine optical recordings of SV exo-/ endocytosis and ultrastructural analysis with genetic and pharmacological manipulations to show that interdependent mDia1/3-Rho and Rac1 signaling pathways cooperate to facilitate SV endocytosis by controlling presynaptic F-actin.

## RESULTS

### Actin dynamics and actin nucleating mDia1/3 proteins facilitate presynaptic endocytosis and SV recycling

Previous studies using pharmacological inhibitors of actin polymerization and depolymerization have yielded inconclusive, often conflicting data regarding the function of actin in SV endocytosis and recycling in different models (Bleckert *et al*, 2012; Del Signore *et al*, 2021; Piriya Ananda Babu *et al*, 2020; Sankaranarayanan *et al*., 2003; Shupliakov *et al*., 2002; Wu & Chan, 2022). In contrast, genetic ablation of β- or γ-actin genes in mouse neurons suggests that actin is required for all forms of endocytosis at several types of synapses (Wu *et al*., 2016). Prompted by these findings we revisited the role of actin in presynaptic endocytosis by analyzing the effects of impaired F-actin dynamics in the combined presence of the G-actin sequestering drug latrunculin A, the F-actin stabilizer jasplakinolide, and Y-27632, an inhibitor of ROCK kinase signaling. This cocktail was shown to preserve the existing cytoskeleton architechture while blocking actin assembly, disassembly and rearrangement (Peng *et al*, 2011). We optically recorded the stimulation-induced exo-endocytosis of the SV protein Synaptophysin (Syph) fused to a pH-sensitive super-ecliptic green fluorescent protein (pHluorin) (Miesenbock *et al*, 1998) that is de-quenched during exocytosis and undergoes re-quenching as SVs are internalized and re-acidified in hippocampal neurons at physiological temperature. Under these conditions (i.e. trains of APs, 37°C), SV endocytosis occurs on a timescale of > 10 seconds, e.g. a timescale that is slower than re-quenching of pHluorin due to reacidification (Lopez-Hernandez *et al*, 2022; Soykan *et al*., 2017). Hence, the decay of pHluorin signals can serve as a measure of the time course of SV endocytosis. Perturbation of actin dynamics in the combined presence of latrunculin A, jasplakinolide, and Y-27632 significantly slowed down the endocytic retrieval of exogenously expressed Syph-pHluorin (Figure 1A,B). Application of Y-27632 alone or a combination of latrunculin A and jasplakinolide had no effect on SV endocytosis kinetics, whereas combined use of Y-27632 together with jasplakinolide displayed a mild inhibitory effect (Figure 1- Supplement 1C-F). Impaired actin dynamics in the presence of latrunculin A, jasplakinolide, and Y-27632 did not impact the apparent level of SV exocytosis indicated by the maximal amplitude of Syph-pHluorin fluorescence (Figure 1-Supplement 1A). As optical imaging of pHluorin reporters may lead to artifacts, we analyzed the internalization kinetics of the endogenous vesicular γ-aminobutyric acid transporter (vGAT) using antibodies directed against its luminal domain coupled to the pH-sensitive fluorophore CypHer5E (Hua *et al*, 2011; Lopez-Hernandez *et al*., 2022). The cyanine-based dye CypHer5E is quenched at neutral pH but exhibits bright fluorescence when present in the acidic lumen of SVs and, thus can serve as a tracer for the exo-endocytic cycling of endogenous SV proteins. Perturbation of actin dynamics significantly delayed the endocytic retrieval of endogenous vGAT in response to train stimulation with 200 APs at physiological temperature (Figure 1C,D), consistent with the results from exogenously expressed Syph-pHluorin. vGAT exocytosis proceeded unperturbed (Figure 1-Supplement 1B). Hence, a dynamic actin cytoskeleton facilitates presynaptic endocytosis in hippocampal neurons.

**Figure 1.**
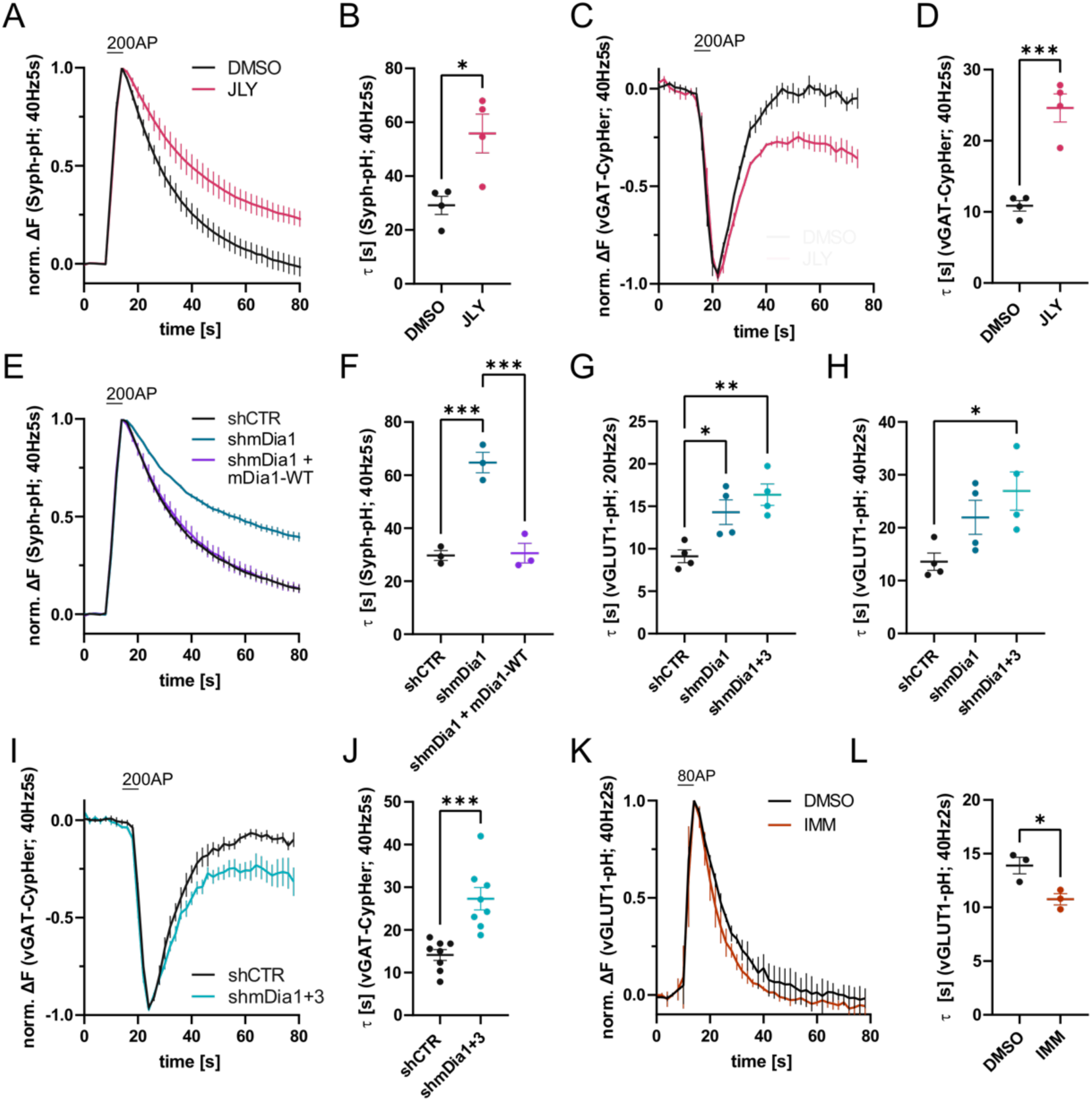
Actin dynamics and actin-nucleating mDia1/3 proteins facilitate SV endocytosis. **(A)** Averaged normalized Synaptophysin-pHluorin (Syph-pH) fluorescence traces from transfected hippocampal neurons stimulated with 200 APs (40 Hz, 5s) at physiological temperature (37.5°C). Neurons were treated with 0.1% DMSO or JLY cocktail (containing 8 µM Jasplakinolide, 5 µM Latrunculin A and 10 µM Y-27632) as indicated. Data shown represent the mean ± SEM. N = 4 independent experiments from n_DMSO_ = 23 videos; n_JLY_ = 36 videos. **(B)** Endocytic decay constants (τ) of Synaptophysin-pHluorin traces in A: τ_DMSO_ = 29.1 ± 3.4 s; τ_JLY_ = 55.8 ± 7.2 s; p < 0.05, two-tailed student’s t-test. Data shown represent mean ± SEM. **(C)** Averaged normalized bleach-corrected vGAT-CypHer fluorescence traces from hippocampal neurons treated with DMSO or JLY cocktail in response to 200 AP (40 Hz, 5s) stimulation. Data shown represent the mean ± SEM. N = 4 independent experiments from n_DMSO_ = 23 videos; n_JLY_ = 29 videos. **(D)** Endocytic decay constants of vGAT-CypHer traces in C: τ_DMSO_ = 10.9 ± 0.7 s, τ_JLY_ = 24.6 ± 2.0 s; p < 0.001, two-tailed unpaired student’s t-test. Data shown represent the mean ± SEM. **(E)** Averaged normalized Synaptophysin-pHluorin fluorescence traces from hippocampal neurons transfected with shRNA-encoding plasmids against no mammalian target (shCTR) or mDia1 (shmDia1) in response to 200 AP (40 Hz, 5s) stimulation. Neurons were co-transfected with mDia1-mCherry (mDia1-WT) or mCherry alone (shCTR & shmDia1) to exclude artefacts from overexpression. Data shown represent the mean ± SEM. N = 3 independent experiments from n_shCTR_ = 28 videos, n_shmDia1_ = 21 videos, n_shmDia1 + mDia1-WT_ = 21 videos. **(F)** Endocytic decay constants of Synaptophysin-pHluorin traces in E: τ_shCTR_ = 29.7 ± 1.9 s; τ_shmDia1_ = 64.7 ± 3.9 s; τ_shmDia1 + mDia1-WT_ = 30.6 ± 3.7 s; p_shCTR vs shmDia1_ < 0.001, p_shmDia1 vs shmDia1 + mDia1-WT_ < 0.001, one-way ANOVA with Tukey’s post-test. Data shown represent mean ± SEM. **(G)** Endocytic decay constants of averaged normalized vGLUT1-pHluorin fluorescence traces (Figure 1–Supplement 1I) from hippocampal neurons transduced with shCTR (τ_shCTR_ = 9.1 ± 0.8 s), shmDia1 (τ_shmDia1_ = 14.3 ± 1.5 s) or shmDia1+3 (τ_shmDia1+3_ = 16.4 ± 1.3 s) in response to 40 AP (20 Hz, 2s) stimulation (p_shCTR vs shmDia1_ < 0.05, p_shCTR vs shmDia1+3_ < 0.01, one-way ANOVA with Tukey’s post-test). Data shown represent mean ± SEM. N = 4 independent experiments from n_shCTR_ = 17 videos; n_shmDia1_ = 19 videos; n_shmDia1+3_ = 18 videos. **(H)** Endocytic decay constants of averaged normalized vGLUT1-pHluorin fluorescence traces (Figure 1–Supplement 1K) of neurons transduced with lentiviral vectors encoding shCTR (τ_shCTR_ = 13.6 ± 1.6 s), shmDia1 (τ_shmDia1_ = 22.0 ± 3.2 s) or shmDia1+3 (τ_shmDia1+3_ = 26.9 ± 3.6 s) in response to 80 AP (40 Hz, 2s) stimulation (p_shCTR vs shmDia1+3_ < 0.05, one-way ANOVA with Tukey’s post-test). Data shown represent mean ± SEM. N = 4 independent experiments from n_shCTR_ = 12 videos, n_shmDia1_ = 15 videos, n_shmDia1+3_ = 18 videos. **(I)** Averaged normalized bleach-corrected vGAT-CypHer fluorescence traces from hippocampal neurons transduced with shCTR or shmDia1+3 in response to 200 AP (40 Hz, 5s) stimulation. Data shown represent the mean ± SEM. N = 8 independent experiments from n_shCTR_ = 37 videos, n_shmDia1+3_ = 35 videos. **(J)** Endocytic decay constants of vGAT-CypHer traces in I: τ_shCTR_ = 14.1 ± 1.3 s; τ_shmDia1+3_ = 27.3 ± 2.6 s; p < 0.001, two-tailed unpaired student’s t-test. Data shown represent the mean ± SEM. **(K)** Averaged normalized vGLUT1-pHluorin fluorescence traces from transduced neurons in response to 80 AP (40 Hz, 2s) stimulation. Cells were treated with 0.1% DMSO or 10 µM mDia activator (IMM) in the imaging buffer. Data shown represent mean ± SEM. N = 3 independent experiments from n_DMSO_ = 18 videos; n_IMM_ = 16 videos. **(L)** Endocytic decay constants of vGLUT1-pHluorin traces in K: τ_DMSO_ = 14.9 ± 0.8 s; τ_IMM_= 9.8 ± 0.5 s; p < 0.05, two-tailed unpaired student’s t-test. Data shown represent mean ± SEM.

Previous studies have suggested that formins, in particular mDia1, promote SV endocytosis (Soykan *et al*., 2017), likely by inducing F-actin assembly (Ganguly *et al*, 2015). We confirmed that shRNA-mediated depletion of mDia1 from hippocampal neurons led to significantly delayed Syph-pHluorin endocytosis and re-acidification in response to stimulation with 200 APs. Re-expression of shRNA-resistant wild-type mDia1 fully restored normal endocytosis kinetics (Figure 1E,F). Impaired SV endocytosis in lentivirus-mediated mDia1-depleted neurons (Figure 1-Supplement 1H) was also observed, if SV exo-/ endocytosis was triggered by stimulation with 40 APs or 80 APs and probed with vesicular glutamate transporter 1 (vGLUT1)-pHluorin (Figure 1G,H and Figure 1-Supplement 1I,K). Exocytic fusion of vGLUT1-pHluorin-containing SVs was unaffected (Figure 1-Supplement 1G). Co-depletion of the closely related mDia3 isoform tended to further aggravate this phenotype (Figure 1G,H and Figure 1- Supplement 1I,K) without impacting the apparent number of fused SVs (Figure 1- Supplement 1J,L), but not vGAT exocytosis (Figure 1-Supplement 1M). Furthermore, mDia1/3 depletion significantly delayed the endocytic retrieval of endogenous vGAT (Figure 1I,J). Conversely, application of IMM-01, a small molecule activator of mDia-related formins that acts by relieving autoinhibition (Lash *et al*, 2013) (Figure 1-Supplement 1N), led to a moderate but significant acceleration of SV endocytosis monitored by vGLUT1-pHluorin (Figure 1K,L) but no change in apparent SV fusion (Figure 1- Supplement 1O). Ultrastructural analysis of hippocampal synapses from mDia1/3-depleted neurons by electron microscopy revealed a reduction in SV density compared to controls (Figure 2A,B). This phenotype was rescued upon prior silencing of endogenous neuronal network activity in the presence of tetrodotoxin (TTX) (Figure 2C and Figure 2-Supplement 1A,B). A similar, slightly more pronounced depletion of SVs was observed in hippocampal synapses from mDia1 knockout (KO) mice (Peng *et al*, 2007; Peng *et al*, 2003; Shinohara *et al*., 2012) (Figure 2D,E).

**Figure 2.**
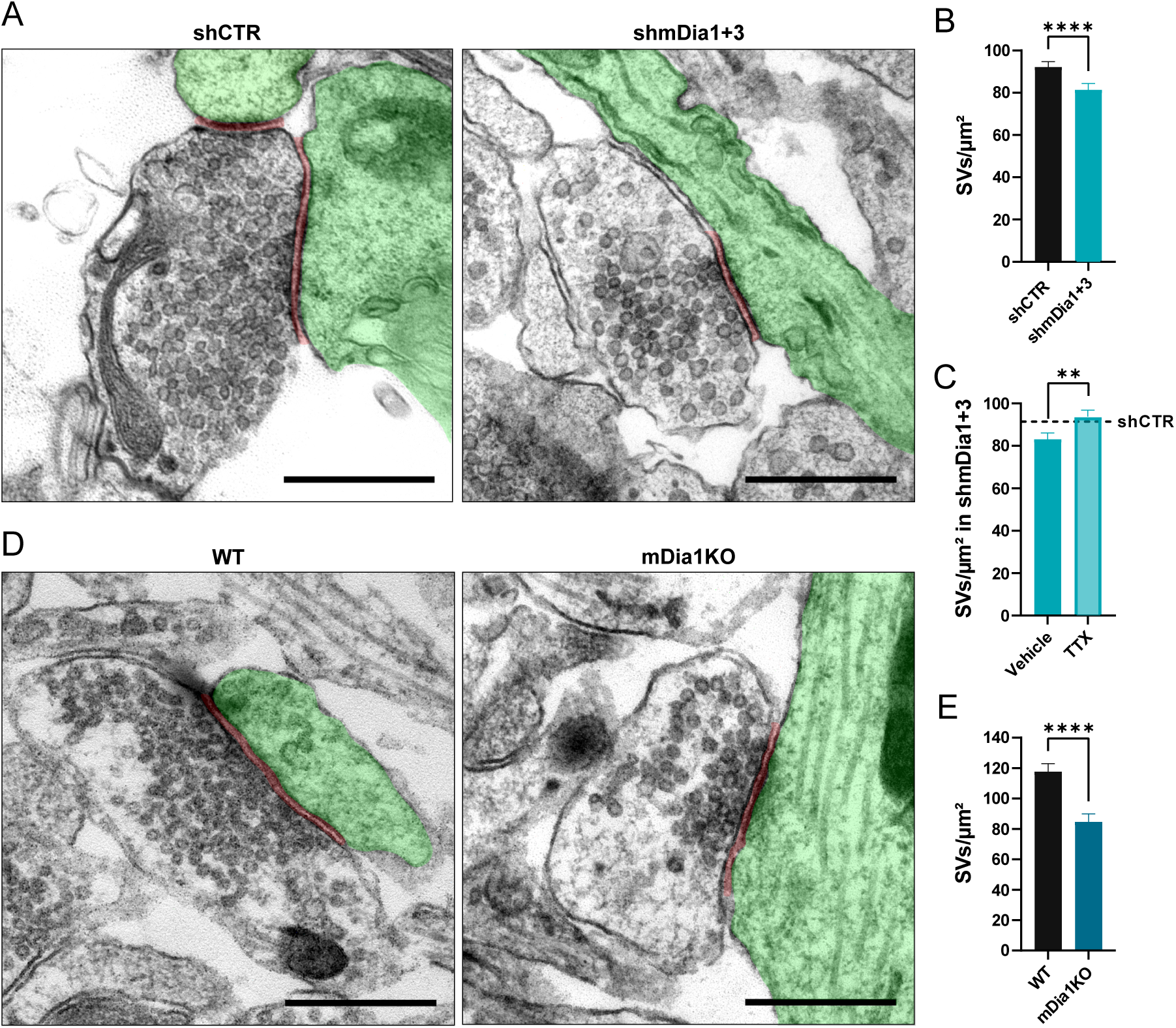
Loss of mDia1/3 causes an activity-dependent reduction of the SV pool. **(A)** Representative synaptic electron micrographs from hippocampal neurons transduced with lentiviruses encoding shCTR or shmDia1+3. Postsynapse and postsynaptic cleft are colored in green and maroon, respectively. Scale bar, 250 nm. **(B)** Average number of SVs per μm^2^ in boutons from shCTR (92.2 ± 2.5) and shmDia1+3 (81.4 ± 2.9; p < 0.0001, Mann-Whitney test) treated neurons. Data shown represent mean ± SEM. N = 3 independent experiments from n_shCTR_ = 326 synapses, n_shmDia1+3_ = 321 synapses. **(C)** Average number of SVs per μm^2^ in synaptic boutons from hippocampal neurons transduced with lentiviruses encoding shmDia1+3 and treated with 0.1 % Vehicle (10 µM NaOAc; 83.2 ± 2.9) or 1 µM Tetrodotoxin (TTX; 93.8 ± 3.1; p < 0.01, Mann-Whitney test) for 36 h before chemical fixation. Data shown represent mean ± SEM from two independent experiments and n_Vehicle_ = 225 synapses, n_TTX_ = 221 synapses. Representative synaptic electron micrographs are shown in Figure 2–Supplement 1A. The dotted line represents the average SV numbers/ μm^2^ in shCTR boutons treated with DMSO from the same experiments as a reference (Figure 2- Supplement 1A,B). **(D)** Representative electron micrographs of synapses in hippocampal neurons from WT or mDia1 KO mice. Postsynapse and postsynaptic cleft are colored in green and maroon, respectively. Scale bar, 250nm. **(E)** Average number of SVs per μm^2^ in WT (117.6 ± 5.3) and mDia1 KO (84.6 ± 5.2; p < 0.0001, Mann-Whitney test) boutons. Data shown represent the mean ± SEM from n_WT_ = 103, n_KO_= 96 synapses (N = 1).

These data are consistent with a role for actin dynamics (see Fig. 1) and actin nucleating mDia1/3 proteins in presynaptic endocytosis and SV recycling.

### mDia facilitates SV endocytosis by regulating presynaptic F-actin

mDia1 in addition to its Rho binding domain and a C-terminal actin-assembling formin homology 2 (FH2) domain comprises an unstructured N-terminal region involved in membrane binding (Eisenmann *et al*., 2005; Ramalingam *et al*, 2010) (Figure 3A). This architecture suggests that mDia specifically promotes the nucleation of unbranched actin filaments at membranes, for example to promote endocytosis at the presynaptic plasma membrane. To test this hypothesis, we assayed the ability of mutant mDia1 lacking its N-terminal membrane-binding region (mDia1-ΔN) (Ramalingam *et al*., 2010) to rescue defective SV endocytosis in hippocampal neurons depleted of endogenous mDia1. Mutant mDia1-ΔN displayed reduced association with membranes in transfected HEK293T cells (Figure 3-Supplement 1A,B) and was unable to restore normal kinetics of Syph-pHluorin endocytosis in hippocampal neurons transfected with mDia1-specific microRNA (Figure 3B and Figure 3-Supplement 1C). Hence, mDia may act on SV endocytosis by specifically promoting actin assembly at presynaptic membranes. We probed this model at several levels. We analyzed the nanoscale localization and distribution of mDia1 via multicolor time-gated stimulated emission depletion (gSTED) microscopy. Hippocampal neurons were stained for mDia1, the presynaptic active zone marker Bassoon, and post-synaptic Homer1. The distributions of all three proteins along line profiles perpendicular to the synaptic cleft (i.e. the space between Bassoon and Homer1 clusters) were analyzed (Gerth *et al*, 2017) (Figure 3-Supplement 1H). This analysis revealed that endogenous mDia1 is primarily localized to the presynaptic compartment, where it is concentrated at or very close to the plasma membrane (Figure 3C,D), i.e. < 50 nm away from the presynaptic membrane (see (Dani *et al*, 2010)). Prompted by these results we dissected the interactome of mDia1 by unbiased quantitative proteomic analyses of immunoprecipiates from detergent-extracted mouse synaptosomes (Figure 3-Supplement 1D). In addition to actin, we identified a variety of additional actin-regulatory proteins including Myosin IIB (Chandrasekar *et al*, 2013) and the small GTPase Rac1 (Figure 3E). We confirmed the association of mDia1 with actin and Myosin IIB (Figure 3-Supplement 1E) and found Myosin IIB to be concentrated at or near the presynaptic plasma membrane (Figure 3-Supplement 1F,G) akin to the localization of mDia1. The association of mDia1 with Myosin IIB is consistent with the inhibitory effects of the Myosin II inhibitor blebbistatin on SV endocytosis (Chandrasekar *et al*., 2013; Soykan *et al*., 2017). Interestingly, we also observed an enrichment of several endocytic proteins previously implicated in SV recycling in mDia1 immunoprecipitates. These include the lipid phosphatase Synaptojanin1, the bin-amphiphysin-rvs (BAR) domain proteins Amphiphysin, Endophilin A1, and PACSIN1/2 and, the GTPase Dynamin1 (Chanaday *et al*., 2019; Gan & Watanabe, 2018; Saheki & De Camilli, 2012; Soykan *et al*., 2016) (Figure 3E) as confirmed by immunoblotting (Figure 3F). Based on the association of mDia1 with endocytic proteins including Dynamin1, we hypothesized that mDia1 might get stalled at endocytic intermediates that accumulate under conditions of perturbed Dynamin function (Ferguson *et al*., 2009; Raimondi *et al*., 2011). Application of the partially non-selective Dynamin inhibitor Dynasore (Macia *et al*, 2006) (Figure 3C,H) or expression of dominant-negative mutant Dynamin1 K44A (Figure 3G,I) led to a significant accumulation of mDia1 at presynaptic sites marked by Bassoon. mDia1 levels at the postsynapse were also increased (Figure 3-Supplement 1I). Dynasore did not impact the levels (Figure 3- Supplement 1J) or distribution of Bassoon or Homer1.

**Figure 3.**
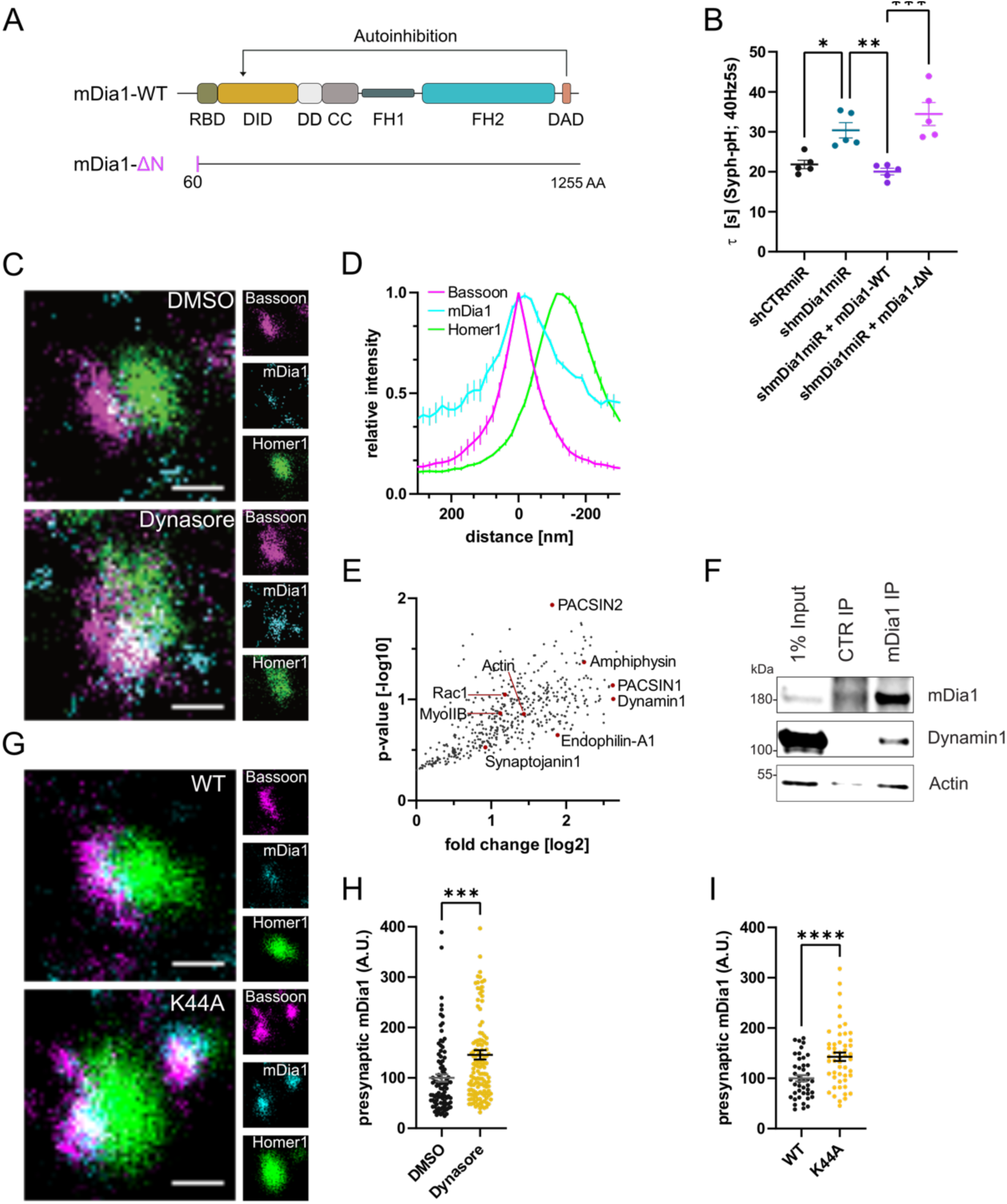
mDia1 associates with endocytic proteins and localizes to presynaptic sites. **(A)** Schematic representation of functional domains of mDia1. Rho-binding domain (RBD), Diaphanous inhibitory domain (DID), Dimerization domain (DD), Coiled coil domain (CC), Formin homology domain 1 (FH1), Formin homology domain 2 (FH2), Diaphanous autoinhibitory domain (DAD). The unstructured N-terminus (first 60 amino acids) contains three basic stretches and was truncated in the ΔN mutant. **(B)** Endocytic decay constants of Synaptophysin-pHluorin traces (Figure 3-Supplement 1C) from hippocampal neurons transfected with shRNAmiR against no mammalian target (shCTRmiR) or mDia1 (shmDia1miR) in response to 200 AP (40 Hz, 5s) stimulation. For rescue experiments, neurons were co-transfected with plasmids encoding mDia1-WT-mCherry (τ_shmDia1miR + mDia1-WT_ = 20.0 ± 0.8 s), mDia1-ΔN-mCherry (τ_shmDia1miR + mDia1-ΔN_ = 34.5 ± 2.9 s) or mCherry alone (τ_shCTRmiR_ = 21.8 ± 1.1 s, τ_shmDia1miR_ = 30.4 ± 1.9 s) to exclude artefacts from overexpression (p_shCTRmiR vs shmDia1miR_ < 0.05; p_shmDia1miR vs shmDia1miR + mDia1-WT_ < 0.01; p_shmDia1miR + mDia1-WT vs shmDia1miR + mDia1-ΔN_ < 0.01, one-way ANOVA with Tukey’s post-test). Data shown represent mean ± SEM. N = 5 independent experiments from n_shCTRmiR_ = 41 videos, n_shmDia1miR_ = 51 videos, n_shmDia1miR + mDia1-WT_ = 35 videos, n_shmDia1miR + mDia1-ΔN_ = 37 videos. **(C)** Representative three-channel time-gated STED images of synapses from hippocampal cultures treated with 0.1% DMSO or 80 μM Dynasore for 10 min before fixation and immunostained for Bassoon (presynaptic marker, magenta), mDia1 (cyan) and Homer1 (postsynaptic marker, green). Scale bar, 250 nm. **(D)** Averaged normalized line profiles for synaptic distribution of mDia1 and Homer1 relative to Bassoon (Maximum set to 0 nm). Data represent mean ± SEM. N = 3 independent experiments from n = 235 synapses. **(E)** Volcano plot of proteins associating with synaptic mDia1 analyzed by label-free proteomics of anti-mDia1 versus control (CTR) immunoprecipitates from detergent-extracted mouse synaptosomes (P2’ fraction). The logarithmic ratios of protein intensities are plotted against negative logarithmic p-values derived from two-tailed student’s t-test. N = 3 independent experiments. Each dot represents one protein/gene. Selected cytoskeletal hits include: Actin, Myosin IIB (MyoIIB) and Rac1. Selected endocytic hits include Amphiphysin (p < 0.05), Dynamin1, Endophilin-A1, PACSIN1, PACSIN2 (p < 0.05) and Synaptojanin1. **(F)** Endogenous immunoprecipitation of mDia1 from detergent-extracted mouse synaptosomes (P2’ fraction) using mDia1-specific antibodies. Immunoprecipitates were analyzed by immunoblotting for mDia1, Dynamin1 and β-Actin. **(G)** Representative three-channel time-gated STED images of synapses from hippocampal cultures transduced with wildtype Dynamin1 (WT) or GTPase-deficient Dynamin1 (K44A) in response to 200 AP (40 Hz, 5s) stimulation. Cells were immunostained for Bassoon (magenta), mDia1 (cyan) and Homer1 (green). Scale bar, 250 nm. **(H)** Presynaptic mDia1 levels in synapses treated with 0.1% DMSO (100 ± 7.3) or 80 µM Dynasore (145.8 ± 9.3; p = 0.0001; one sample Wilcoxon test) for 10 min in response to 200 AP (40 Hz, 5s) stimulation. Absolute line profiles of mDia1 overlapping with Bassoon (presynapse) distribution were integrated. Data shown are normalized to DMSO (set to 100) and expressed as mean ± SEM. N = 3 independent experiments from n_DMSO_ = 92 synapses, n_Dynasore_ = 135 synapses. **(I)** Presynaptic mDia1 levels in synapses from hippocampal neurons transduced with wildtype Dynamin1 (WT; 100 ± 6.2) or GTPase-deficient Dynamin1 (K44A; 142.9 ± 8.3, p < 0.0001, one sample Wilcoxon test) in response to 200 AP (40 Hz, 5s) stimulation. Line profiles of mDia1 overlapping with Bassoon distribution were integrated. Data shown are normalized to Dynamin1-WT (set to 100) and expressed as mean ± SEM. N = 2 independent experiments from n_WT_ = 43 synapses, n_K44A_ = 51 synapses.

Based on these results we next investigated the relationship between mDia1/3 and presynaptic actin. Multicolor gSTED imaging confirmed that F-actin labeled by phalloidin was highly abundant at the postsynaptic compartment. In addition, F-actin was also visible, albeit at reduced levels, in presynaptic boutons marked by Bassoon (Figure 4A,B). Inhibition of Dynamin in the presence of Dynasore (Figure 4C and Figure 4-Supplement 1A) or expression of Dynamin1 K44A (Figure 4-Supplement 1B,C) significantly elevated presynaptic F-actin levels within the Bassoon area. These data are consistent with a role for presynaptic actin in SV endocytosis. We directly tested the role of mDia in presynaptic actin nucleation and actin-dependent SV endocytosis in several ways. First, we generated a mutant version of mDia1 that lacks the ability to nucleate F-actin due to site-specific inactivation of its FH2 domain (Daou *et al*, 2014; Higashi *et al*, 2008; Xu *et al*, 2004). We found that actin polymerization-defective mutant mDia1 (K994A) failed to restore delayed vGLUT1-pHluorin endocytosis in mDia1-depleted hippocampal neurons (Figure 4D and Figure 4-Supplement 1D). Second, we analyzed the effects of depleting hippocampal neurons of mDia1/3 on presynaptic F-actin. We found that loss of mDia1/3 significantly reduced the amount of detectable F-actin within the Bassoon area (Figure 4A,E). To unequivocally distinguish between presynaptic actin at exo-/ endocytic sites and the abundant pool of postsynaptic actin associated with the postsynaptic density, we used the ORANGE strategy for CRISPR-based genome-engineering (Willems *et al*, 2020) to generate eGFP-β-actin knock-in (KI) hippocampal neurons. Due to the low knockin efficiency, this strategy allowed us to visualize endogenously tagged eGFP-β-actin by 2-color gSTED imaging exclusively in presynaptic boutons marked by vGLUT1. This assay confirmed the significant reduction in endogenous β-actin at presynapses from mDia1/3-depleted neurons (Figure 4F,G). Third, we reasoned that if loss of presynaptic actin in mDia-depleted neurons was causal for the observed defect in SV endocytosis, pharmacological stabilization of F-actin might rescue the endocytic phenotype. Treatment of mDia1/3-depleted neurons with jasplakinolide indeed fully restored the kinetics of SV endocytosis monitored by vGLUT1-pHluorin (Figure 4H and Figure 4-Supplement 1H) and rescued presynaptic F-actin levels to that of controls (actin levels normalized to shCTR + DMSO set to 100: shCTR + DMSO = 100 ± 6.3; shmDia1+3 + DMSO = 47.7 ± 4.3; shCTR + Jasp = 150.6 ± 11.9; shmDia1+3 + Jasp = 94.3 ± 11.5) (Figure 4-Supplement 1G).

**Figure 4.**
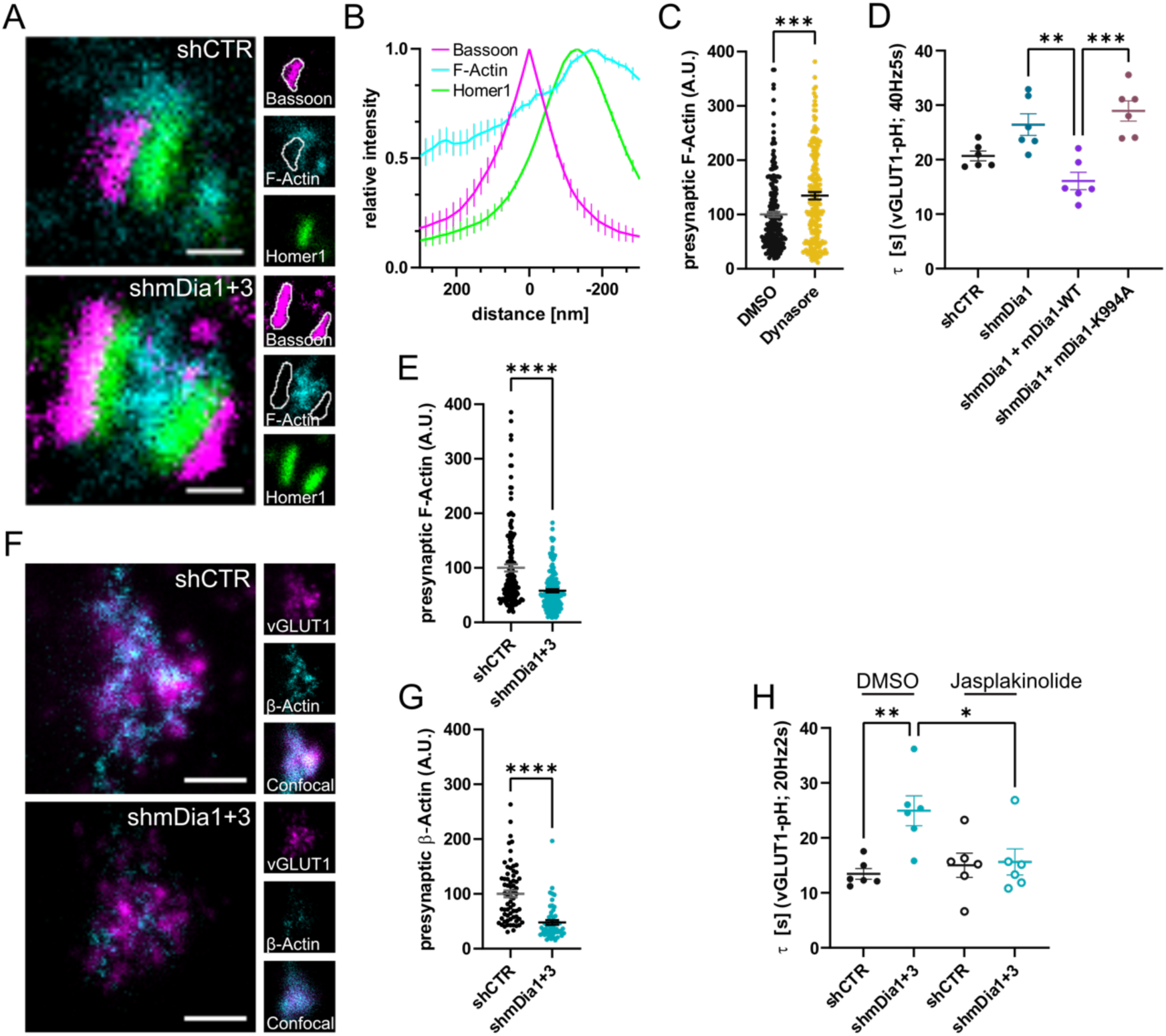
mDia facilitates SV endocytosis by regulating presynaptic F-actin. **(A)** Representative three-channel time-gated STED images of synapses from hippocampal cultures transduced with shCTR or shmDia1+3, fixed and immunostained for Bassoon (presynaptic marker, magenta), F-Actin (cyan) and Homer1 (postsynaptic marker, green). Scale bar, 250 nm. **(B)** Averaged normalized line profiles for synaptic distribution of F-Actin and Homer1 relative to Bassoon (Maximum set to 0 nm). Data are expressed as mean ± SEM. N = 4 independent experiments from n = 154 synapses. **(C)** Presynaptic F-Actin levels in synapses treated with 0.1% DMSO (100 ± 4.8) or 80 µM Dynasore (134.7 ± 6.8; p = 0.001, one sample Wilcoxon test) for 10 min before fixation (Represenative images in Figure 4–Supplement 1A). Cells were immunostained for Bassoon (magenta), F-Actin (cyan), and Homer1 (green). Absolute line profiles of F-Actin overlapping with Bassoon (presynapse) distribution were integrated. Data shown are normalized to DMSO (set to 100) and expressed as mean ± SEM. N = 3 independent experiments from n_DMSO_ = 207 synapses, n_Dynasore_ = 211 synapses. **(D)** Endocytic decay constants of vGLUT1-pHluorin traces from hippocampal neurons transduced with lentiviral particles encoding shCTR (τ_shCTR_ = 20.7 ± 0.9 s) or shmDia1 (τ_shmDia1_ = 26.4 ± 2.0 s) in response to 200 AP (40 Hz, 5s) stimulation. For rescue experiments, neurons were co-transduced with lentiviruses encoding mDia1-WT-SNAP (τ_shmDia1 + mDia1-WT_ = 16.1 ± 1.9 s) or mDia1-K994A-SNAP (τ_shmDia1 + mDia1-K994A_ = 29.0 ± 1.9 s) (p_shmDia1 vs shmDia1 + mDia1-WT_ < 0.01; p_shmDia1 + mDia1-WT vs shmDia1 + mDia1-K994A_ < 0.001, one-way ANOVA with Tukey’s post-test). Data are expressed as mean ± SEM. N = 6 independent experiments from n_shCTR_ = 21 videos; n_shmDia1_ = 21 videos, n_shmDia1 + mDia1-WT_ = 16 videos, n_shmDia1 + mDia1-K994A_ = 19 videos. **(E)** Presynaptic F-Actin levels in synapses from hippocampal cultures transduced with shCTR (100 ± 6.4) or shmDia1+3 (58.1 ± 2.9; p < 0.001, one sample Wilcoxon test). Line profiles of F-Actin overlapping with Bassoon (presynapse) distribution were integrated. Data shown are normalized to shCTR (set to 100) and expressed as mean ± SEM. N = 4 independent experiments from n_shCTR_ = 155 synapses, n_shmDia1+3_ = 158 synapses. **(F)** Representative confocal and two-channel time-gated STED images of endogenous β-Actin (cyan) in vGLUT1 (magenta) positive boutons from hippocampal neurons transduced with lentiviruses encoding shCTR or shmDia1+3. Scale bar, 250 nm. **(G)** Analysis of presynaptic endogenous β-Actin levels in vGLUT1 positive boutons from shCTR (100 ± 6.3) and shmDia1+3 (47.7 ± 4.3; p < 0.0001 one sample Wilcoxon test) transduced neurons. β-Actin STED mean intensity was measured using confocal vGLUT1 signal as a mask. Data shown are normalized to shCTR (set to 100) and expressed as mean ± SEM from two independent experiments and n_shCTR_ = 67 synapses, n_shmDia1+3_ = 53 synapses. **(H)** Endocytic decay constants of vGLUT1-pHluorin traces (Figure 4–Supplement 1H) for neurons transduced with shCTR or shmDia1+3 in response to 40 AP (20 Hz, 2s) stimulation. Neurons were pre-incubated with 0.1 % DMSO or 1 µM Jasplakinolide (Jasp) for 30 min in the media before imaging (τ_shCTR + DMSO_ = 13.4 ± 1.0 s, τ_shCTR + Jasp_ = 15.0 ± 2.2 s, τ_shmDia1+3 + DMSO_ = 25.0 ± 2.7 s, τ_shmDia1+3 + Jasp_ = 15.6 ± 2.4 s; p_shCTR vs shmDia1+3_ < 0.01; p_shmDia1+3 + DMSO vs shmDia1+3 + Jasp_ < 0.05, one-way ANOVA with Tukey’s post-test). Data are expressed as mean ± SEM. N = 6 independent experiments from n_shCTR + DMSO_ = 32 videos, n_shmDia1+3 + DMSO_ = 35 videos, n_shCTR + Jasp_ = 33 videos; n_shmDia1+3 + Jasp_ = 34 videos).

We conclude that mDia1/3 facilitates SV endocytosis by regulating presynaptic F-actin.

### mDia1/3-Rho and Rac1 signaling pathways cooperatively act to facilitate presynaptic endocytosis

In non-neuronal cells mDia proteins have been shown to be disinhibited by active Rho family GTPases, in particular RhoA (Otomo *et al*, 2005) (Figure 5A and Figure 1-Supplement 1N). We reasoned that Rho proteins might serve a similar function at presynaptic nerve terminals in hippocampal neurons. Multicolor gSTED imaging revealed that endogenous RhoA was mostly located within the presynaptic compartment (Figure 5B,C). Interference with Rho function by co-expression of dominant-negative (DN) variants of RhoA and RhoB (Figure 5D and Figure 5- Supplement 1A) or co-depletion of these Rho isoforms (Figure 5-Supplement 1C) delayed SV endocytosis in response to stimulation with 200 APs without impacting the apparent levels of exocytosis (Figure 5- Supplement 1B,D). These results are consistent with a model in which active Rho promotes mDia function and, thereby F-actin nucleation.

**Figure 5.**
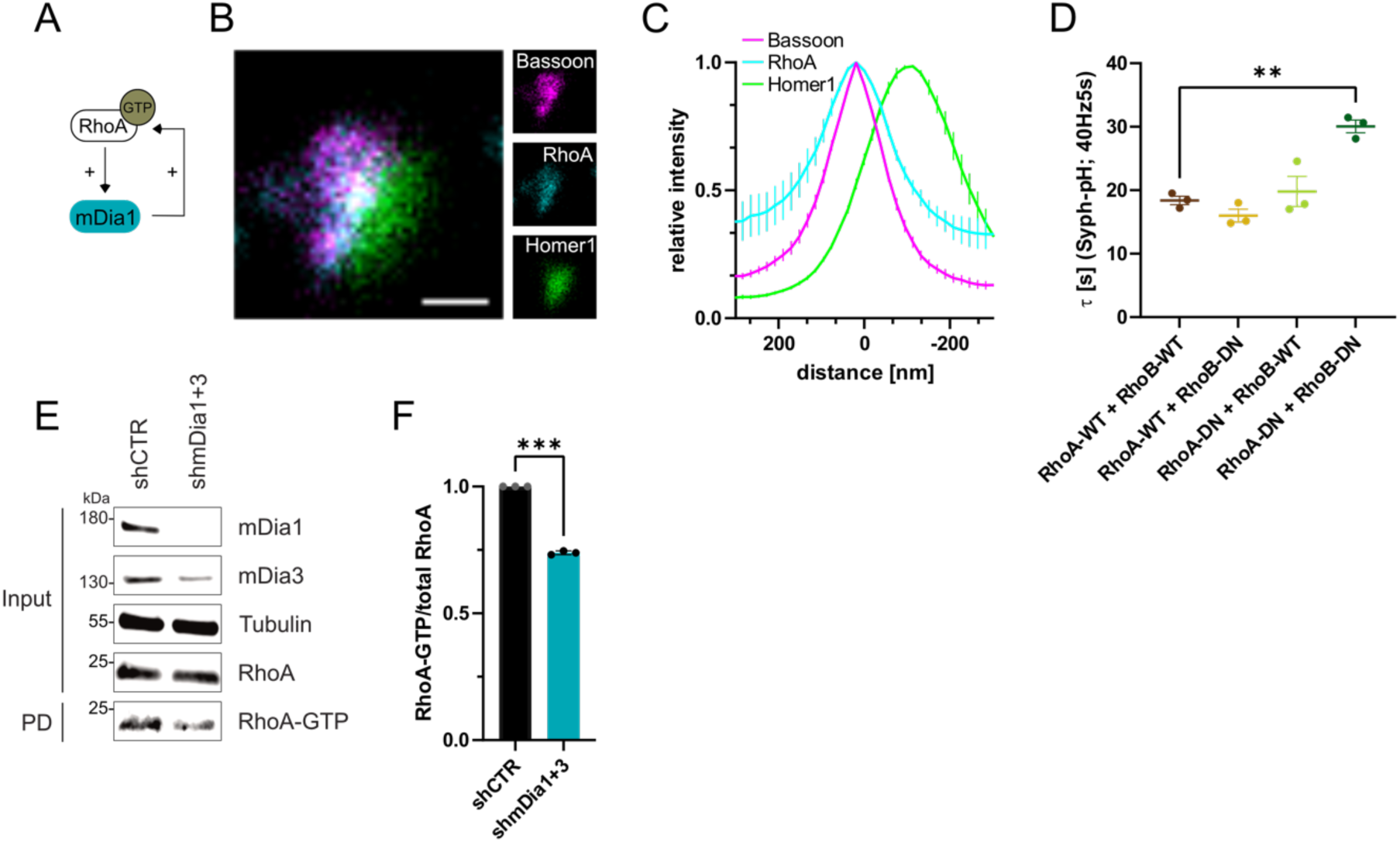
RhoA/B facilitate presynaptic endocytosis and are regulated by mDia1/3. **(A)** Schematic representation of activation of mDia1 by RhoA-GTP and positive feedback loop of mDia1 on RhoA-GTP levels through GEF stimulation. **(B)** Representative three-channel time-gated STED image of a synapses from hippocampal cultures, fixed and immunostained for Bassoon (magenta), RhoA (cyan) and Homer1 (green). Scale bar, 250 nm. **(C)** Averaged normalized line profiles for synaptic distribution of RhoA and Homer1 relative to Bassoon (Maximum set to 0 nm). Data are expressed as mean ± SEM. N = 5 independent experiments from n = 230 synapses. **(D)** Endocytic decay constants of averaged normalized Synaptophysin-pHluorin fluorescence traces (Figure 5–Supplement 1A) in response to 200 AP (40 Hz, 5s) stimulation. Neurons were transfected with the annotated combinations of plasmids encoding wild-type (WT) or dominant-negative (DN, T19N mutation) RhoA and RhoB (τ_RhoA-WT + RhoB-WT_ = 18.4 ± 0.7 s, τ_RhoA-WT + RhoB-DN_ = 16.0 ± 1.0 s, τ_RhoA-DN + RhoB-WT_ = 19.8 ± 2.4 s, τ_RhoA-DN + RhoB-DN_ = 30.1 ± 1.0 s; p_RhoA-WT + RhoB-WT vs RhoA-DN + RhoB-DN_ < 0.01, one-way ANOVA with Tukey’s post-test). Data shown represent mean ± SEM. N = 3 independent experiments from n_RhoA-WT + RhoB-WT_ = 21 videos, n_RhoA-DN + RhoB-WT_ = 31 videos, n_RhoA-WT + RhoB-DN_ = 23 videos, n_RhoA-DN + RhoB-DN_ = 22 videos. **(E)** Analysis of RhoA activity by RhoA-GTP pulldown (PD) from whole-cell lysates (input) of mouse hippocampal neurons expressing shCTR or shmDia1+3 using immobilized Rhotekin as a bait. Samples were analyzed by immunoblotting for mDia1, mDia3, RhoA and Tubulin using specific antibodies. Input, 10% of material used for the pulldown. Contrast of pulldown and input blots was seperatly adjusted for visualisation purposes. **(F)** Densitometric quantification of RhoA-GTP normalized to total RhoA levels (input) in lysates from neurons transduced with shCTR or shmDia1+3 (0.7 ± 0.0, p < 0.001, one sample t-test) from immunoblots exemplified in E. Values for shCTR were set to 1. Data are expressed as mean ± SEM from N = 3 independent experiments.

F-actin dynamics are known to be controlled by interdependent signaling networks characterized by feedback regulation between components within the same pathway and between key regulatory switches (e.g. distinct Rho/Rac family members) that drive distinct forms of F-actin (see below) (Lawson & Ridley, 2018). For example, active mDia is known to stimulate RhoA activation (Kitzing *et al*, 2007) (Figure 5A). Consistently, we found that RhoA activity was reduced in hippocampal neurons depleted of mDia1/3 (Figure 5E,F). Work in non-neuronal cells has further revealed that RhoA activity antagonizes activation of Rac1 (Chauhan *et al*, 2011), a key factor for promoting the formation of branched F-actin networks at the cell cortex (Hodge & Ridley, 2016; Lawson & Ridley, 2018) (Figure 6A). We hypothesized that reciprocally interdependent Rho/ Rac1 signaling might control presynaptic F-actin assembly and, thereby, SV endocytosis. In agreement with the alleged antagonistic regulation of RhoA and Rac1 function (Lawson & Ridley, 2018), we found Rac1 activity to be significantly elevated in hippocampal neurons depleted of mDia1/3 (Figure 6B,C), e.g. under conditions of reduced Rho-GTP levels. A similar increase in active Rac1 levels was observed upon pharmacological inhibition of RhoA/B in the presence of Rhosin (Figure 6-Supplement 1A). Multicolor gSTED imaging showed that Rac1 was equally distributed between pre- and postsynaptic compartments in unperturbed hippocampal neurons (Figure 6D,E). Elevated active Rac1 levels might conceivably ameliorate presynaptic endocytic phenotypes elicted by mDia1/3 loss. In support of this hypothesis, we found that selective pharmacological inhibition of Rac1 reduced presynaptic actin levels (Figure 6- Supplement 1B,C) and caused a delay in the endocytic retrieval of endogenous vGAT (Figure 6F,G). Moreover, Rac1 inhibition appeared to further aggravate impaired vGAT endocytosis in mDia1/3-depleted neurons, although the effect remained below statistical significance. Expression of constitutively active GTP-locked Rac1 (Rac1-CA) restored normal Syph-pHluorin endocytosis kinetics in mDia1/3-depleted neurons, whereas Syph-pHluorin endocytosis was delayed upon expression of dominant-negative Rac1 (Rac1-DN) and exacerbated the endocytic phenotype of mDia1/3 loss (Figure 6H and Figure 6-Supplement 1E). No overt effect of Rac1 inhibition or overexpression of either Rac1 form on SV exocytosis was observed (Figure 6-Supplement 1D,F). Perturbation of the related Cdc42 protein, another actin regulatory GTPase (Lawson & Ridley, 2018) found at synapses (Figure 6-Supplement 1G-I), did not significantly affect the kinetics of SV endocytosis in control or mDia1/3-depleted neurons (Figure 6- Supplement 1J,K). These data suggest that interdependent mDia1/3 and Rac1-based pathways control SV endocytosis via presynaptic actin.

**Figure 6.**
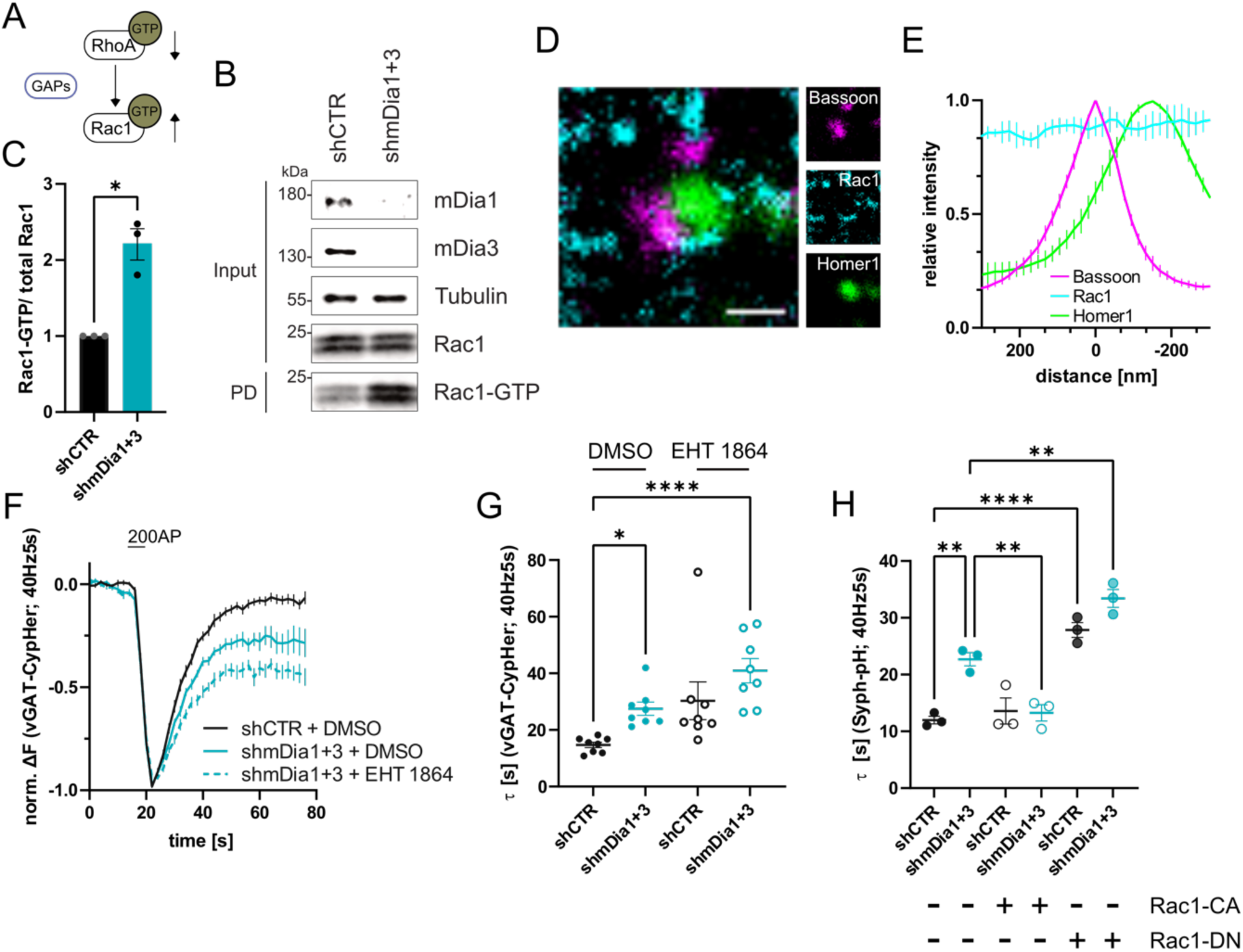
mDia1/3-Rho and Rac1 signaling facilitates presynaptic endocytosis. **(A)** Schematic of the interplay between RhoA and Rac1 signaling via GTPase activating proteins (GAPs) common for RhoA and Rac1. **(B)** Analysis of Rac1 activity by Rac1-GTP pulldown (PD) from whole-cell lysates (input) of mouse hippocampal neurons expressing shCTR or shmDia1+3 utilizing immobilized PAK as a bait. Samples were analyzed by immunoblotting for mDia1, mDia3, Rac1 and Tubulin using specific antibodies. Input, 10% of material used for the pulldown. Contrast of pulldown and input blots was seperatly adjusted for visualisation purposes. **(C)** Densitometric quantification of Rac1-GTP normalized to total Rac1 levels (input) in lysates from neurons transduced with shCTR or shmDia1+3 (2.2 ± 0.2; p < 0.05, one sample t-test) from immunoblots exemplified in (B). Values for shCTR were set to 1. Data are expressed as mean ± SEM from N = 3 independent experiments. **(D)** Representative three-channel time-gated STED image of a synapses from hippocampal cultures, fixed and immunostained for Bassoon (magenta), Rac1 (cyan) and Homer1 (green). Scale bar, 250 nm. **(E)** Averaged normalized line profiles for synaptic distribution of Rac1 and Homer1 relative to Bassoon (Maximum set to 0 nm). Data represent mean ± SEM. N = 3 independent experiments from n = 79 synapses. **(F)** Averaged normalized vGAT-CypHer fluorescence traces for neurons transduced with shCTR or shmDia1+3 in response to 200 AP (40 Hz, 5 s) stimulation. Cells were acutely treated with 0.1 % DMSO or 10 µM Rac1 Inhibitor (EHT 1864) in the imaging buffer. Data shown represent the mean ± SEM. N = 8 independent experiments from n_shCTR + DMSO_ = 46 videos, n_shmDia1+3 + DMSO_ = 45 videos, n_shCTR + EHT 1864_ = 42 videos, n_shmDia1+3 + EHT 1864_ = 43 videos. **(G)** Endocytic decay constants of vGAT-CypHer traces in F: τ_shCTR + DMSO_ = 14.7 ± 0.9 s, τ_shmDia1+3 + DMSO_ = 27.5 ± 2.3 s, τ_shCTR + EHT 1864_ = 30.3 ± 6.7 s, τ_shmDia1+3 + EHT 1864_ = 41.0 ± 4.3 s; p_shCTR + DMSO vs shmDia1+3 + DMSO_ < 0.05, p_shCTR + DMSO vs shmDia1+3 + EHT 1864_ < 0.0001, Kruskal-Wallis test with Dunn’s post-test. Data represent mean ± SEM. **(H)** Endocytic decay constants of Synaptophysin-pHluorin traces (Figure 6–Supplement 1E) of neurons transduced with shCTR (τ_shCTR_ = 12.0 ± 0.7 s) or shmDia1+3 (τ_shmDia1+3_ = 22.7 ± 2.0 s) and transfected with constitutively active Rac1 (Rac1-CA; Q61L variant; τ_shCTR + Rac1-CA_ = 13.6 ± 1.2 s, τ_shmDia1+3 + Rac1-CA_ = 13.3 ± 1.4 s) or dominant negative Rac1 (Rac1-DN; T17N variant; τ_shCTR + Rac1-DN_ = 27.8 ± 1.3 s, τ_shmDia1+3 + Rac1-DN_= 33.4 ± 1.6 s) in response to 200 AP (40 Hz, 5s) stimulation (p_shCTR vs shmDia1+3_ < 0.01; p_shCTR vs shCTR + Rac1-DN_ < 0.0001, p_shCTR vs shmDia1+3 + Rac1-DN_ < 0.01, p_shmDia1+3 vs shmDia1+3 + Rac1-DN_ < 0.01, one-way ANOVA with Tukey’s post-test). Data are expressed as mean ± SEM. N = 3 independent experiments from n_shCTR_ = 12 videos, n_shmDia1+3_ = 23 videos; n_shCTR + Rac1-CA_ = 10 videos, n_shmDia1+3 + Rac1-CA_ = 14 videos, n_shCTR + Rac1-DN_ = 9 videos; n_shmDia1+3 + Rac1-DN_ = 13 videos.

We probed this model by analyzing the ultrastructure of synapses from hippocampal neurons depleted of mDia1/3 or following inhibition of Rac1 activity. Synapses from mDia1/3- depleted neurons displayed an accumulation of non-coated plasma membrane invaginations that were found both in the vicinty of the active zone as well as distal from the synaptic contact area (Figure 7A,B). Moreover, we observed elevated numbers of endosome-like vacuoles (ELVs) (Figure 7A,C) that might serve as a donor membrane for SV reformation (Kononenko & Haucke, 2015; Kononenko *et al*., 2014; Watanabe *et al*., 2014) as suggested by the observed reduction in SV numbers in mDia-depleted neurons (compare Figure 2). A prominent accumulation of plasma membrane invaginations (Figure 7-Supplement 1A) and endosome-like vacuoles (Figure 7- Supplement 1B) was similarly found in hippocampal neurons from mDia1 KO mice. Inhibition of Rac1 function by EHT 1864 also led to the accumulation of non-coated plasma membrane invaginations (Figure 7A,D), whereas the number of endosome-like vacuoles was not significantly altered (Figure 7A, E). Pharmacological blockade of Rac1 activity in neurons depleted of mDia1/3 further increased the number of plasma membrane invaginations significantly, while the additional effect on the number of endosome-like vacuoles remained insignificant (Figure 7F,G,H). mDia1/3 loss and Rac1 perturbation exhibited similar phenotypes with respect to the length and width of tubular membrane invaginations (Figure 7-Supplement 1C,D).

**Figure 7.**
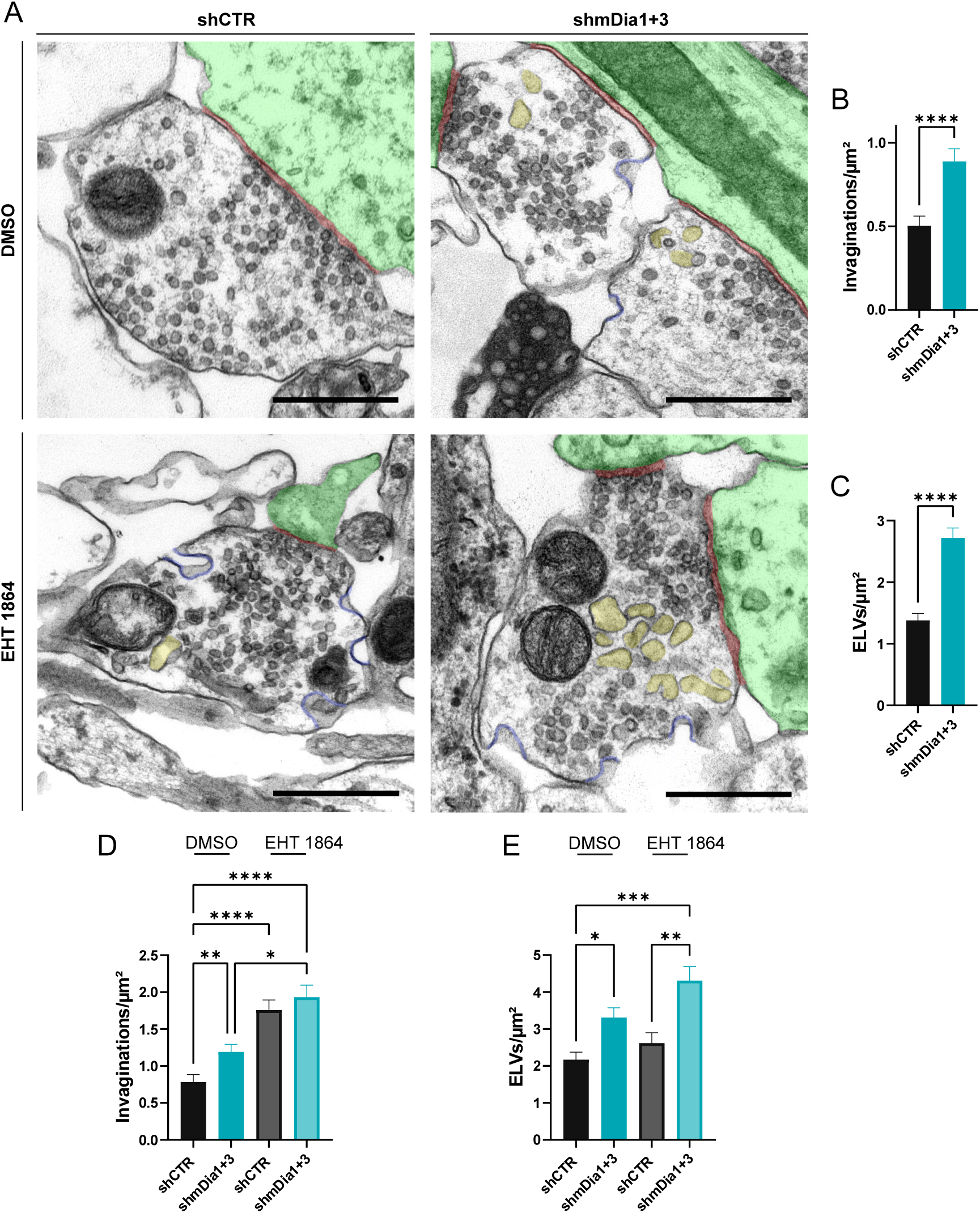
Defects in presynaptic ultrastructure induced by interference with mDia1/3-Rho and Rac1 signaling. **(A)** Representative synaptic electron micrographs from hippocampal neurons transduced with lentiviruses encoding shCTR or shmDia1+3 and treated with 0.1 % DMSO or 10 µM Rac1 Inhibitor (EHT 1864) for 2 h before fixation. Invaginations and ELVs are colored in blue and yellow, while postsynapse and synaptic cleft are colored in green and maroon, respectively. Scale bar, 250 nm. **(B)** Average number of invaginations per μm^2^ in shCTR (0.5 ± 0.1) and shmDia1+3 (0.9 ± 0.1; p < 0.0001, Mann-Whitney test) boutons. Data represent mean ± SEM. N = 3 independent experiments from n_shCTR_ = 326 synapses, n_shmDia1+3_ = 323 synapses. **(C)** Average number of ELVs per μm^2^ in shCTR (1.4 ± 0.1) and shmDia1+3 (2.7 ± 0.2; p < 0.0001, Mann-Whitney test) boutons. Data represent mean ± SEM. N = 3 independent experiments from n_shCTR_ = 326 synapses, n_shmDia1+3_ = 323 synapses. **(D)** Average number of invaginations per μm^2^ in shCTR and shmDia1+3 boutons treated with 0.1 % DMSO (0.8 ± 0.1 for shCTR; 1.2 ± 0.1 for shmDia1+3, p_shCTR + DMSO vs shmDia1+3 + DMSO_ < 0.01) or 10 µM EHT 1864 (1.8 ± 0.1 for shCTR, p_shCTR + DMSO vs shCTR + EHT 1864_ < 0.0001; 1.9 ± 0.2 for shmDia1+3, p_shCTR + DMSO vs shmDia1+3 + EHT 1864_ < 0.0001, p_shmDia1+3 + DMSO vs shmDia1+3 + EHT 1864_ < 0.05, Kruskal-Wallis test with Dunn’s post-test) for 2 h before fixation. Data represent mean ± SEM from n_shCTR + DMSO_ = 144 synapses, n_shmDia1+3 + DMSO_ = 143 synapses, n_shCTR + EHT 1864_ = 136 synapses, n_shmDia1+3 + EHT 1864_ = 153 synapses. **(E)** Average number of ELVs per μm^2^ in shCTR and shmDia1+3 boutons treated with 0.1 % DMSO (2.2 ± 0.2 for shCTR; 3.3 ± 0.3 for shmDia1+3, p_shCTR + DMSO vs shmDia1+3 + DMSO_ < 0.05) or 10 µM EHT 1864 (2.6 ± 0.3 for shCTR; 4.3 ± 0.4 for shmDia1+3, p_shCTR + DMSO vs shmDia1+3 + EHT 1864_ < 0.001, p_shCTR + EHT 1864 vs shmDia1+3 + EHT 1864_ < 0.01, Kruskal-Wallis test with Dunn’s post-test) for 2 h before fixation. Data represent mean ± SEM from n_shCTR + DMSO_ = 144 synapses, n_shmDia1+3 + DMSO_ = 143 synapses, n_shCTR + EHT 1864_ = 136 synapses, n_shmDia1+3 + EHT 1864_ = 153 synapses.

Collectively, these findings demonstrate that mDia1/3-Rho and Rac1 signaling pathways cooperatively act to facilitate presynaptic endocytosis and SV recycling.

## DISCUSSION

We found that loss of mDia1 either alone or in conjunction with the closely related mDia3 isoform slows the kinetics of SV endocytosis of exogenous pHluorin-tagged SV proteins and of endogenous vGAT without perturbing exocytic SV fusion. These phenotypes are accompanied by the accumulation of plasma membrane invaginations and vacuolar structures as well as an activity-dependent reduction of the total SV pool (Figures 1, 2, and 7). Furthrmore, we observe that interdependent mDia1/3-Rho and Rac1 signaling pathways cooperatively act to facilitate presynaptic endocytosis (Figures 5, 6 and 7). Several lines of evidence indicate that these endocytic phenotypes are a consequence of perturbed presynaptic actin levels or dynamics. (i) Endogenous presynaptic F-actin is reduced in mDia1/3-depleted hippocampal synapses (Figure 4) and (ii) actin polymerization-defective mDia1 fails to restore normal endocytosis kinetics in hippocampal neurons depleted of mDia1 (Figure 4). Conversely, (iii) pharmacological stabilization of F-actin by jasplakinolide (Figure 4) or (iv) expression of constitutively-active Rac1 to drive compensatory actin polymerization (for example via other formins and/ or ARP2/3) (Figure 6) rescues endocytosis in mDia1/3-KD hippocampal neurons. Moreover, (v) we find mDia1 (Figure 3) and F-actin (Figure 4) to accumulate at presynapses under conditions of impaired dynamin-dependent endocytosis. Together with the observation that inhibition of F-actin dynamics (Peng *et al*., 2011) in the combined presence of latrunculin A, jasplakinolide, and Y-27632 (Figure 1) slows the kinetics of SV endocytosis, our data support the hypothesis that presynaptic actin facilitates SV endocytosis downstream of mDia/ RhoA and Rac1-based signaling pathways.

Our data extend previous studies using conditional genetics that have identified an actin requirement for SV endocytosis (Wu *et al*., 2016). The endocytic phenotype described here for loss of mDia1/3, however, is substantially milder than that elicited by complete loss of β− or γ- actin, likely reflecting the fact that several partially redundant pathways cooperate to promote F-actin assembly at synapses (see below). In contrast to β- or γ-actin KO hippocampal neurons (Wu *et al*., 2016) we did not observe strong exocytic depression in response to train stimulation in mDia-depleted neurons (e.g. Figure 1-Supplement 1A-M) although we cannot rule out mild differences in synaptic strength and/ or release probability as observed in Rac1 KO hippocampal neurons and at the calyx of Held (Keine *et al*., 2022; O’Neil *et al*., 2021).

At the ultrastructural level, we find that genetic (KO) or lentiviral induced (KD) loss of mDia1 or mDia1/3 partially phenocopies acute perturbation of formin-mediated actin assembly (Soykan *et al*., 2017) with respect to the accumulation of plasma membrane invaginations. Unlike acute formin inhibition (Soykan *et al*., 2017), we find elevated numbers of post-fission endosome-like vacuoles in mDia-depleted hippocampal synapses. The latter correlate with a mild reduction of SV numbers, suggesting that mDia may have a further role in SV reformation downstream of the actual endocytosis reaction (Chanaday *et al*., 2019; Gan & Watanabe, 2018; Kononenko *et al*., 2014). The identified function of mDia1/3-mediated presynaptic actin assembly in presynaptic endocytosis and SV recycling is consistent with the actin binding properties of many endocytic proteins including Dynamin (Ferguson *et al*., 2009; Gu *et al*, 2014), PACSINs (Kessels & Qualmann, 2006), or Epsins, among others (McMahon & Boucrot, 2011; Merrifield *et al*., 2005; Saheki & De Camilli, 2012). How exactly actin functions to facilitate endocytosis of SV membranes remains to be fully understood. The comparably small size of the membrane invaginations that accumulate following pharmacological inhibition of formins (< 100 nm) or Rac1 or upon genetic loss of mDia1 (typically < 150 nm, although larger ones are observed) render an actomyosin-based constriction mechanism for SV endocytosis unlikely (consistent with (Sankaranarayanan *et al*., 2003)). We rather favor a -possibly indirect - function of F-actin in presynaptic endocytosis via formation of an F-actin ring that couples exocytic membrane compression to endocytic pit formation (Ogunmowo *et al*., 2023). Such a model is supported by the abundance of large plasma membrane invaginations distal from the active zone where SV fusion occurs at synapses following mDia1/3 and Rac1 loss of function (Figure 7). Consistent with this, it has been shown that mDia1 controls presynaptic membrane contractility (Deguchi *et al*, 2016), e.g. via the recently identified presynaptic actin corrals visualized in genome-engineered rat hippocampal neurons (Bingham *et al*, 2023).

A key result from our study is the finding that presynaptic endocytosis is regulated by interdependent, yet partially redundant signaling for F-actin assembly downstream of Rho and Rac1, e.g. pathways known to control linear and branched F-actin networks (Lawson & Ridley, 2018; Muller *et al*., 2020) that were shown to co-exist within hippocampal presynaptic boutons (Bingham *et al*., 2023). We demonstrate that loss of mDia1/3 or inhibition of Rho cause the hyperactivation of the actin-promoting GTPase Rac1 and that dual loss of mDia1/3 loss and Rac1 function results in synergistic inhibitory effects with respect to presynaptic endocytosis and ultrastructural defects (Figures 5-7). These findings suggest that the signaling network that regulates presynaptic actin assembly may have been evolutionary selected for plasticity and resilience. A resilient network for the control of actin dynamics at the presynapse may also help to explain the partially contradicting results from drug-based manipulations of actin and actin regulatory factors in different models of presynaptic neurotransmission reported in the past (Bleckert *et al*., 2012; Del Signore *et al*., 2021; Hori *et al*., 2022; Piriya Ananda Babu *et al*., 2020; Richards *et al*, 2004; Sankaranarayanan *et al*., 2003; Shupliakov *et al*., 2002; Wu & Chan, 2022; Wu *et al*., 2016). Moreover, accumulating evidence indicates that different pools of actin might be differentially amenable to drug treatments (Bleckert *et al*., 2012), e.g. as a consequence of the association of actin filaments with molecules such as tropomyosin (Gormal *et al*, 2017) that render them resistant to drugs, and/ or variations in the fraction of F-actin *vs*. monomeric actin pools (Higashida *et al*, 2008), providing a possible explanation for previously discrepant results.

A number of open questions remain. We predict that presynaptic actin assembly via mDia/ Rho and Rac1 facilitates all forms of presynaptic endocytosis that operate on timescales of milliseconds to seconds (Chanaday & Kavalali, 2018; Clayton & Cousin, 2009; Watanabe & Boucrot, 2017) and at many types of synapses (e.g. small central synapses, ribbon synapses, the neuromuscular junction), but further studies will be needed to test this hypothesis. We note that fast endophilin-mediated endocytosis (FEME), a process with resemblance to endocytosis at synapses (Watanabe & Boucrot, 2017), has been shown to be differentially regulated by actin and Rho GTPase family members (Boucrot *et al*, 2015; Renard *et al*, 2015). At present we do not understand if and how RhoA/B and Rac1 GTPase activation is nested into the SV cycle and whether these GTPases operate at the same or different nanoscale sites and on the same or different membranes. Future studies will be required to tackle these questions.

## MATERIALS AND METHODS

### Key Resources Table

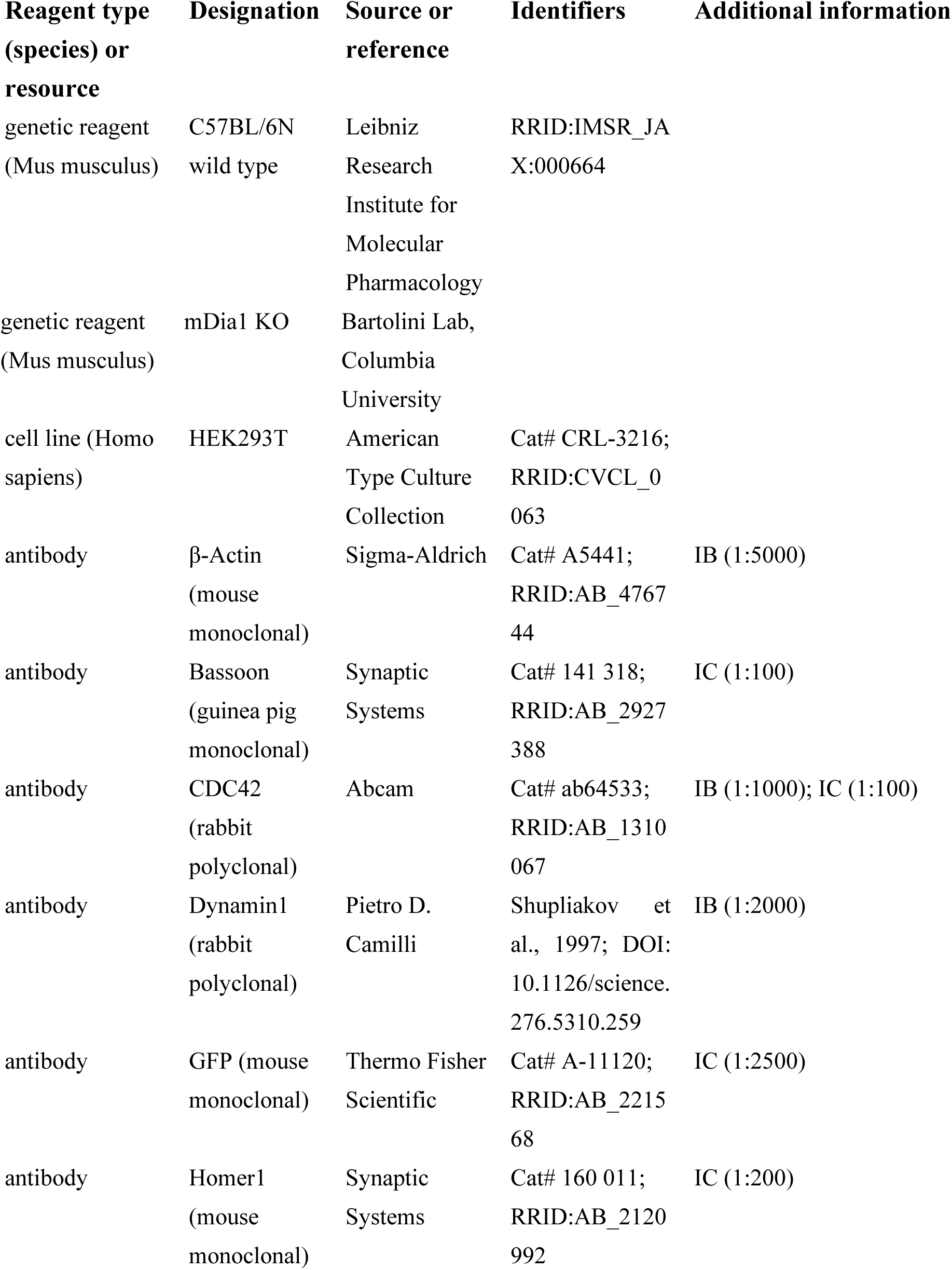
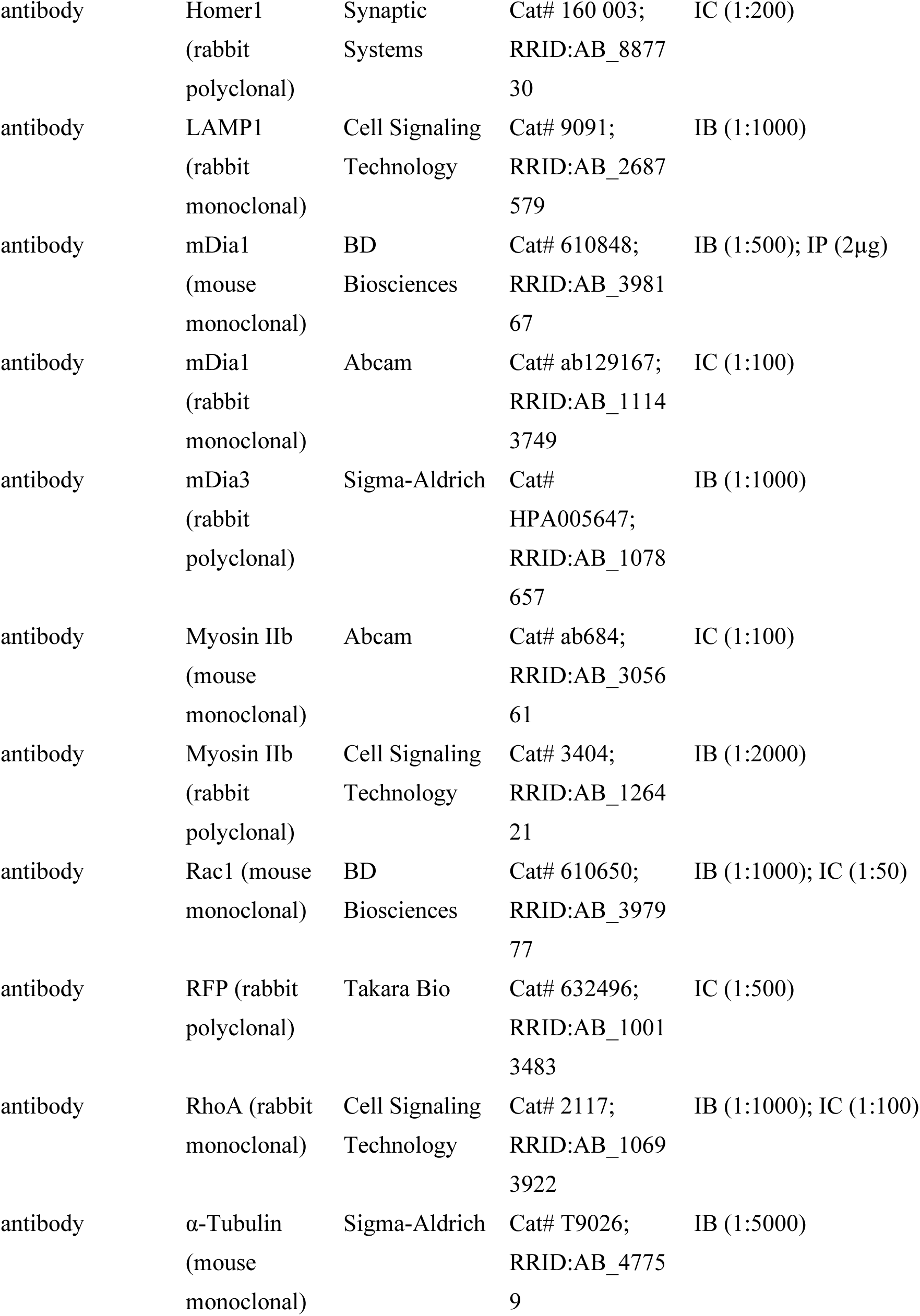
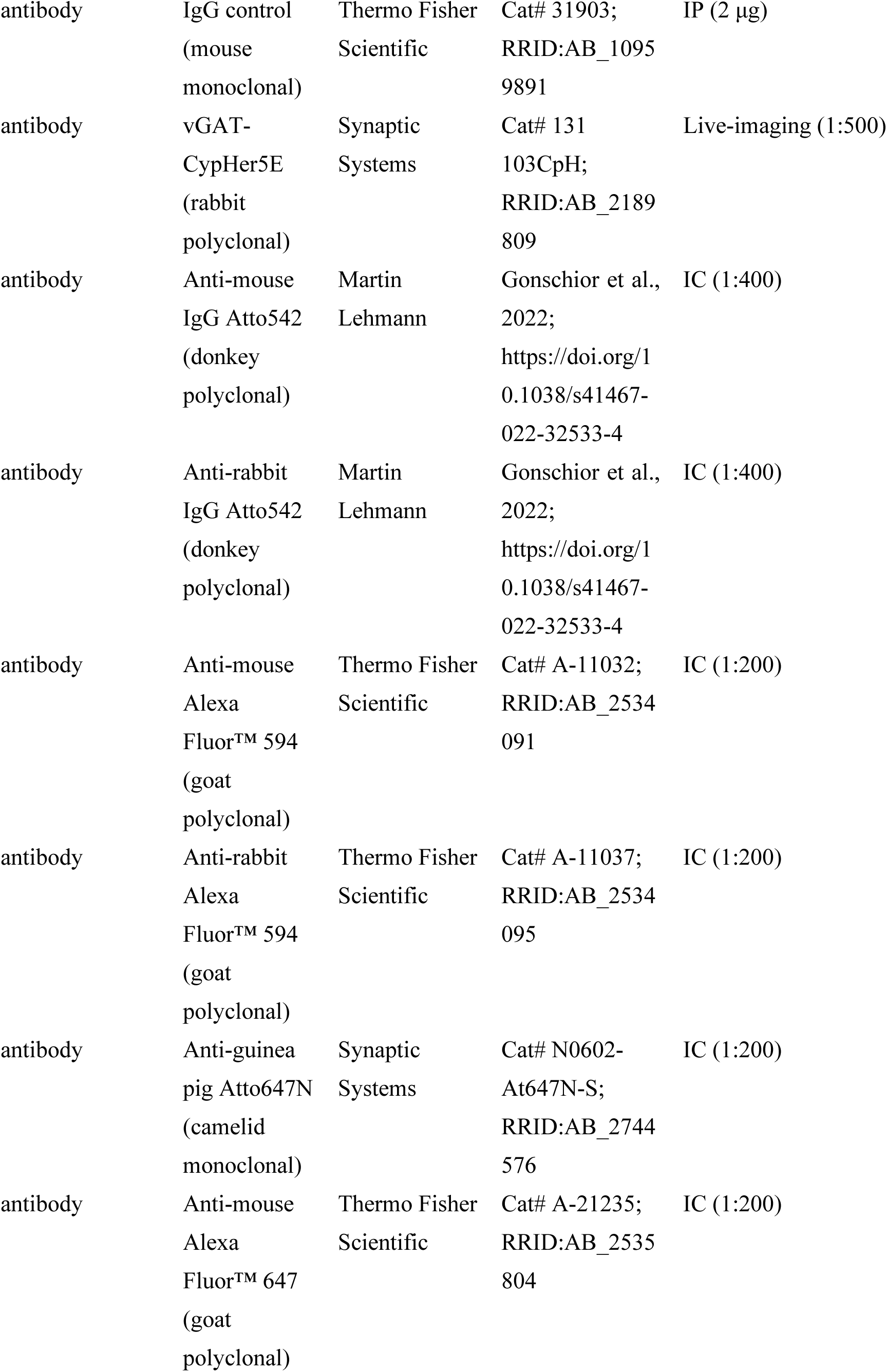
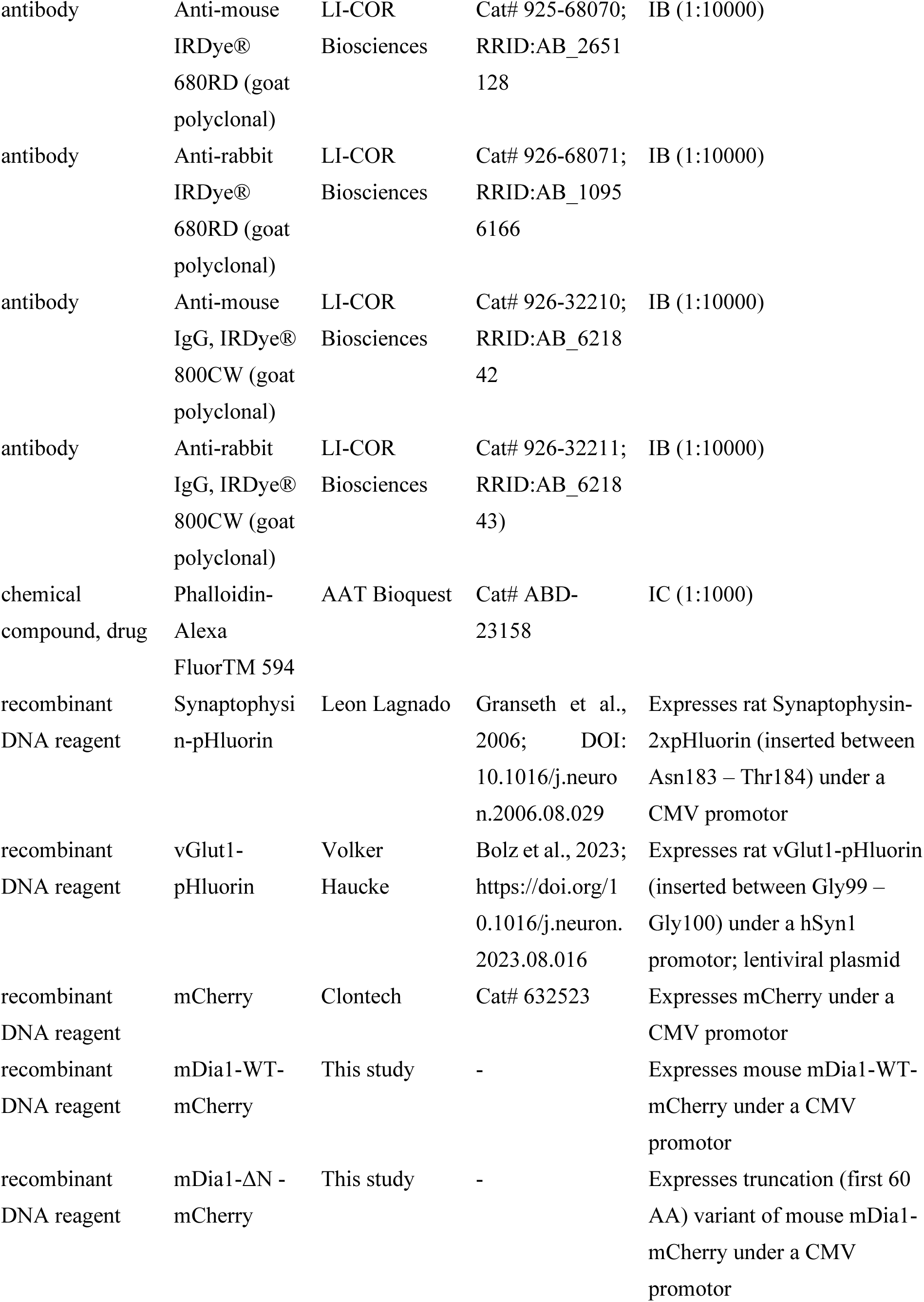
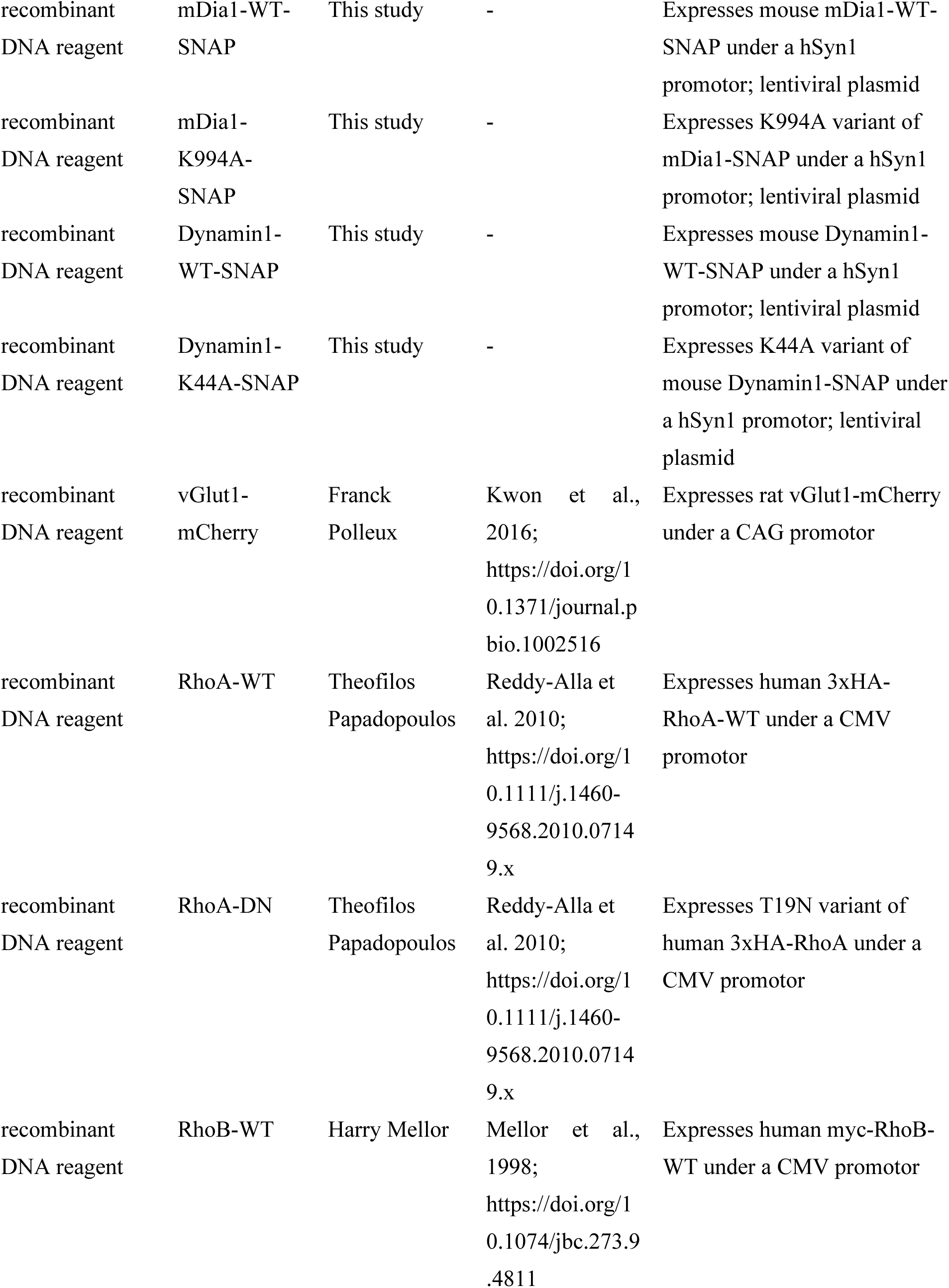
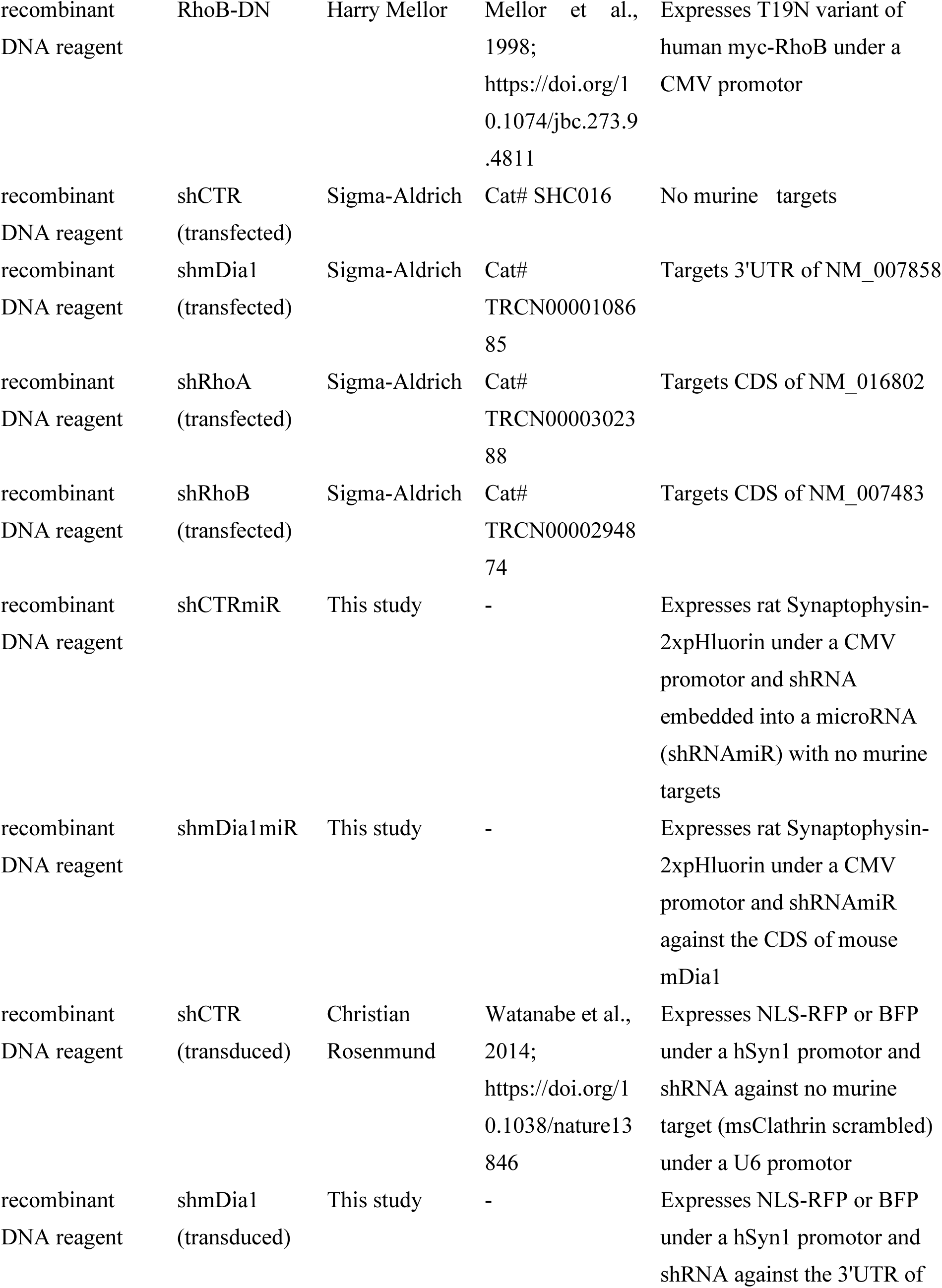
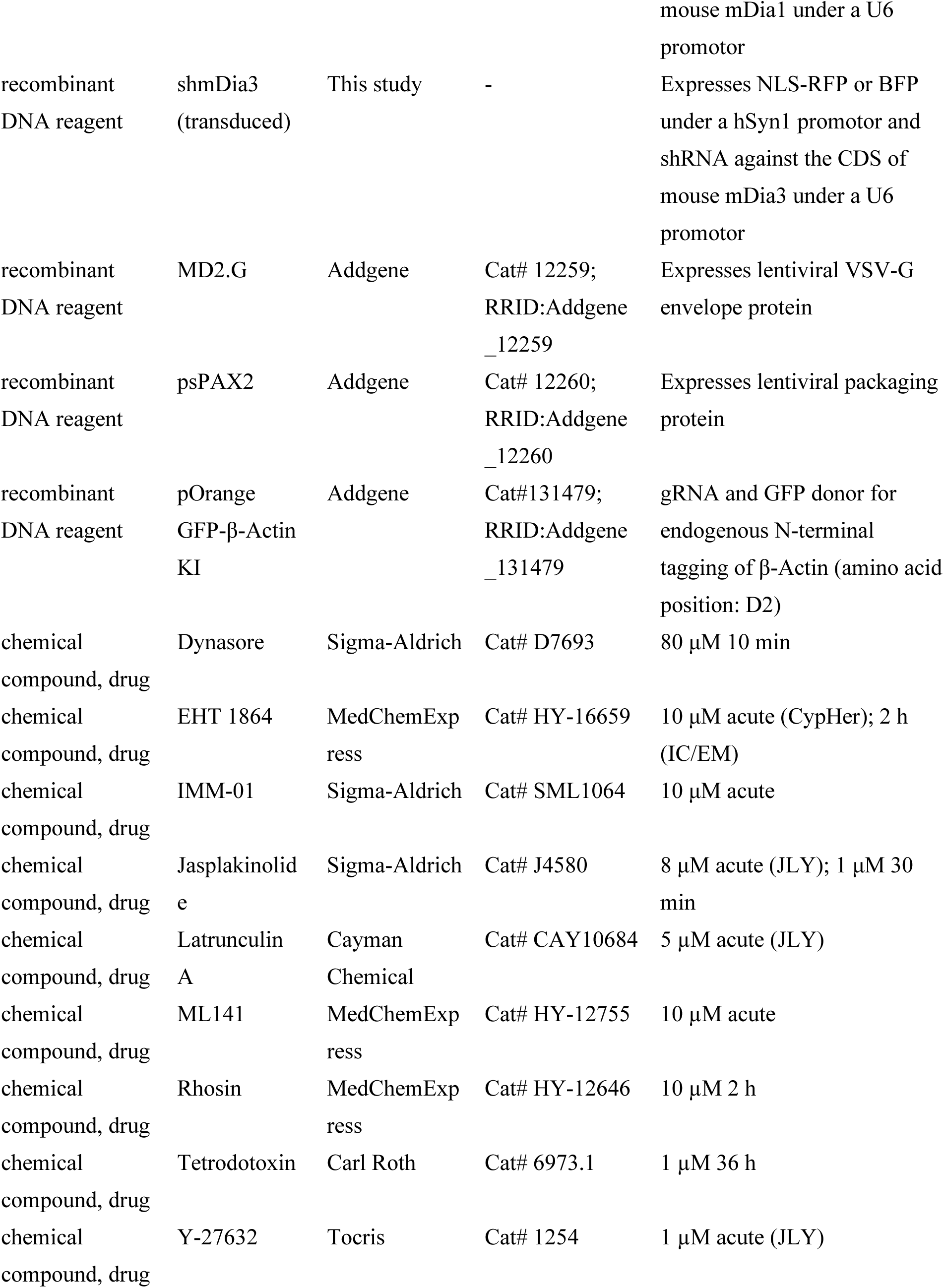
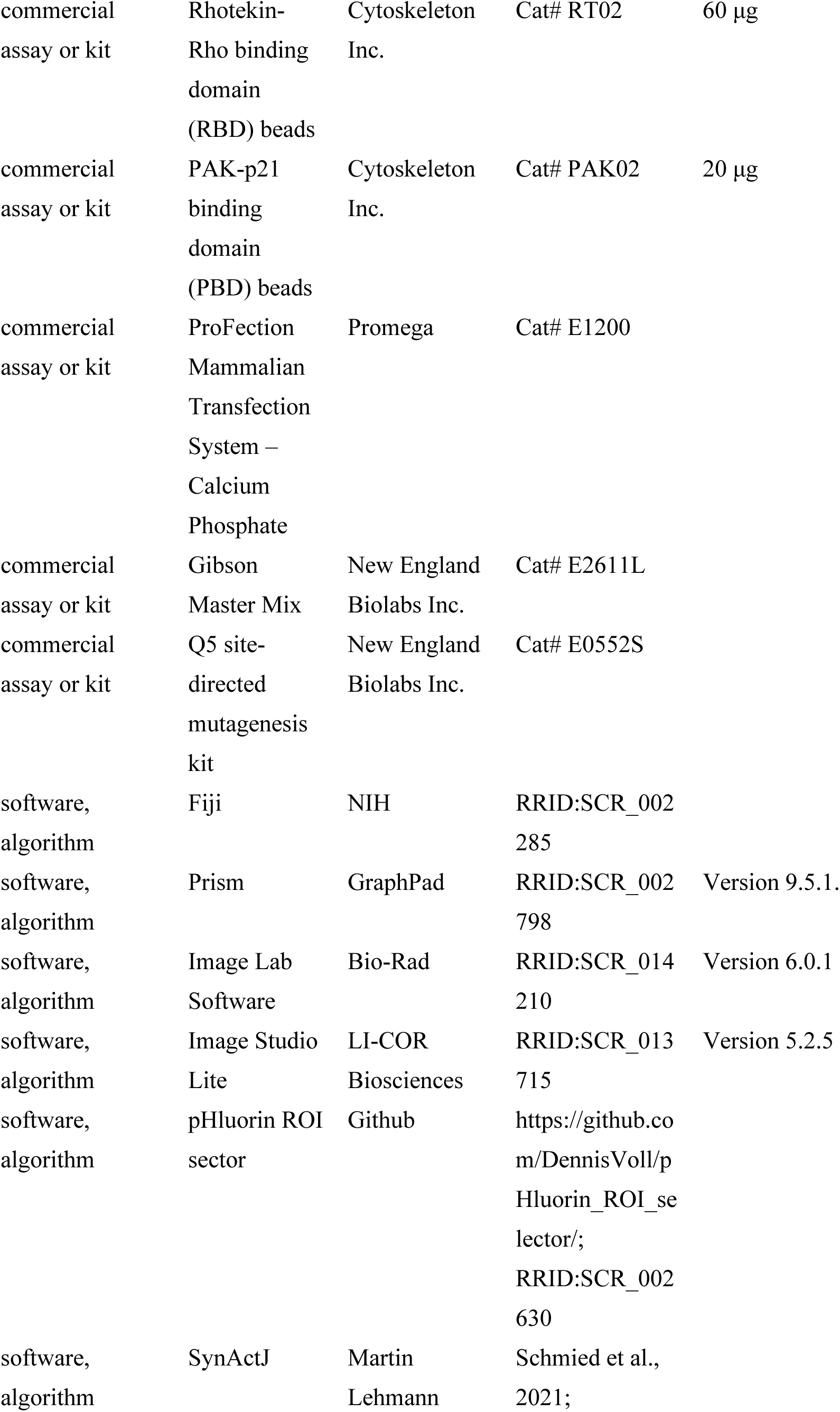
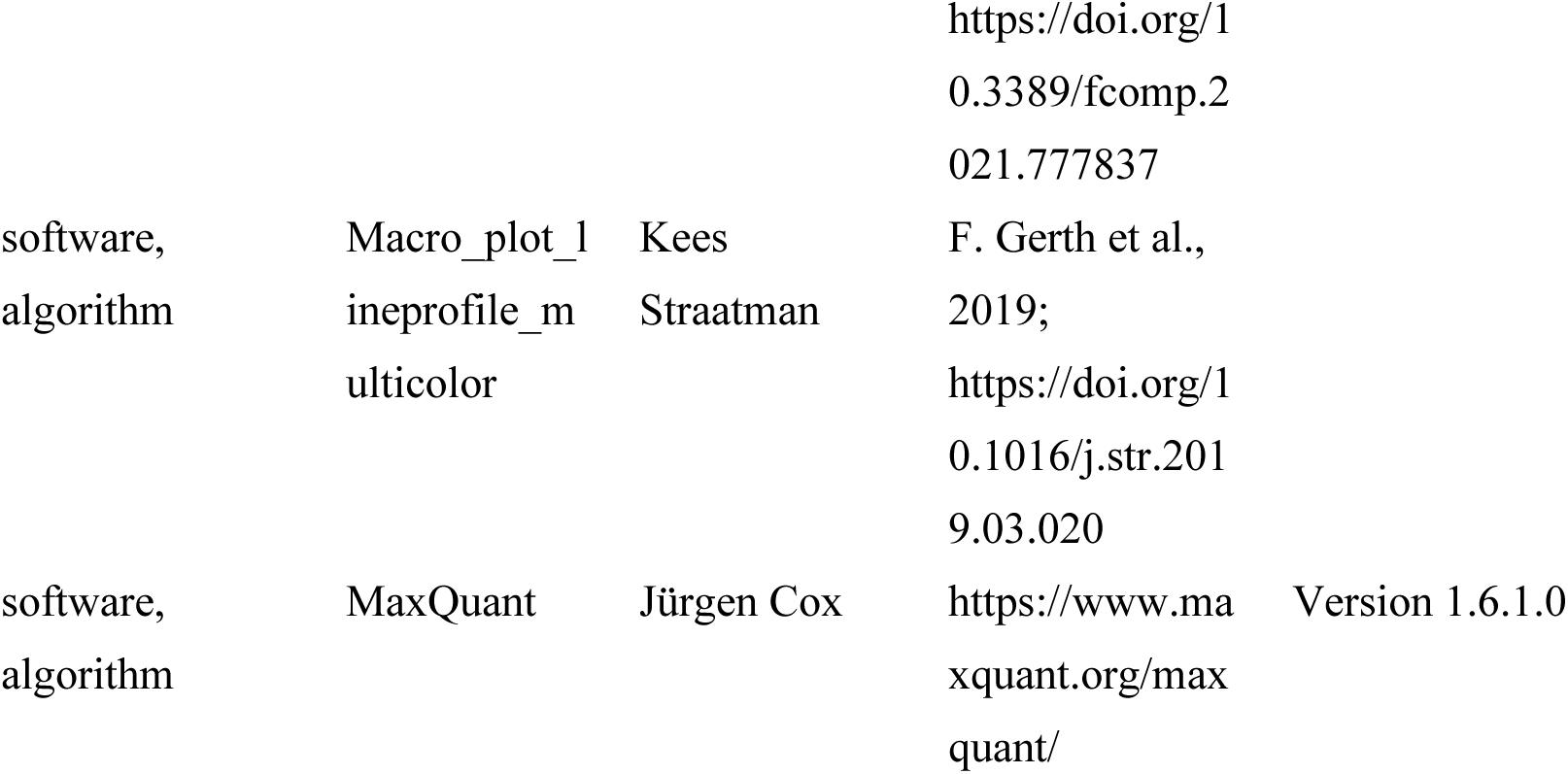

## Materials

### Animals

Primary neurons were obtained from either wild-type C57BL/6J (Charles River, RRID: IMSR_JAX:000664) or mDia1 KO mice (Peng *et al*., 2007; Peng *et al*., 2003). All animal experiments were reviewed and approved by the ethics committee of the *Landesamt für Gesundheit und Soziales* (LAGeSo) Berlin or the Committee on the Ethics of Animal Experiments of Columbia University and conducted according to the committees’ guidelines (LAGeSo) or the Guide for the Care and Use of Laboratory Animals of the National Institutes of Health (for mDia1 KO mice). At the facilities, animal care officers monitored compliance with all regulations. Mice were group-housed under 12/12 h light/dark cycle with access to food and water *ad libitum*. Mice from both genders were used and cultures were randomly allocated to experimental groups (e.g., different treatments). Multiple independent experiments using several biological replicates were carried out as indicated in the Figure legends.

### Antibodies

Antibodies and their working dilutions used in this study are denoted in the Key Resource Table (IB: Immunoblot; IC: Immunocytochemistry; IP: Immunoprecipitation). Antibodies were stored according to the manufacturer’s recommendations. All secondary antibodies are species-specific (highly cross-adsorbed).

### Cell lines

Human embryonic kidney 293T (HEK293T) cells were obtained from the American Type Culture Collection (Cat# CRL-3216; RRID:CVCL_0063). Cells were cultured in Dulbecco’s modified Eagle’s medium supplemented with glucose (DMEM; 4.5 g/L; Thermo Fisher Scientific) and 10% heat-inactivated fetal bovine serum (FBS; Gibco), penicillin (100 U/ml; Gibco), and streptomycin (100 μg/ml; Gibco) at 37°C and 5% CO_2_. Cells were routinely tested for mycoplasma contamination.

### Chemicals

Compounds were dissolved in dimethyl sulfoxide (DMSO), unless indicated otherwise, and diluted 1: 1000 to their working concentrations (s. Key Resource Table). For acute pharmacological treatment (in pHluorin/CypHer assays), drugs were added to the imaging buffer. For longer incubations, the conditioned cell media was supplemented with the chemicals (s. Key Resource Table for incubation time).

For silencing neuronal network activity, sodium channels were inhibited by addition of tetrodotoxin (TTX; in 10 mM sodium acetate, pH 5.3) to the neuronal culture medium at day in vitro (DIV) 12 for 36 h. As a control, cells from the same preparation were treated with equal volumes of 10 mM sodium acetate (annotated as Vehicle).

### Plasmids

All recombinant DNA reagents used for protein expression are listed in the Key Resource Table: Synaptophysin-pHluorin was a kind gift from Dr. L. Lagnado (Univ. of Sussex, UK), vGLUT1-pHluorin was generated in-house by Svenja Bolz as previously described, while vGLUT1-mCherry was kindly provided by Dr. Franck Polleux (New York, USA). HA-tagged RhoA-WT and RhoA-T19N were kind gifts from Dr. Theofilos Papadopoulos (Göttingen, Germany) and myc-tagged RhoB-WT and RhoB-T19N were kindly gifted by Dr. Harry Mellor (Bristol, UK). All other expression vectors were generated for this study. Plasmids based on pEGFP-C1, pmCherry-N1, pCAG and pcDNA3 utilising a CMV or CAG promotor (s. Key Resource table) were used for overexpression based on transfection of neuronal cells, while pFUGW vectors carrying a human Synapsin1 promotor (hSyn1) were used for the generation of lentiviral particles for the transduction of neurons (s. Key Resource Table annotated as “lentiviral plasmid”). In the course of this study several approaches to deplete mDia1 have been carried out and are annotated in the Figure legends: For knockdown of mDia1, RhoA and RhoB by transfection, commercially available lentiviral small hairpin RNA (shRNA) vectors based on the pLKO.1 backbone (s. Key Resource Table; annotated as “transfected”) were purchased from Sigma-Aldrich. To reduce the amount of DNA needed for transfection to perform pHluorin assays, vectors expressing shRNA embedded into a microRNA (miRNA) context for mDia1 together with Synaptophysin-2x-pHluorin as a reporter were cloned based on pRRLsinPPT-emGFP-miR Control, a kind gift of Dr. Peter S. McPherson (Montreal, Canada). Finally, lentiviral vectors for knockdown of mDia1 and mDia3 (annotated in Key Resource Table as “transduced”) were generated based on the backbone f(U6) sNLS-RFPw shCTR, a kind gift from Prof. Christian Rosenmund (Berlin, Germany). All vectors used for genetic depletion via RNA interference are listed in the Key Resource Table and express gene-specific shRNA under a U6 promotor, that either targets the coding sequence (CDS) or the 3’-untranslated region (3’-UTR).

### Oligonucleotides

Plasmids generated for this study were cloned using oligonucleotides (BioTez GmbH, Berlin, Germany) listed in Supporting Table 1 (lower case denotes nucleotides that do not anneal to the backbone; underlined nucleotides denote sense and antisense sequences of shRNA).

## Methods

### Generation of expression plasmids

Expression plasmids generated for this study were cloned by PCR amplification (Phusion^TM^ High-Fidelity DNA polymerase) and restriction enzyme (Thermo Fisher Scientific; Fast digest) digest according to the manufacturers’ manual. pSNAP-N1 was generated in-house by Hannes Gonschior by sub-cloning pSNAP_f_ (New England BioLabs Inc., Cat#N9183S) via PCR and the restriction enzymes *AgeI* and *NotI* into pmCherry-N1. pFUG_hSyn_MCS was cloned by Amirreza Ohadi to enable simple insertion of SNAP-tagged proteins by inserting a multiple cloning site (MCS) on annealed oligonucleotides into a pFUG_hSyn1 backbone (a gift from Christian Rosenmund, Berlin, Germany) cut by *AgeI* and *EcoRI*. To generate mDia1-WT-mCherry, the coding sequence of mDia1 was cut from mDia1-mEmerald-N1 (Addgene, Cat#54157) and pasted into pmCherry-N1 by *AgeI* and *XhoI* digestion. mDia1-WT-SNAP was cloned by cutting out the coding sequence of mDia1 from mDia1-mEmerald-N1 and pasting it into pSNAP-N1 by *AgeI* and *NheI* digest of both vector and insert. To subclone mDia1-SNAP into a lentiviral vector the coding sequence of mDia1-SNAP was pasted from mDia1-WT-SNAP-N1 into pFUG_hSyn1_MCS utilizing common cut sites for *NheI* and *NotI* digest. For the generation of a lentiviral vector expressing Dynamin-WT, the coding sequence of Dynamin1 was first extracted from Dynamin1-pmCherry-N1 (Addgene; Cat#27697) by *EcoRI* and *XmaI* digestion and cloned into pSNAP-N1 with the same enzymes. Subsequently, Dynamin1-SNAP was isolated by *NheI* and *NotI* digest and transformed into pFUG_hSyn1_MCS. The mDia1- ΔN truncation variant was generated by introducing a new start codon on a primer shifted by 60 amino acids (s. Supporting Table 1), PCR amplifying the truncated DNA and subcloning the template into pmCherry-N1 by *BglII* and *AgeI* digest. For introducing point mutations in mDia1 (Lysine-994 to Alanine; K994A) and Dynamin1 (Lysine-44 to Alanine; K44A), the Q5 site-directed mutagenesis kit (New England Biolabs Inc.; E0552S) was used according to the manufacturer’s manual and oligonucleotides listed in Supporting Table 1.

All vectors were confirmed by Sanger sequencing (LGC Genomics, Berlin, Germany) and amplified by self-made chemically competent TOP10 *E.coli* before purifying respective endotoxin-free DNA by 2-propanole precipitation.

### Isolation, culture and transfection of primary hippocampal neurons

Neuronal hippocampal cell cultures were prepared as described before (Lopez-Hernandez *et al*., 2022; Soykan *et al*., 2017). In short, hippocampi from postnatal mice (p0-p3) were surgically isolated and dissociated into single cells by trypsin (5 g/L, Sigma-Aldrich) digestion. Neurons (100000 cells/ well of a 6-well) were plated onto poly-L-lysine-coated coverslips and grown in modified Eagle medium (MEM; Thermo Fisher) supplemented with 5% FCS and 2% B-27 (Gibco) and maintained at 5% CO_2_ and 37°C in humidified incubators. In addition, 2 μM cytosine β-D-arabinofuranoside (AraC) was added to the cell culture media in the first 2 days in vitro (DIV) to limit glial proliferation. For transient protein expression, neurons were transfected on DIV 7-9 utilizing a Calcium phosphate transfection kit (Promega; Cat# E1200): In brief, 1-6 μg plasmid DNA (per well of a 6-well plate) were mixed with 250 mM calcium phosphate (CaCl_2_) in ultrapure nuclease-free water. The resulting solution was added to equal volumes of 2x 4-(2-hydroxyethyl)- 1-piperazineethanesulfonic acid buffered saline (2x HEPES; 100 µL) and incubated at room temperature for 20 min. Resulting precipitates were added dropwise to cells that had been transferred to osmolarity-adjusted Neurobasal-A (NBA; Gibco) media to induce starvation. After incubation at 37°C and 5% CO_2_ for 30 min, neurons were washed thrice with osmolarity-adjusted Hank’s balanced salt solutions (HBSS; Gibco) and transferred back to their original conditioned media.

### shRNA cloning and lentivirus production

Knockdown of proteins was achieved through RNA interference either by CaCl_2_ transfection of gene-specific shRNA encoding vectors (pLKO.1; Sigma-Aldrich or shRNAmiR) on DIV 7 or by transduction of cells with lentiviral particles harboring the gene-specific shRNA on DIV 2. The respective method is indicated in the Figure legends as *transfected* or *transduced*. To reduce the amount of DNA needed for transfection and to improve neuronal health, shRNAmiR were expressed from the 3’UTR of Synaptophysin-pHluorin based on (Ritter *et al*, 2017): The reporter protein/synthetic cassette was generated by PCR amplifying Synaptophysin-2x-pHluorin with oligos (s. Supporting Table 1) harboring *XbaI* and *SalI* sites. The eGFP in the RRLsinPPT-eGFP-miRCTR plasmid (a kind gift from Dr. Peter S. McPherson, McGill University, Canada) was replaced by the PCR product through similar restriction digest to yield shCTRmiR. miRmDia1 (start position 4892, targeting sequence matches open-reading frame) was designed with BLOCK-iT^TM^ (Thermo Fisher Scientific) and subcloned into shCTRmiR to yield shmDia1miR following protocol in (Ritter *et al*., 2017).

As both transfection strategies were limited by low efficiency, lentiviral knockdown was carried out: Lentiviral particles were based on a shuttle vector (pFUGw) driving the expression of a nuclear-targeted red fluorescent protein (NLS-RFP) under a human Synapsin1 (hSyn1) promotor to monitor infection efficiency in neurons and a scrambled mouse shRNA against Clathrin without any murine targets, which was used as the control virus (f(U6) sNLS-RFPw shCTR). To prevent crosstalk between the NLS-RFP and other mCherry constructs, a similar backbone expressing the blue fluorescent protein (BFP) as a reporter was generated in house by Klaas Ypermann: The backbone was digested by *XbaI* and *PacI*, the hSyn1 promotor and mTagBFP (Addgene; #Cat 105772) were amplified with oligos (s. Supporting Table 1), gel extracted and assembled utilizing a Gibson master mix (New England BioLabs Inc.; Cat#E2611L) to yield f(U6) BFP shCTR. To knockdown mDia1, a shRNA sequence based on 5’-GCCTAAATGGTCAAGGAGATA-3’ as the sense nucleotide corresponding to the 3’UTR of mouse mDia1 (NM_007858.4; Sigma-Aldrich, Cat# TRCN0000108685) was designed as an oligonucleotide with overhangs (s. Supporting Table 1) and annealed into the f(U6) sNLS-RFPw shCTR backbone cut with *BamHI* and *PacI* to yield the vector f(U6) sNLS-RFPw/BFP shmDia1. For mDia3 shRNA, a sense sequence based on 5’-GCCCTAATCCAGAATCTTGTA-3’ corresponding to nucleotides 2473-2493 of mouse mDia3 (DIAPH2; NM_172493.2; Sigma-Aldrich, Cat# TRCN0000108782) was used to construct f(U6) sNLS-RFPw/BFP shmDia3 as described above. Production of lentiviral particles was conducted using the second-generation packaging system: In brief, HEK293T were co-transfected with lentiviral shRNA constructs and the packaging plasmids psPAX2 (Addgene; Cat# 12260) and MD2.G (Addgene; Cat# 12259) using CaCl_2_. After 12 h, the cell media was replaced. Virus-containing supernatants were collected at 48 and 72 h after transfection, filtered to remove cell debris and particles were concentrated (30-fold) via low-speed centrifugation (Amicon Ultra-15, Ultracel-100; Merck Millipore; Cat# UFC9100) before aliquoting and storage at −70°C. For all experiments, an infection rate of over 95% was achieved at DIV 14-16.

## Microscopy

### Live-imaging of SV recycling

For live-imaging of Synaptophysin and vesicular glutamate transporter 1 (vGLUT1) recycling, SV proteins fused to the green-fluorescent protein-based pH-sensitive fluorescent reporter pHluorin at their luminal domain were overexpressed by plasmid transfection (Synaptophysin-pHluorin; DIV 7) or lentiviral transduction (vGLUT1-pHluorin; DIV 2).

For following endogenous vesicular gamma-aminobutyric acid transporter (vGAT) recycling, spontaneously active synaptic boutons were labeled by incubating cells with CypHer5E-conjugated antibodies against the luminal domain of vGAT (1:500 from a 1 mg/ml stock; Synaptic Systems; Cat#131,103CpH) for 2 h in their respective conditioned culture media at 37 °C and 5 % CO_2_ prior to imaging.

To investigate kinetics of SV recycling, neurons at DIV 14-16 were placed into an RC-47FSLP stimulation chamber (Warner Instruments) in osmolarity-adjusted imaging buffer [170 mM sodium chloride (NaCl), 20 mM N-Tris(hydroxyl-methyl)-methyl-2-aminoethane-sulphonic acid (TES), 5 mM Glucose, 5 mM sodium hydrogencabonate (NaHCO_3_), 3.5 mM potassium chloride (KCl), 1.3 mM CaCl_2_, 1.2 mM sodium sulfate (Na_2_SO_4_), 1.2 mM magnesium chloride (MgCl_2_), 0.4 mM potassium dihydrogenphosphate (KH_2_PO_4_), 50 μM *(2R)-*amino-5-phosphonovaleric acid (AP5) and 10 μM 6-cyano-7-nitroquinoxaline-2,3-dione (CNQX); pH 7.4] at 37°C (Tempcontrol 37-2 digital). Cells were subjected to electrical field-stimulation (MultiStim System-D330; Digimeter Ltd.) with annotated stimulation trains to evoke action potentials (APs) [40 Hz, 5s (200 APs), 40 Hz, 2s (80 APs), 20 Hz, 2s (40 APs); at 100 mA]. Following changes in fluorescence were tracked by an inverted Zeiss Axiovert 200M microscope, equipped with a 40x oil-immersion EC Plan Neofluar objective (NA 1.30), an EM-CCD camera (Evolve Delta 512) and a pE-300^white^ LED light source (CoolLED). The scanning format was set to 512 x 512 pixels with 16-bit sampling. eGFP (Excitation: BP470-40; Emission: BP535-50; Zeiss filter set 38) or Cy5 (Excitation: BP640-30; Emission: BP525-50; Zeiss filter set 50) filter sets were used for pHluorin or CypHer assays, respectively. Images were acquired at 0.5 Hz frame rate for 100 s with 50 (Syph-pHluorin) or 100 (vGLUT1-pH, vGAT-CypHer) ms exposure with an electron multiplying gain of 250 operated through Fiji-based MicroManager 4.11 software.

Analysis of responding boutons was performed through custom written macros to identify regions of interest (ROIs) (https://github.com/DennisVoll/pHluorin_ROI_selector/) in an automated manner using SynActJ (Schmied *et al*, 2021). Such analysis averaged fluorescence for each time point in an image series (video) of at least > 20 responding boutons and corrected values for background fluorescence yielding raw background-corrected fluorescence (F). The fold increase of fluorescence after stimulation (F_max_) can serve as a measure for exocytic fusion and is calculated by normalising F by the mean intensity of F before stimulation (Baseline/basal fluorescence (F_0_) = mean intensity of first 5 frames for pHluorin; first 10 frames for CypHer) for each time point (surface normalisation = F/F_0_ for pHluorin; F_0_ – F_min_ for CypHer). To account for boutons with varying pHluorin expression in one image series and to compare reacidification kinetics, the fluorescence of each time point was subtracted by the basal fluorescence (F_0_) to yield ΔF (F-F_0_), which was then normalised by its peak value (ΔF_max_). Resulting peak normalised curves (ΔF/ ΔF_max_) are annotated as norm. ΔF. Endocytic decay constants (τ) were calculated by averaging and then fitting the norm. ΔF (ΔF/ΔF_max_) traces of all videos in one condition (N) to a mono-exponential decay curve [y_0_ + A*e ^(-t/τ)^] with the constraints of y_0_ = 1 and offset = 0 in Prism 9 (Graphpad).

CypHer-traces had to be corrected for photobleaching: The decay constant of the bleaching curve was determined by fitting the data points of the first 10 frames (20 s, prior to stimulation) to a mono-exponential decay curve. The corresponding value of the photobleaching curve at a given time *t* was added to the raw fluorescence intensity measured at each corresponding time point to correct for the loss of intensity due to bleaching.

### Immunocytochemistry

For immunostainings, neuronal cultures were chemically fixed on DIV 14-16 using 4 % p-formaldehyde (PFA) and 4 % sucrose in phosphate buffered saline (PBS) for 15 min at room temperature (RT). For experiments in Figure 3C, G-I, cultures were stimulated with a 40 Hz train for 5 s in imaging buffer before immediate fixation. After fixation, cells were washed thrice with PBS and incubated with permeabilization buffer (10% normal goat serum, 0.3 % Triton X-100 in PBS) for 30 min at RT, followed by primary antibody incubation of proteins of interest in permeabilization buffer at indicated dilutions (see Key Resource Table) at 4°C overnight. Subsequently, unbound antibodies were removed by three PBS washes while bound antibodies were decorated by corresponding fluorophore-coupled secondary antibodies in permeabilization buffer for 1h at RT. For F-Actin staining, Phalloidin-Alexa Fluor^TM^ 594 (1:1000 stock; AAT Bioquest; Cat# ABD-23158) was added in the secondary incubation step. Finally, neurons were washed thrice with PBS, followed by two washes with ultra-pure water. Coverslips were dried for 2 h before mounting in ProLong^TM^ Gold Antifade (Thermo Fisher Scientific; #P36934) on glass slides (Thermo Fisher Scientific; VWR; Cat#630-1985). The slides were cured for at least 72 h at RT before imaging.

### Multicolor time-gated STED imaging

STED images were acquired from fixed samples by a HC PL APO CS2 100 x oil objective (1.40 NA) on a Leica SP8 TCS STED 3x microscope (Leica Microsystems) equipped with a pulsed white-light excitation laser (WLL; ∼ 80 ps pulse width, 80 MHz repetition rate; NKT Photonics) and a STED laser for depletion (775 nm). The scanning format was set to 1024 x 1024 pixels, with 8-bit sampling, 4x line averaging, 4x frame accumulation and 6 x optical zoom, yielding a final pixel size of 18.9 nm. Three-color imaging was performed by sequentially exciting following fluorophores (excitation filter = Exf; emission filter = Emf): Atto647N (Exf: 640 nm; Emf: 650-700 nm); Alexa Fluor^TM^ 594 (Exf: 590 nm; Emf: 600-640 nm) and Atto542 (Exf: 540 nm; Emf: 550-580 nm) operated by the Leica Application Suite X (Leica Microsystems, 2020). For stimulated depletion of the signal, the 775 nm STED laser was applied to all emissions. Detection of the resulting signal was time-gated by 0.3-6 ns to allow enough time for stimulated depletion and collected by two sensitive HyD detectors at appropriate spectral regions distinct from the STED laser wavelength. Settings in independent experiments were similar between conditions to allow quantification of signals. Raw data obtained from three-channel time-gated STED (gSTED) imaging were analyzed with Fiji. For analysis, only synapses oriented in the xy-plane exhibiting a clear separation of Bassoon (presynapse) and Homer1 (postsynapse) clusters were taken into account. To determine synaptic localisation and presynaptic levels of proteins of interest, multicolor line profiles were measured: A line (1.0 μm length, 0.4 μm width) perpendicular to the synaptic cleft (space between Bassoon & Homer1 cluster) was drawn (Gerth *et al*., 2017) and fluorescence intensity of all three channels along this line was measured using a Fiji Macro (Macro_plot_lineprofile_multicolor; Dr. Kees Straatman, University of Leicester, UK). Resulting profiles were aligned to the maximum Bassoon intensity, which was set to 0 nm. For localization analysis, all three profiles were normalized to their maxima, which were set to 1. For quantification of presynaptic protein levels, only the fractions of the non-normalized line profiles of proteins of interest overlapping with the normalized averaged Bassoon distribution (between 151.4 and −37.8 nm; Figure 3-Supplement 1H) were integrated. Intensities were normalized to controls (DMSO, shCTR, WT) which were set to 100.

### Imaging of endogenous β-Actin levels

To visualize endogenous β-Actin levels, eGFP knockin following the ORANGE method was performed (Willems *et al*., 2020): Neurons were transfected with vGLUT1-mCherry (Franck Polleux) and pOrange GFP-β-Actin KI (Addgene; Cat#131479) by CaCl_2_ on DIV 2. Directly following transfection, cells were transduced with shCTR or shmDia1+3 lentiviral particles carrying BFP as a reporter to prevent crosstalk between mCherry and NLS-RFP expression. Neuronal cultures were chemically fixed on DIV 15. After fixation, endogenous GFP and overexpressed vGLUT1-mCherry intensities were enhanced by additional antibody (anti-GFP and anti-RFP; see Key Resource Table) incubation according to the immunocytochemistry protocol described above. Images of neurons were taken on the STED microscope with and without stimulated depletion (confocal) with the same settings as indicated above. For analysis, confocal images were filtered by Gaussian blur (sigma = 2 pixel) and auto-thresholded using Otsu’s method. Whenever confocal vGLUT1 signal co-localized with confocal Actin clusters, the area was chosen as a ROI. Actin intensity was then measured in the ROI applied to the STED image. The intensities of β-Actin were normalized to shCTR which was set to 100.

### Transmission electron microscopy

To investigate synaptic effects of F-actin manipulations at the ultrastructural level, cells on coverslips were chemically fixed with 2 % glutaraldehyde in cacodylat buffer (CDB; 0.1 M sodium cacodylat) for 1 h at RT. Coverslips were washed thrice with CDB before osmification by 1 % (w/v) osmium tetroxide (OsO_4_) and 1.5 % (w/v) potassium hexacyanoferrat (K_3_Fe(CN)_6_) in CBD for 1 h at 4°C. After postfixation, cells were stained with 1 % (w/v) uranyl acetate, dehydrated by methanol gradients and finally embedded by epoxy resin (Sigma Aldrich; Cat# 45359) infiltration. After polymerization (60°C, 30 h), coverslips were removed and ultra-thin (70 nm) sections were cut and contrasted with 2 % (w/v) uranyl acetate and 80 mM lead citrate. 8-bit Images were obtained on a Zeiss 900 transmission electron microscope equipped with Olympus MegaViewIII or Olympus Morada G2 digital cameras at 30000 x magnification yielding a pixel size of 1.07 nm. Subsequently, morphometry (density of synaptic vesicles, endosome-like vacuoles, clathrin-coated vesicles, clathrin-coated pits and non-coated invaginations) was analyzed from synaptic profiles with clearly distinguishable active zones and adjacent synaptic vesicles in a blinded manner.

## Biochemistry

### Immunoblotting

To compare protein levels between experimental conditions, immunoblotting was performed: Protein concentrations in lysates were measured by BCA assay (Thermo Fisher Scientific; Cat#23227) and equal protein amounts were diluted in Laemmli sample buffer [final (1 x) concentration: 31.5 mM 2-Amino-2-(hydroxymethyl)propane-1,3-diol (Tris), 1 % (w/v) sodium dodecyl sulfate (SDS), 10 % (v/v) glycerol, 0.001% (w/v) 3,3-Bis(3,5-dibromo-4-hydroxyphenyl)-2,1λ^6^-benzoxathiole-1,1(3*H*)-dione (bromophenol blue), 5 % (v/v) 2-mercaptoethanol; pH 6.8] and denatured at 55°C for 20 min (unless indicated otherwise). Samples were resolved by SDS-polyacrylamide gel electrophoresis (SDS-PAGE) with self-made Bis(2-hydroxyethyl)amino-tris(hydroxymethyl)methan (BisTris; 250 mM) based 4-20 % polyacrylamide (Rotiphorese Gel 30; Carl Roth) gradient gels and run (80-120 V; 90 min) in NuPAGE MOPS SDS running buffer (Thermo Fisher Scientific; Cat#NP000102) using Mini-PROTEAN Tetra Vertical Electrophoresis Cells (Bio-Rad, Cat#1658004). For membrane fractionation experiments, RFP-fluorescence of mDia1 variants in respective fractions was imaged in-gel on a ChemiDoc XRS+ (Bio-Rad) controlled by the Image Lab software (version 6.0.1) utilizing the Alexa546 preset (605/50 Filter3; Light Green Epi illumination). Separated proteins were wet-blotted (110 V; 90 min; 4°C) on fluorescence-optimized polyvinylidene difluoride (PVDF) membranes (Immobilon-FL; Merck; IPFL00010) in transfer buffer (25 mM Tris (pH 7.6), 192 mM glycine, 20% (v/v) methanol, 0.03% (w/v) SDS). Subsequently, membranes were blocked with blocking buffer [5 % bovine serum albumin (BSA), in Tris-buffered saline (TBS) containing 0.01% Tween 20 (TBS-T)] for 1 h at RT and incubated (4 °C; overnight) with primary antibodies under constant agitation at indicated dilutions (s. Key Resource Table) in blocking buffer. Membranes were washed thrice with TBS-T and incubated with corresponding pairs of IRDye 680RD-or 800CW-conjugated secondary antibodies in TBS-T for 1 h at RT. After three washes with TBS-T, bound antibodies were visualized by the Odyssey Fc Imaging System (LI-COR Biosciences) controlled and analyzed by Image Studio Lite (Version 5.2.5). For colorimetric analysis of protein levels, intensity of bands was measured by assigning shapes of equal size to all lanes at similar heights and subtracting the individual background for each shape. Signals were normalized to controls on the same blot. The PageRuler Prestained (Thermo Fisher Scientific; Cat#26616) was used as a ladder to control for protein size.

### Immunoprecipitation from synaptosomes

To immunoprecipitate synaptic mDia1, synaptosomes (P2’) were prepared as follows: One mouse brain (p28) was homogenized (900 rpm; 12 strokes with glass-teflon homogenizer) in 7 mL of ice-cold homogenization buffer [(320 mM sucrose, 4 mM HEPES; pH 7.4) supplemented with mammalian protease inhibitor cocktail (PIC)] at 4°C. Large cellular debris and nuclei were sedimented by centrifugation at 900 g for 10 min at 4°C. The supernatant was further centrifuged at 12500 g for 15 min at 4°C. The resulting pellet (P2) was resuspended in 15 mL of homogenization buffer and pelleted at 12500 g for 15 min at 4°C yielding the crude synaptosomal fraction P2’. Subsequently, the pellet was resuspended in 2 mL of immunoprecipitation buffer [20 mM HEPES, 130 mM NaCl, 2 mM MgCl_2_, 1 % (w/v) 3-[Dimethyl[3-(3α,7α,12α-trihydroxy-5β-cholan-24-amido)propyl]azaniumyl]propane-1-sulfonate (CHAPS), PIC, phosphatase inhibitor cocktail II and III (Sigma-Aldrich); pH 7.4] and lysed for 30 min at 4°C under light agitation. The lysate was cleared by centrifugation at 15000 g at 4°C and protein concentration was measured using the BCA assay.

For identification of the protein environment of mDia1, P2’ lysate (2 mg; 2 g/L) was incubated with either 2 μg of anti-mDia1 antibody (BD Biosciences, Cat# 610848) or equal amounts of immunoglobulin G (IgG) isotype control mouse antibody (Thermo Fisher Scientific; Cat# 31903; RRID:AB_10959891) for 1 h at 4°C before the addition of 25 μL of Pierce Protein A/G Magnetic Beads (Thermo Fisher Scientific) for an additional 2 h under constant rotation. Subsequently, unbound proteins were removed from the beads by three consecutive wash cycles with immunoprecipitation buffer. Bound proteins were eluted and boiled (10 min; 95°C) in 2x Laemmli buffer, resolved by SDS-PAGE and analyzed by immunoblotting.

### LC-MS analysis of protein environment of mDia1

For the analysis of interaction partners of synaptic mDia1, mDia1 was immunoprecipitated from synaptosomes as described above. Eluted proteins were reduced (5 mM dithiothreitol; 30 min at 55°C), alkylated (15 mM iodoacetamide; 20 min at RT in the dark), and submitted to LC-MS analysis: Proteins were subjected to SDS-PAGE following excision of 3 bands per lane and in-gel digestion of proteins by trypsin (1:100 (w/w); overnight at 37°C). Resulting tryptic peptides were separated by reverse-phase high-performance liquid chromatography (RP-HPLC; Ultimate^TM^ 3000 RSLCnano system; Thermo Scientific) using a 50 cm analytical column (in-house packed with Poroshell 120 EC-C18; 2.7 µm; Agilent Technologies) with a 120 min gradient. RP-HPLC was coupled on-line to an Orbitrap Elite mass spectrometer (Thermo Fisher Scientific) that performed precursor ion (MS1) scans at a mass resolution of 60000, while fragment ion (MS2) scans were acquired with an automatic gain control (AGC) target of 5 x 10^3^ and a maximum injection time of 50 ms. Data analysis including label free quantification was performed with MaxQuant (version 1.6.1.0) using the following parameters: The initial maximum mass deviation of the precursor ions was set at 4.5 parts per million (ppm), and the maximum mass deviation of the fragment ions was set at 0.5 Da. Cysteine carbamidometyl and propionamide, Methionine oxidation and N-terminal acetylation were set as variable modifications. For identification of proteins, data were searched against the SwissProt murine database (Mouse_2016oktuniprot-proteome%3AUP000000589.fasta). False discovery rates were < 1 % at the protein level based on matches to reversed sequences in the concatenated target-decoy database. The statistical analysis was done with Perseus (version 1.6.7.0).

### Effector pulldown assays

To analyze the activity of small Rho GTPases, pulldowns utilizing effector protein domains which exclusively bind to their GTP-bound forms were performed: Neurons (200 000 cells) were washed with ice-cold PBS and harvested in GTPase lysis buffer (50 mM HEPES, 500 mM NaCl, 10mM MgCl_2_, PIC, phosphatase Inhibitor cocktails II + III; pH 7.4). Cells were lysed for 5 min under repetitive mixing before centrifugation (15000 g; 5 min; 4°C). Cleared lysates were incubated with Rhotekin-Rho binding domain (RBD) beads (60 µg; Cytoskeleton Inc.) or with PAK-p21 binding domain (PBD) beads (20 µg; Cytoskeleton Inc.) at 4°C under constant rotation to bind active RhoA or Cdc42 and Rac1, respectively. After 2 h, the beads were pelleted by centrifugation (1000 g; 1 min; 4°C) and washed with washing buffer (50 mM HEPES, 150 mM NaCl, 10 mM MgCl_2_; pH 7.4). After repeated centrifugation, unbound proteins were discarded with the supernatant, while bound proteins were eluted from the beads by addition of 2 x Laemmli buffer and boiling (10 min; 95°C). Finally, activity of small Rho GTPases was resolved by SDS-PAGE and analyzed by immunoblotting input and pulldown samples.

### Membrane fractionation

To characterize membrane association of the truncation mDia1 mutant, HEK293T cells were transfected with wild-type (mDia1-WT) or mDia1 truncation mutant (mDia1-ΔN) using CaCl_2_. 48 h after transfection cells were washed with ice-cold PBS and harvested in ice-cold resuspension buffer (20 mM HEPES, 130 mM NaCl, PIC; pH 7.4). Cells were lysed by forcing the suspension through 18-gauge syringes to crack the plasma membrane in between three freeze-thaw cycles in liquid nitrogen. Nuclei were removed by centrifugation at 1000 g for 5 min at 4°C. The supernatant (total lysate) was sedimented at 100000 g for 30 min at 4°C to yield the membrane fraction as the pellet and the cytosolic fraction as the high-speed supernatant. The pellet was resuspended in resuspension buffer to the same volume of the cytosolic fraction. Equal volumes of all fractions were analyzed by SDS-PAGE and immunoblotting to allow interpretation of protein enrichment in cytosolic or membrane fractions with respect to the total lysate.

### Statistical analysis

All data in this study are presented as the mean ± standard error of the mean (SEM) and was obtained from *N* independent experiments with a total sample number of *n* (e.g., number of images, videos, synapses, etc.) as annotated in the Figure legends. For analysis of protein levels in STED microscopy and synaptic structures in EM, statistical differences between groups were calculated considering *n*, while in all other experiments statistical differences were calculated between independent experiments *N* (In pHluorin/CypHer assays, at least 20 responding boutons/video were analyzed). For n > 100 or N > 5, data were tested for Gaussian distribution following D’Agostino-Pearson tests to determine parametric versus non-parametric statistical testing. The statistical significance between two groups was evaluated with either two-tailed unpaired student’s t-tests for normally distributed data or two-tailed unpaired Mann-Whitney tests, if data did not follow Gaussian distribution. In experiments that necessitated normalization (to 100 or 1) before analysis, one-sample t-tests or one-sample Wilcoxon rank tests were performed for normal and non-normal distributed data, respectively. The statistical significance between more than two experimental groups of normally distributed data was analyzed by one-way ANOVA, followed by a Tukey’s post hoc test, while Kruskal-Wallis tests with post hoc Dunn’s multiple comparison test were used when datasets did not follow Gaussian distribution.

Corresponding statistical tests are indicated in the Figure and significance levels are annotated as asterisks (*p < 0.05, **p < 0.01, ***p < 0.001, and ****p < 0.0001). Differences that are not significant are not stated or indicated as ns (p > 0.05). Statistical data evaluation was performed using GraphPad Prism 9.5.1 (733) and all calculated p-values are annotated in the source data table. All Figures were assembled using Affinity Designer (version 1.10.6.1665).

## Data availability

All data generated and analyzed in this study are included in the manuscript and supporting files. Numerical data as well as raw images of western blots are provided as source data files. Values from the same experiment (N) are shaded in similar colors.

## Contact for Reagent and Resource Sharing

Further information and requests for resources and reagents should be directed to and will be fulfilled by the corresponding contact V.H.

## ACKNOWLEDGEMENTS

We are indebted to Sabine Hahn, Delia Löwe and Silke Zillmann for expert technical assistance with the preparation of neuronal cultures. We further wish to thank Dr. Martin Lehmann (FMP, Berlin), Hannah Gelhaus and Gresy Bregu (both FU Berlin) for aid with STED imaging, Dr. Dmytro Puchkov (FMP Imaging Core Facility) for supervising electron microscopy analysis, and Heike Stephanowitz and Prof. Fan Liu (FMP Core Facility Proteomics) for proteomic analyses. Supported by grants from the Deutsche Forschungsgemeinschaft (SFB958/ TP A01) to V.H..

## LEGENDS for SUPPLEMENTS

### List of supplemental figures

**Figure 1-figure supplement 1 | Role of formins and mDia1/3 in SV endocytosis.**

**Figure 2-figure supplement 1 | SV depletion in mDia1/3-depleted neurons is activity dependent.**

**Figure 3-figure supplement 1 | mDia1 binds membranes and localizes to presynaptic endocytic sites.**

**Figure 4-figure supplement 1| mDia1 regulates presynaptic actin and SV endocytosis.**

**Figure 5-figure supplement 1 | RhoA/B regulate SV endocytosis.**

**Figure 6-figure supplement 1 | Cooperative action of mDia1/3 and Rac1 pathways in presynaptic endocytosis.**

**Figure 7-figure supplement 1 | mDia1/3 and Rac1 cooperatively regulate the SV cycle and presynaptic ultrastructure.**

**Supporting Table 1 - Oligonucleotides used in this study**

**Source Data Table**

**Raw uncropped immunoblot images (merged PDF)**

---

**Figure 1-Supplement 1.**
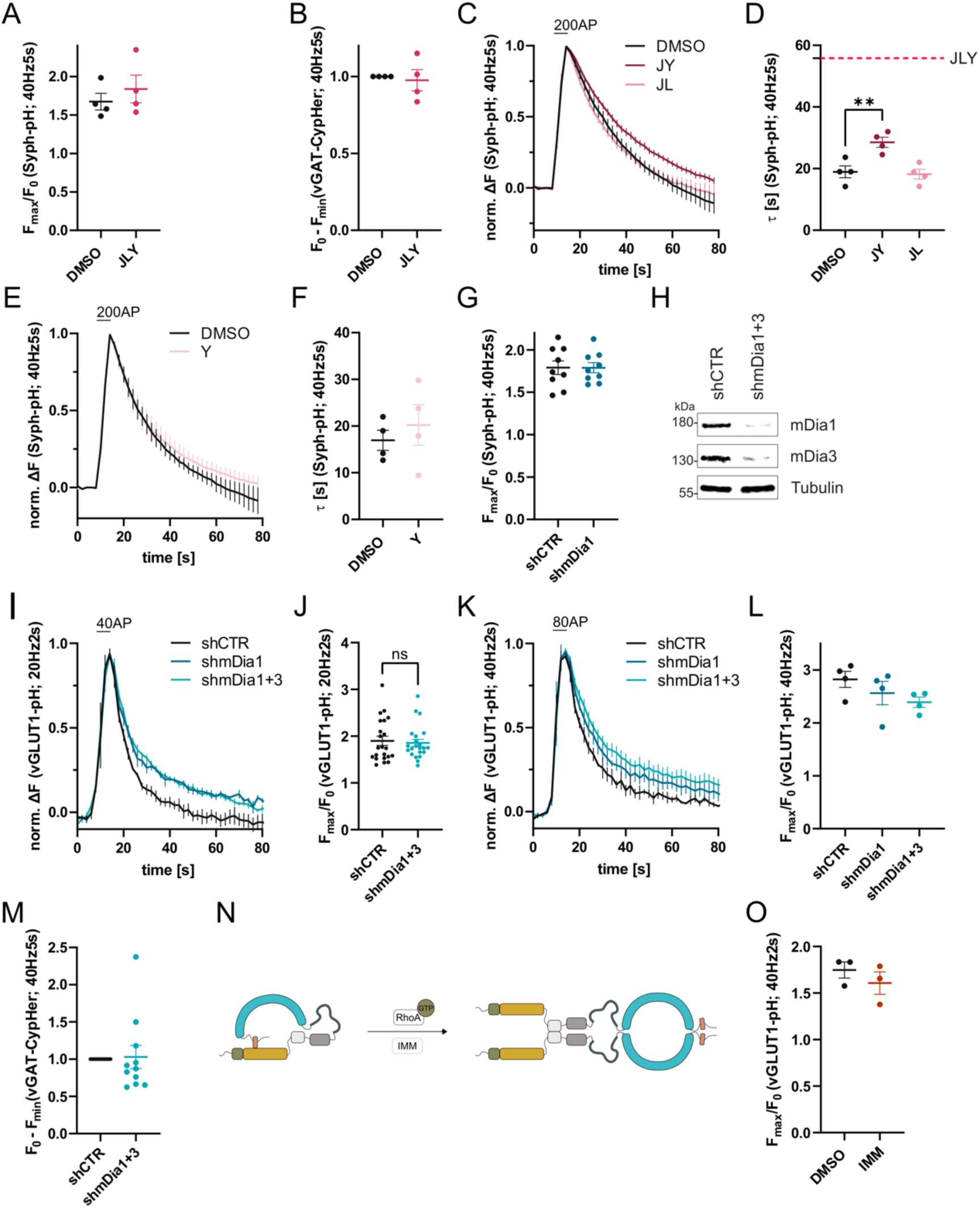
Role of formins and mDia1/3 in SV endocytosis. **(A)** Maxima of background-corrected Syph-pHluorin fluorescence traces (surface normalized) for neurons treated with 0.1 % DMSO (1.7 ± 0.1) or JLY cocktail (1.8 ± 0.2) in response to 200 AP stimulation (40 Hz, 5 s). Data represent mean ± SEM. N = 4 independent experiments from n_DMSO_ = 23 videos; n_JLY_ = 36 videos. **(B)** Minima of background-corrected vGAT-CypHer fluorescence traces (surface normalization) for neurons treated with 0.1 % DMSO or JLY cocktail (1.0 ± 0.1) in response to 200 AP stimulation (40 Hz, 5 s). Data represent mean ± SEM. Values for DMSO were set to 1. N = 4 independent experiments from n_DMSO_ = 23 videos; n_JLY_ = 29 videos. **(C)** Averaged normalized Syph-pH fluorescence traces from transfected hippocampal neurons treated with 0.1% DMSO, JY or JL combinations (containing 8 µM Jasplakinolide, 5 µM Latrunculin A and 10 µM Y-27632) stimulated with 200 APs (40 Hz, 5s). Data shown represent the mean ± SEM. N = 4 independent experiments from n_DMSO_ = 24 videos; n_JY_ = 19 videos; n_JL_ = 21 videos. **(D)** Endocytic decay constants of Syph-pHluorin traces in C: τ_DMSO_ = 19.0 ± 1.9 s; τ_JY_ = 28.5 ± 1.6 s; τ_JL_ = 18.2 ± 1.6 s; p_DMSO vs JY_ < 0.01, one-way ANOVA with Tukey’s post-test. Data shown represent mean ± SEM. **(E)** Averaged normalized Syph-pH fluorescence traces from transfected hippocampal neurons treated with 0.1% DMSO or 10 µM Y-27632 following 200 AP (40 Hz, 5s) stimulation. Data shown represent the mean ± SEM. N = 4 independent experiments from n_DMSO_ = 25 videos; n_Y_ = 18 videos. **(F)** Endocytic decay constants of Syph-pHluorin traces in C: τ_DMSO_ = 17.0 ± 2.2 s; τ_Y_ = 20.2 ± 4.3 s. Data shown represent mean ± SEM. **(G)** Maxima of background-corrected Syph-pHluorin fluorescence traces (surface normalized) for neurons transfected with shCTR (1.8 ± 0.1) or shmDia1 (1.8 ± 0.1) in response to 200 AP stimulation (40 Hz, 5 s). Data represent mean ± SEM. N = 9 independent experiments from n_shCTR_ = 49 videos; n_shmDia1_ = 42 videos. **(H)** Analysis of knockdown efficiency of lentiviral particles carrying shRNA against no mammalian target (shCTR) or mDia1 and mDia3 (shmDia1+3) in mouse hippocampal cultures harvested 12 days after transduction. Protein abundance of mDia1, mDia3 and Tubulin were immunoblotted with specific antibodies. **(I)** Averaged normalized vGLUT1-pHluorin fluorescence traces from stimulated (40 APs; 20 Hz, 2 s) hippocampal neurons transduced with lentiviruses encoding shCTR, shmDia1 or both shmDia1 and shmDia3 combined (shmDia1+3). Data represent mean ± SEM. N = 4 independent experiments from n_shCTR_ = 17 videos, n_shmDia1_ = 19 videos, n_shmDia1+3_ = 18 videos. The corresponding endocytic decay constants are shown in Figure 1G. **(J)** Maxima of background-corrected vGLUT1-pHluorin fluorescence traces (surface normalized) for neurons transduced with shCTR (1.9 ± 0.1) or shmDia1+3 (1.9 ± 0.1) in response to 40 AP stimulation (20 Hz, 2 s). Data represent mean ± SEM. N = 22 independent experiments from n_shCTR_ = 105 videos and n_shmDia1+3_ = 128 videos.**(K)** Averaged normalized vGLUT1-pHluorin fluorescence traces for neurons transduced with shCTR, shmDia1 or shmDia1+3 in response to 80 AP stimulation (40 Hz, 2 s). Data represent mean ± SEM. N = 4 independent experiments from n_shCTR_ = 12 videos; n_shmDia1_ = 15 videos; n_shmDia1+3_ = 18 videos. Corresponding endocytic decay constants are shown in Figure 1H. **(L)** Maxima of background-corrected vGLUT1-pHluorin fluorescence traces (surface normalized) for neurons transduced with shCTR (2.8 ± 0.2), shmDia1 (2.6 ± 0.2), or shmDia1+3 (2.4 ± 0.1) in response to 80 AP stimulation (40 Hz, 2 s). Data represent mean ± SEM. N = 4 independent experiments from n_shCTR_ = 12 videos; n_shmDia1_ = 15 videos and n_shmDia1+3_ = 18 videos. **(M)** Minima of background-corrected vGAT-CypHer fluorescence traces (surface normalized) for neurons transduced with shCTR or shmDia1+3 (1.0 ± 0.2) in response to 200 AP stimulation (40 Hz, 5 s). Data represent mean ± SEM. Values for shCTR were set to 1. N = 11 independent experiments from n_shCTR_ = 45 videos and n_shmDia1+3_ = 42 videos. **(N)** Schematic representation of the activation of mDia1. Binding of RhoA-GTP to the RBD domain (green) or application of mDia1 activator (IMM) competes with the intramolecular interaction of the N-terminal DID (yellow) with the C-terminal DAD (red) domain (see Figure 3A for domain structure). The release of autoinhibition leads to dimerization of mDia formins in solution. **(O)** Maxima of background-corrected vGLUT1-pHluorin fluorescence traces (surface normalized) for neurons treated with 0.1 % DMSO (1.7 ± 0.1) or mDia activator (IMM; 1.6 ± 0.1) in response to 80 AP stimulation (40 Hz, 2 s). Data represent mean ± SEM. N = 3 independent experiments from n_DMSO_ = 18 videos; n_IMM_ = 16 videos.

**Figure 2-Supplement 1.**
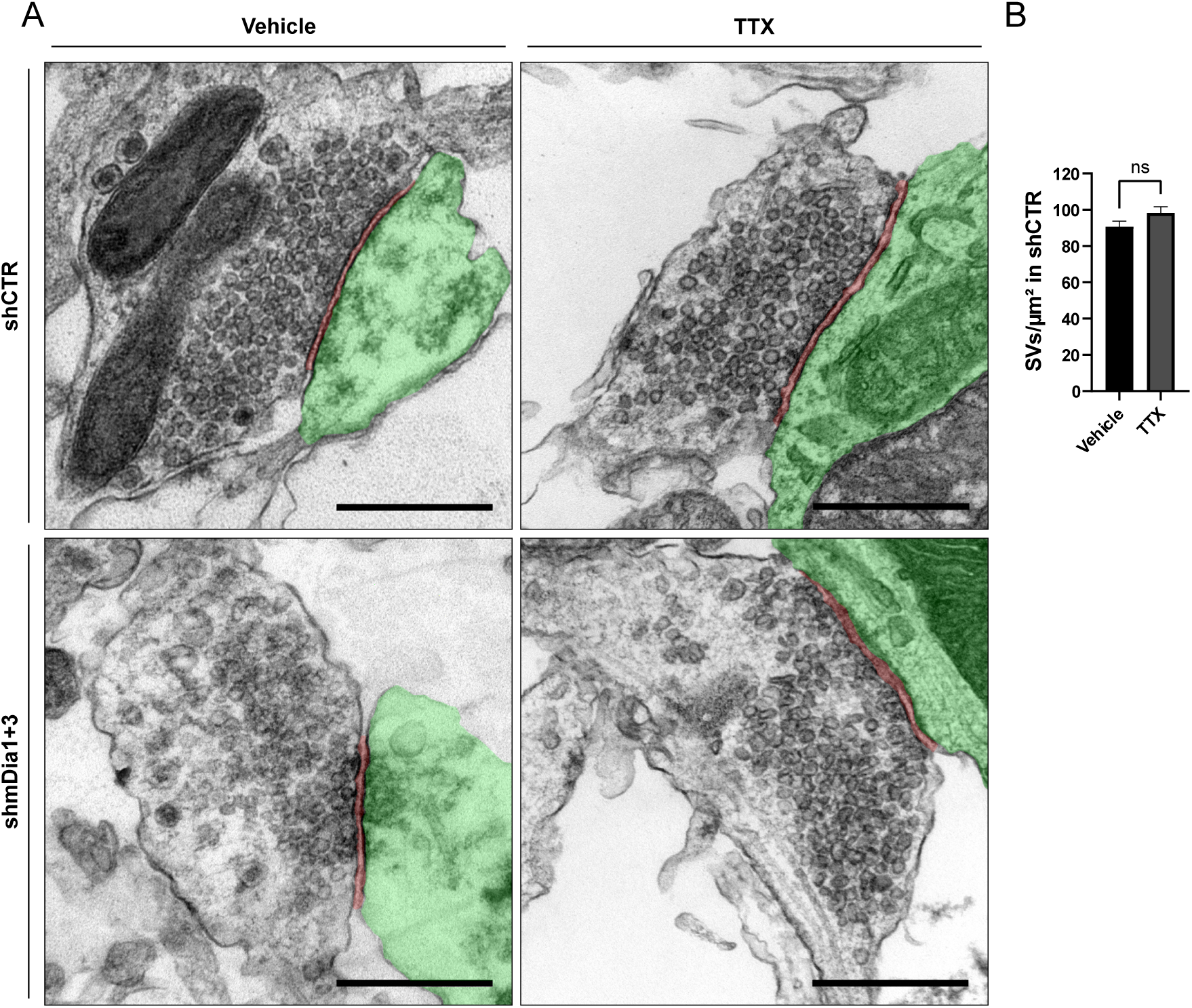
SV depletion in mDia1/3-depleted neurons is activity dependent. **(A)** Representative synaptic electron micrographs from neurons transduced with shCTR or shmDia1+3 and treated with 0.1% Vehicle or 1 µM TTX for 36 h before fixation. Postsynapse and postsynaptic cleft are colored in green and maroon, respectively. Scale bar, 250 nm. **(B)** Average number of SV per μm^2^ in synaptic boutons of neurons transduced with shCTR and treated with 0.1 % Vehicle (A; 90.7 ± 3.1) or 1 µM TTX (B; 98.4 ± 3.3) for 36 h before chemical fixation. Data shown represent the mean ± SEM from two independent experiments and n_Vehicle_ = 180, n_TTX_ = 204 synapses.

**Figure 3-Supplement 1.**
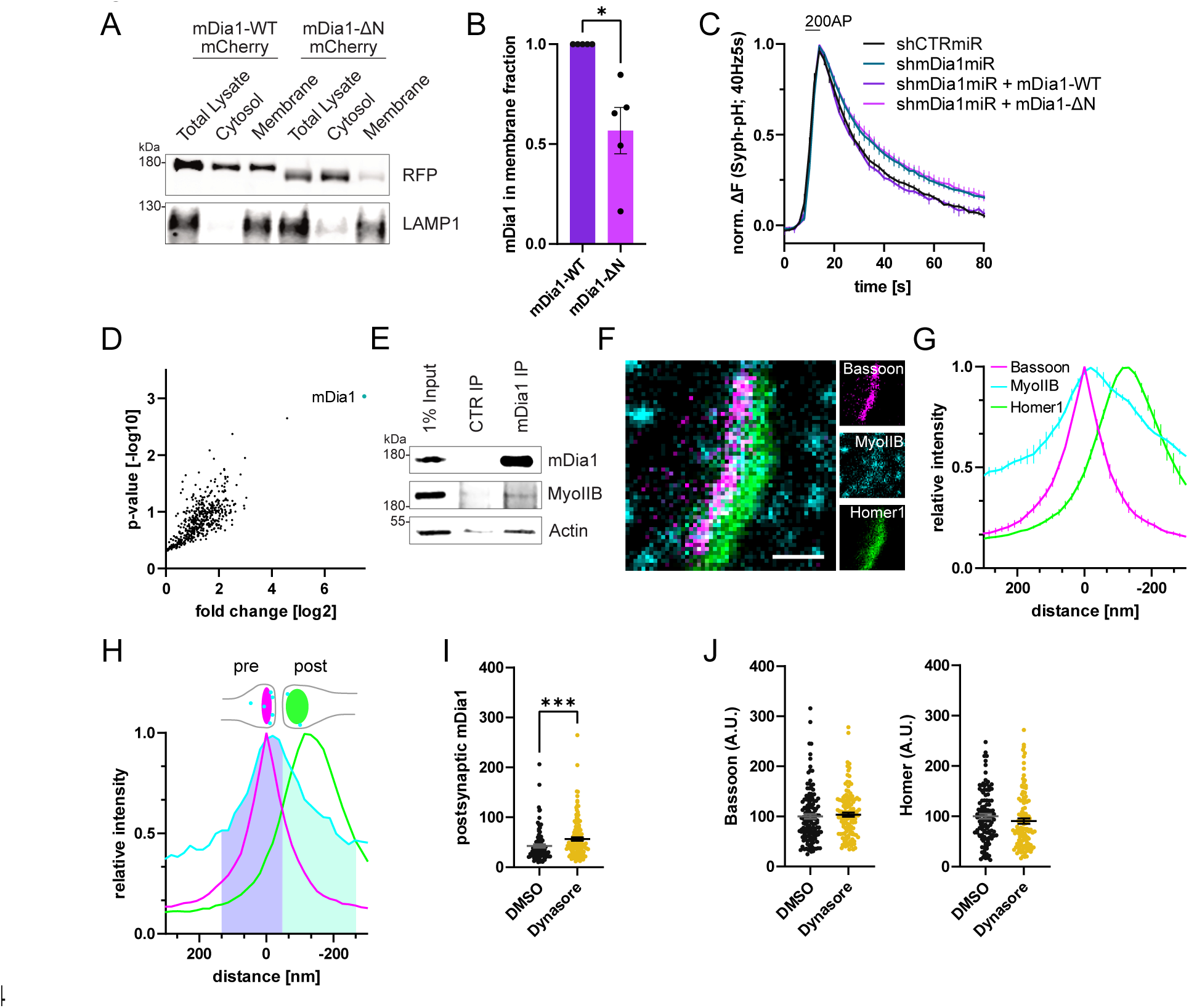
mDia1 binds membranes and localizes to presynaptic endocytic sites. **(A)** Membrane levels of mDia1-WT-mCherry versus mDia1-ΔN-mCherry proteins overexpressed in HEK293T cells. Membrane and cytosolic cellular fractions were isolated by ultracentrifugation and analyzed by immunoblotting with specific antibodies (LAMP1) and in-gel fluorescence of mCherry tags. **(B)** Densitometric quantification of mDia1-WT versus mDia1-ΔN (0.6 ± 0.1; p < 0.05, one sample t-test) membrane-associated protein levels. Data shown are normalized to mDia1-WT (set to 1) and expressed as mean ± SEM. Representative immunoblot is shown in A. N = 5 independent experiments. **(C)** Averaged normalized Synaptophysin-pHluorin fluorescence from stimulated (200 APs, 40 Hz, 5s) hippocampal neurons transfected with shCTRmiR or shmDia1miR. For rescue experiments, neurons were co-transfected with plasmids encoding mDia1-WT-mCherry, mDia1-ΔN-mCherry or mCherry alone (shCTRmiR & shmDia1miR). Endocytic decay constants are shown in Figure 3B. **(D)** Full volcano plot of proteins from Figure 3E associating with synaptic mDia1 analyzed by label-free proteomics of anti-mDia1 versus CTR immunoprecipitates from detergent-extracted mouse synaptosomes (P2’ fraction). The cyan dot shows the specific enrichment of mDia1 as the bait protein of the immunoprecipitation (p < 0.001, two-tailed student’s t-test). N = 3 independent experiments. **(E)** Endogenous co-immunoprecipitation of Myosin IIB by mDia1 from detergent-extracted mouse synaptosomes (P2’ fraction). Samples were analyzed by immunoblotting using specific antibodies against mDia1, Myosin IIB (MyoIIB) and β-Actin. **(F)** Representative three-channel time-gated STED image of a synapse from hippocampal cultures fixed and immunostained for Bassoon (presynaptic marker, magenta), Myosin IIB (cyan) and Homer1 (postsynaptic marker, green). Scale bar, 250 nm. **(G)** Averaged normalized line profiles for synaptic distribution of Myosin IIB and Homer1 relative to Bassoon (Maximum set to 0 nm). Data are expressed as mean ± SEM. N = 3 independent experiments from n = 267 synapses. **(H)** Rationale for quantification of presynaptic protein levels of interest. The presynapse was defined by the normalized Bassoon distribution (purple fraction, cutoff at the crossection with the Homer1 profile) and corresponding absolute individual synaptic line profiles were integrated. **(I)** Postsynaptic F-Actin levels in synapses treated with 0.1% DMSO (42.6 ± 3.4) or 80 µM Dynasore (56.5 ± 3.3; p < 0.001, Mann-Whitney test) for 10 min before fixation from Figure 3C. Data shown are normalized to presynaptic DMSO values from Figure 3H (set to 100) and expressed as mean ± SEM. N = 3 independent experiments from n_DMSO_ = 92 synapses, n_Dynasore_ = 135 synapses. **(J)** Quantification of Bassoon and Homer1 levels in synapses treated with 0.1% DMSO (100.0 ± 4.5 for Bassoon; 100.0 ± 4.3 for Homer1) or 80 µM Dynasore (103.7 ± 4.1 for Bassoon; 98.6 ± 5.3) for 10 min before fixation. Data shown are normalized to DMSO values (set to 100) and expressed as mean ± SEM. N = 3 independent experiments from n_DMSO_ = 132 synapses, n_Dynasore_ = 128 synapses.

**Figure 4-Supplement 1.**
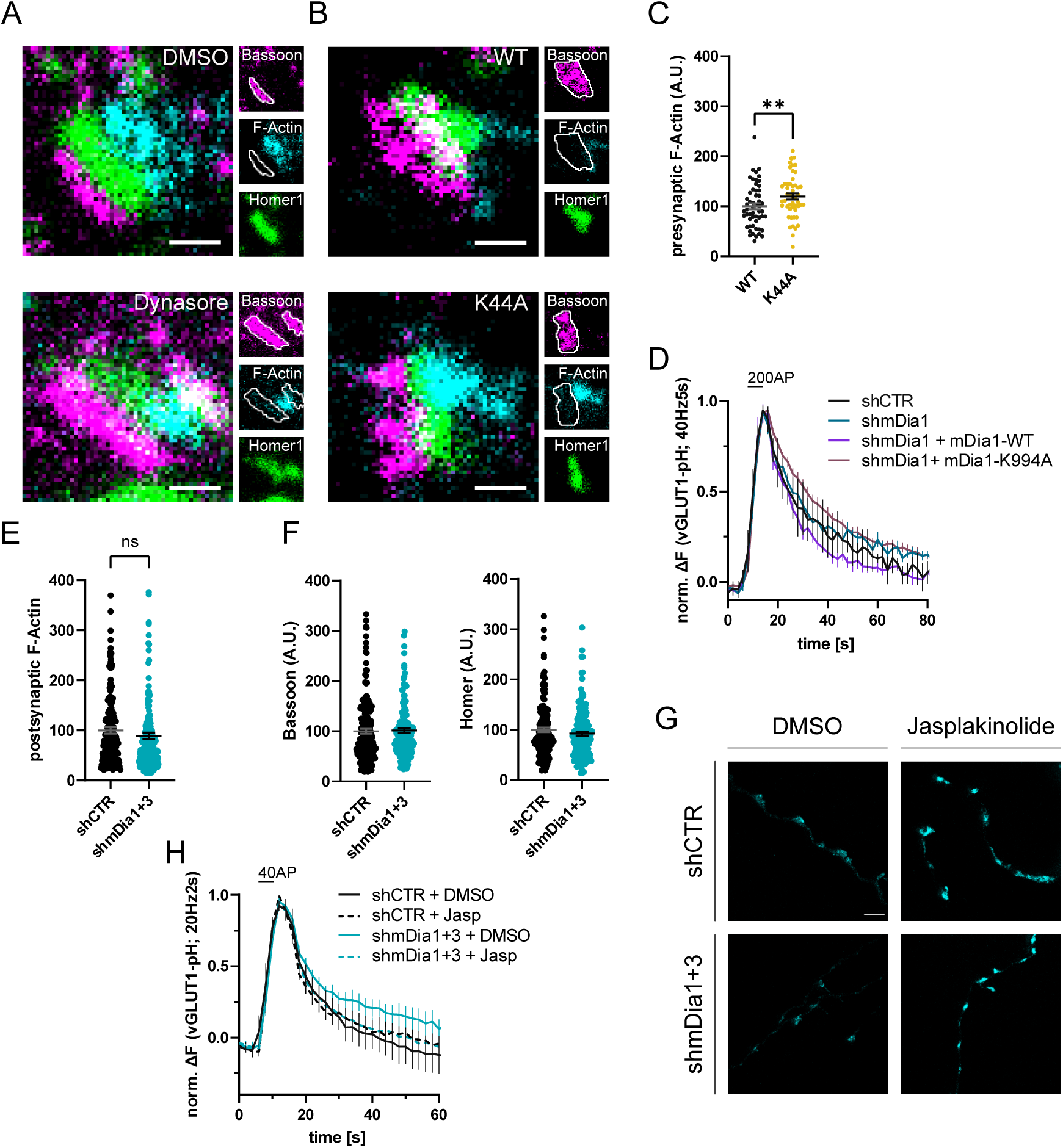
mDia1 regulates presynaptic actin and SV endocytosis. **(A)** Representative three-channel time-gated STED images of synapses from hippocampal cultures treated with 0.1% DMSO or 80 µM Dynasore for 10 min. Cells were fixed and stained for Bassoon (magenta), F-Actin (cyan) and Homer1 (green). Scale bar, 250 nm. Corresponding analysis of presynaptic F-Actin levels is shown in Figure 4C. **(B)** Representative three-channel time-gated STED images of synapses from hippocampal cultures transduced with Dynamin1-WT or Dynamin1-K44A. Cells were fixed and stained for Bassoon (magenta), F-Actin (cyan) and Homer1 (green). Scale bar, 250 nm. **(C)** Presynaptic F-Actin levels in synapses from neurons transduced with Dynamin1-WT (100 ± 5.9) or Dynamin1-K44A (119.8 ± 6.2, p < 0.01, one sample t-test) in B. Absolute line profiles of F-Actin overlapping with Bassoon (presynapse) distribution were integrated. Data shown are normalized to WT (set to 100) and expressed as mean ± SEM. n_WT_ = 54 synapses, n_K44A_ = 49 synapses. **(D)** Averaged normalized vGLUT1-pHluorin fluorescence traces for neurons transduced with shCTR or shmDia1 in response to 200 AP (40 Hz, 5s) stimulation. For rescue purposes, cells were co-transduced with mDia1-WT-SNAP or mDia1-K994A-SNAP. Data are expressed as mean ± SEM. N = 6 independent experiments from n_shCTR_ = 21 videos; n_shmDia1_ = 21 videos; n_shmDia1 + mDia1-WT_ = 16 videos; n_shmDia1 + mDia1-K994A_ = 19 videos. Corresponding endocytic decay constants are shown in Figure 4D. **(E)** Postsynaptic F-Actin levels in synapses transduced with shCTR (100.0 ± 6.4) or shmDia1+3 (89.3 ± 6.4) from Figure 4A,E. Data shown are normalized to shCTR values (set to 100) and expressed as mean ± SEM. N = 3 independent experiments from n_shCTR_ = 206 synapses, n_shmDia1+3_ = 135 synapses. **(F)** Quantification of Bassoon and Homer1 levels in synapses transduced with shCTR (100.0 ± 4.7 for Bassoon; 100.0 ± 4.5 for Homer1) or shmDia1+3 (101.4 ± 4.8 for Bassoon; 92.4 ± 4.0). Data shown are normalized to DMSO values (set to 100) and expressed as mean ± SEM. N = 3 independent experiments from n_shCTR_ = 158 synapses and n_shmDia1+3_ = 159 synapses. **(G)** Representative STED images of endogenous β-Actin in vGLUT1 positive synapses in hippocampal neurons transduced with shCTR or shmDia1+3 and treated with 0.1 % DMSO or 1 µM Jasplakinolide for 45 min. Neurons were co-transfected with pOrange-GFP-β-Actin knockin and vGLUT1-mCherry plasmids before fixation and immunostaining. Scale bar, 2.5 µm. **(H)** Averaged normalized vGLUT1-pHluorin fluorescence traces for neurons transduced with shCTR or shmDia1+3 in response to 40 AP (20 Hz, 2s) stimulation. Neurons were pre-incubated with 0.1 % DMSO or 1 µM Jasplakinolide (Jasp) for 30 min in the cell media before imaging. Data are expressed as mean ± SEM. N = 6 independent experiments from n_shCTR + DMSO_ = 32 videos, n_shmDia1+3 + DMSO_ = 35 videos, n_shCTR + Jasp_ = 33 videos; n_shmDia1+3 + Jasp_ = 34 videos. The corresponding endocytic decay constants are shown in Figure 4I.

**Figure 5-Supplement 1.**
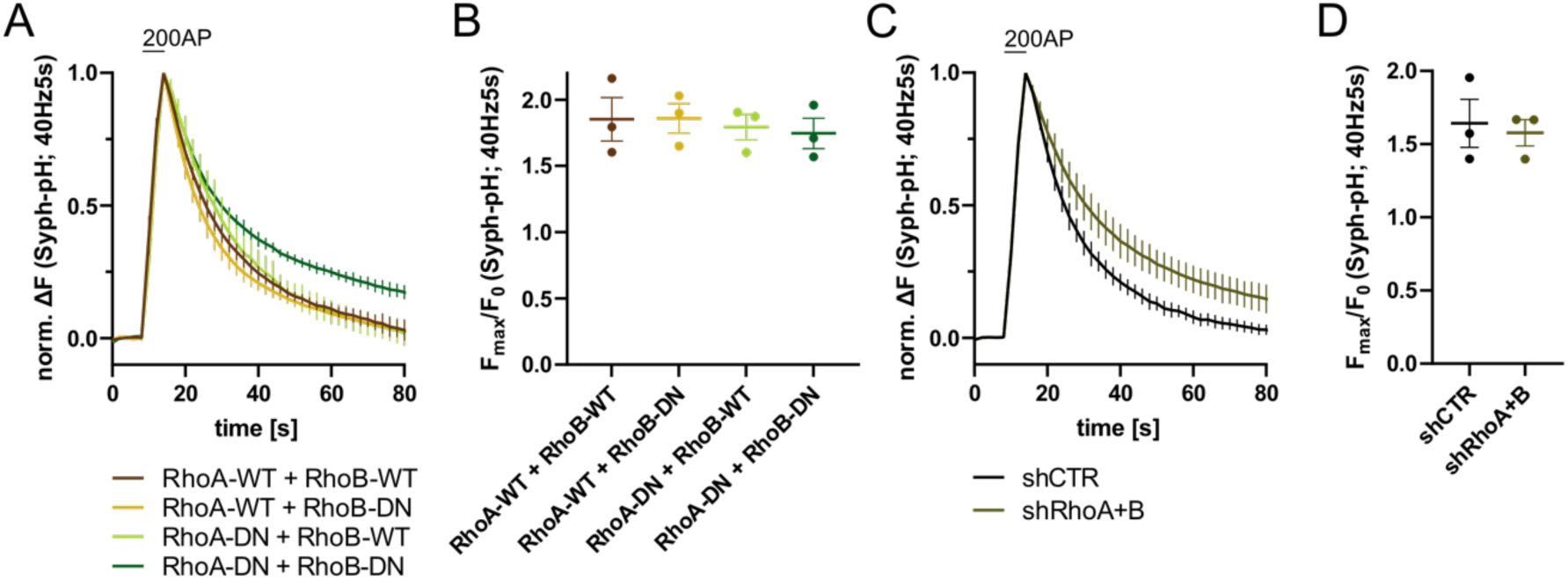
RhoA/B regulate SV endocytosis. **(A)** Averaged normalized Synaptophysin-pHluorin fluorescence traces from stimulated (200 APs; 40 Hz, 5s) hippocampal neurons transfected with plasmids encoding the indicated combinations of WT or DN RhoA and RhoB variants. Data represent mean ± SEM. N = 3 independent experiments from n_RhoA-WT + RhoB-WT_ = 21 videos, n_RhoA-DN + RhoB-WT_ = 31 videos, n_RhoA-WT + RhoB-DN_ = 23 videos, n_RhoA-DN + RhoB-DN_ = 22 videos. Endocytic decay constants are shown in Figure 5D. **(B)** Maxima of background-corrected Syph-pHluorin fluorescence traces (surface normalized) for neurons transfected with indicated combinations of WT or DN RhoA and RhoB variants (1.9 ± 0.2 for RhoA-WT + RhoB-WT; 1.9 ± 0.1 for RhoA-WT + RhoB-DN; 1.8 ± 0.1 for RhoA-DN + RhoB-WT; 1.7 ± 0.1 for RhoA-DN + RhoB-DN) in response to 200 AP stimulation (40 Hz, 5 s). Data represent mean ± SEM. **(C)** Averaged normalized Synaptophysin-pHluorin fluorescence traces from stimulated (200 APs; 40 Hz, 5s) hippocampal neurons transfected with shRNA against no mammalian target (shCTR) or against RhoA and RhoB (shRhoA+B). Data represent mean ± SEM. N = 3 independent experiments from n_shCTR_ = 27 videos, n_shRhoA+B_ = 25 videos. **(D)** Maxima of background-corrected Syph-pHluorin fluorescence traces (surface normalized) for neurons transfected with shCTR (1.6 ± 0.2) or shRhoA+B (1.6 ± 0.1) in response to 200 AP stimulation (40 Hz, 5 s). Data represent mean ± SEM.

**Figure 6-Supplement 1.**
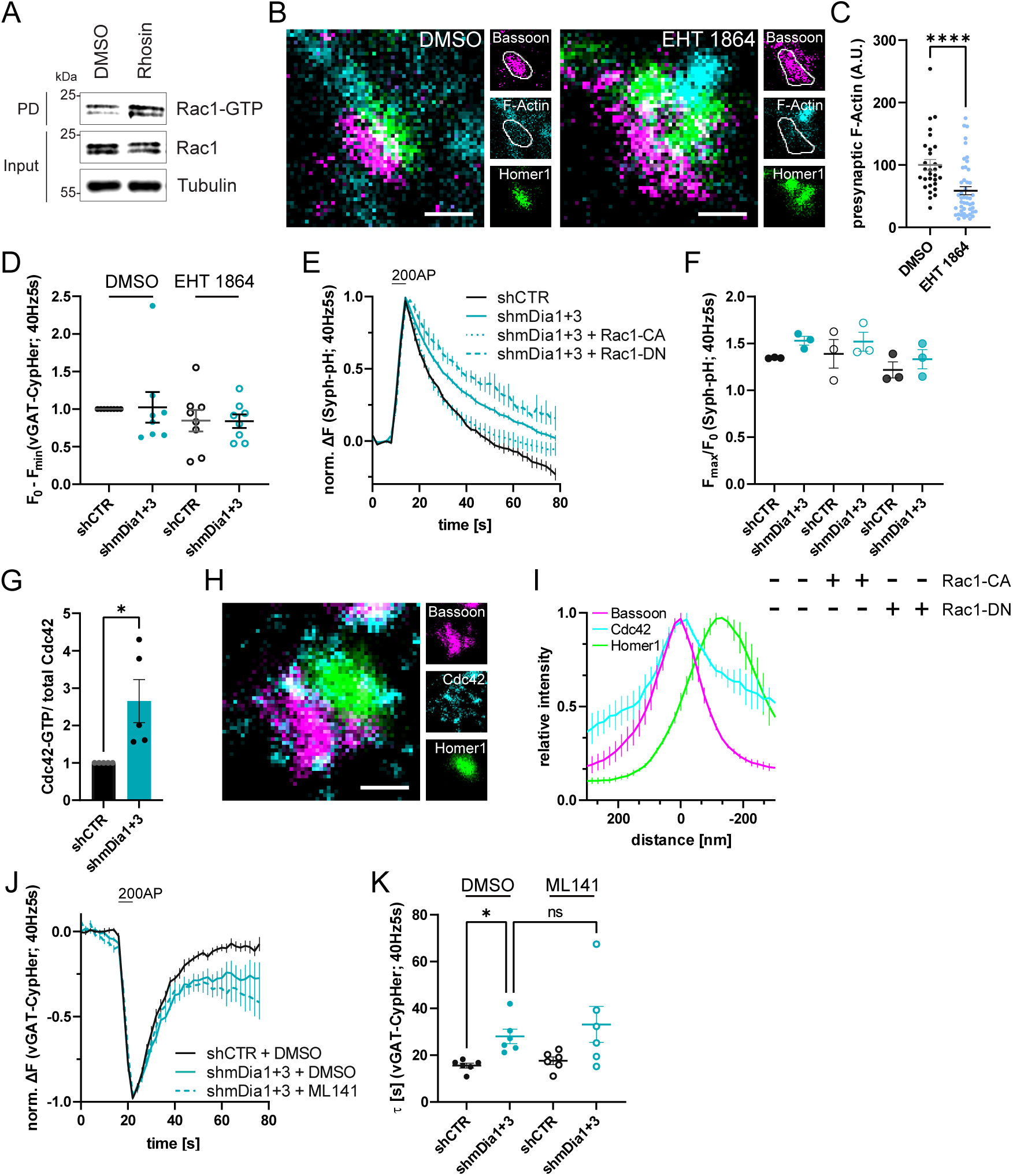
Cooperative action of mDia1/3 and Rac1 pathways in presynaptic endocytosis. **(A)** Analysis of Rac1 activity by Rac1-GTP pulldown (PD) from whole-cell lysates (input) of mouse hippocampal cultures upon inhibition of Rho activity utilizing immobilized PAK as bait. Cells were treated with 0.1 % DMSO or 10 µM Rho Inhibitor (Rhosin) for 2 h before harvest. Samples were analyzed by immunoblotting for Rac1 and Tubulin using specific antibodies. Input, 10% of material used for the pulldown. Contrast of pulldown and input blots was seperatly adjusted for visualisation purposes. **(B)** Representative three-channel time-gated STED images of synapses from hippocampal cultures treated with 0.1 % DMSO or 10 µM Rac1 Inhibitor (EHT 1864) for 2 h. Cells were fixed and stained for Bassoon (magenta), F-Actin (cyan) and Homer1 (green). Scale bar, 250 nm. **(C)** Presynaptic F-Actin levels in synapses of neurons treated with 0.1 % DMSO (100 ± 8.5) or 10 µM Rac1 Inhibitor (EHT 1864; 58.6 ± 6.5; p < 0.0001, one sample Wilcoxon test) for 2 h. Line profiles of F-Actin overlapping with Bassoon (presynapse) distribution were integrated. Data shown are normalized to DMSO (set to 100) and expressed as mean ± SEM. n_DMSO_ = 30, n_EHT 1864_ = 46 from two independent experiments. **(D)** Minima of background-corrected vGAT-CypHer fluorescence traces (surface normalized) for neurons treated with 0.1 % DMSO (1.0 ± 0.2 for shmDia1+3) or 10 µM Rac1 Inhibitor (EHT 1864; 0.8 ± 0.1 for shCTR; 0.8 ± 0.1 for shmDia1+3) in response to 200 AP stimulation (40 Hz, 5 s). Data represent mean ± SEM. Values were normalised to DMSO treated shCTR (set to 1). N = 8 independent experiments from n_shCTR + DMSO_ = 46 videos, n_shmDia1+3 + DMSO_ = 45 videos, n_shCTR + EHT 1864_ = 42 videos, n_shmDia1+3 + EHT 1864_ = 43 videos. **(E)** Averaged normalized Synaptophysin-pHluorin fluorescence traces from stimulated (200 APs; 40 Hz, 5s) hippocampal neurons transduced with lentiviruses encoding shCTR or shmDia1+3 and transfected with plasmids for expression of constitutively-active Rac1 (Rac1-CA; Q61L variant) or dominant-negative Rac1 (Rac1-DN; T17N variant). Data represent mean ± SEM. N = 3 independent experiments from n_shCTR_ = 12 videos, n_shmDia1+3_ = 23 videos, n_shCTR + Rac1-CA_ = 10 videos, n_shmDia1+3 + Rac1-CA_ = 14 videos, n_shCTR + Rac1-DN_ = 9 videos; n_shmDia1+3 + Rac1-DN_ = 13 videos. The corresponding endocytic decay constants are shown in Figure 6H. **(F)** Maxima of background-corrected Synaptophysin-pHluorin fluorescence traces (surface normalized maximum values of traces shown in E) from stimulated (200 APs; 40 Hz, 5s) hippocampal neurons transduced with lentiviruses encoding shCTR (F_max_/F_0_ = 1.3 ± 0.0) or shmDia1+3 (F_max_/F_0_ = 1.5 ± 0.0) and transfected with plasmics encoding CA (F_max_/F_0 shCTR + Rac1-CA_ = 1.4 ± 0.2; F_max_/F_0 shmDia1+3 + Rac1-CA_ = 1.5 ± 0.1) or DN versions ((F_max_/F_0 shCTR + Rac1-DN_ = 1.2 ± 0.1; F_max_/F_0 shmDia1+3 + Rac1-DN_ = 1.3 ± 0.1) of Rac1. Data represent mean ± SEM. **(G)** Densitometric quantification of Cdc42-GTP normalized to total Cdc42 levels in lysates from shmDia1+3 neurons (2.7 ± 0.6; p < 0.05, one sample t-test).Values for shCTR were set to 1. Data are expressed as mean ± SEM from N = 3 independent experiments. **H)** Representative three-channel time-gated STED image of a synapses from hippocampal mouse cultures, fixed and immunostained for Bassoon (magenta), Cdc42 (cyan) and Homer1 (green). Scale bar, 250 nm. **(I)** Averaged normalized line profiles for synaptic distribution of Cdc42 and Homer1 relative to Bassoon (Maximum set to 0 nm). Data are expressed as mean ± SEM (N = 3; n = 96 synapses). **(J)** Averaged normalized vGAT-CypHer fluorescence traces for neurons transduced with shCTR or shmDia1+3 in response to 200 AP (40 Hz, 5s) stimulation. Cells were acutely treated with 0.1 % DMSO or 10 µM Cdc42 Inhibitor (ML141) in the imaging buffer. Data shown represent the mean ± SEM. N = 6 independent experiments from n_shCTR + DMSO_ = 31 videos, n_shmDia1+3 + DMSO_ = 33 videos, n_shmDia1+3 + ML141_ = 32 videos. **(K)** Endocytic decay constants of vGAT-CypHer traces in J: τ_shCTR + DMSO_ = 15.6 ± 1.0 s, τ_shmDia1+3 + DMSO_ = 28.0 ± 3.1 s, τ_shCTR + ML141_ = 17.6 ± 1.6 s, τ_shmDia1+3 + ML141_ = 33.1 ± 7.7 s; p_shCTR + DMSO vs shmDia1+3 + DMSO_ < 0.01, Kruskal-Wallis test with Dunn’s post-test. Data shown represent the mean ± SEM. N = 6 independent experiments from n_shCTR + DMSO_ = 31 videos, n_shmDia1+3 + DMSO_ = 33 videos, n_shCTR + ML141_ = 29 videos, n_shmDia1+3 + ML141_ = 32 videos.

**Figure 7-Supplement 1.**
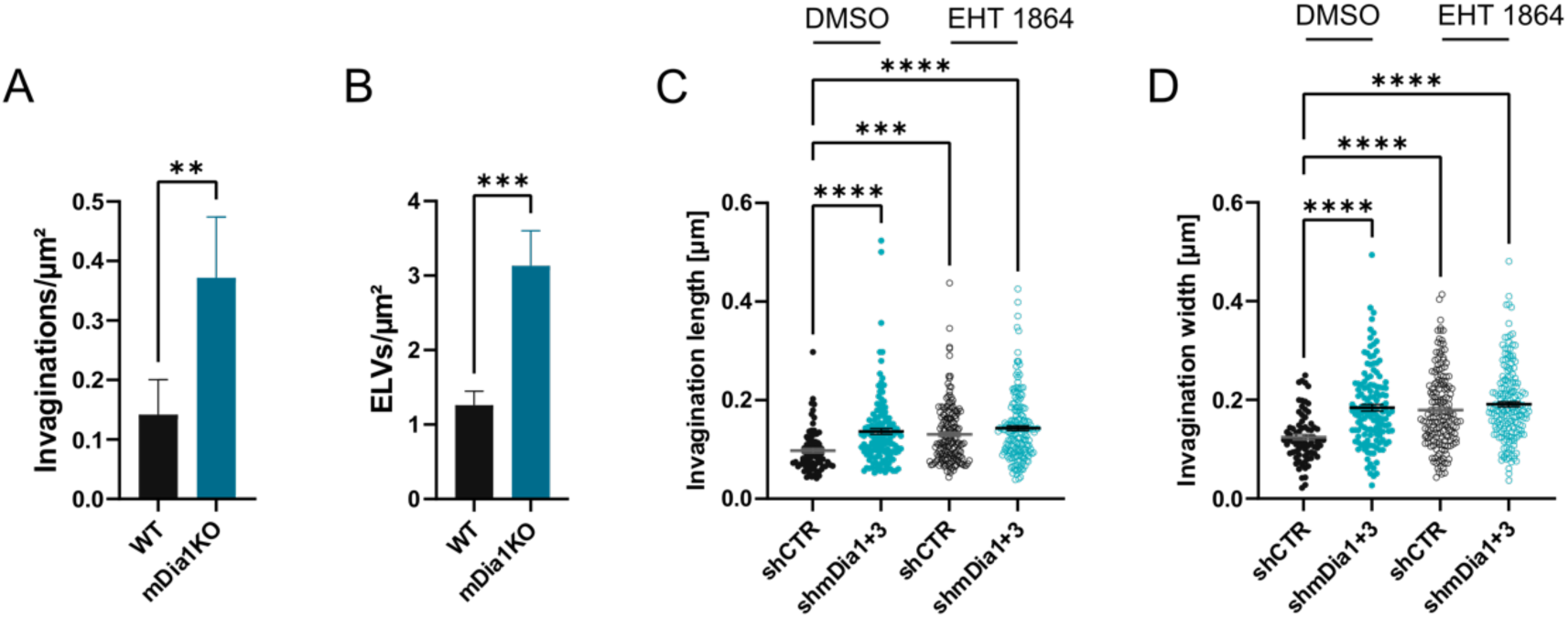
mDia1/3 and Rac1 cooperatively regulate the SV cycle and presynaptic ultrastructure. **(A)** Average number of invaginations per μm^2^ in WT (0.1 ± 0.1) and mDia1 KO (0.4 ± 0.1; p < 0.01, Mann-Whitney test) boutons. Data shown represent the mean ± SEM from n_WT_ = 103 synapses, n_KO_ = 96 synapses. **(B)** Average number of ELVs per μm^2^ in WT (1.3 ± 0.2) and mDia1KO (3.1 ± 0.5; p < 0.001, Mann-Whitney test) boutons. Data shown represent the mean ± SEM from n_WT_ = 103 synapses, n_KO_ = 96 synapses. **(C)** Average invagination length in shCTR and shmDia1+3 boutons treated with 0.1 % DMSO (97.6 ± 4.5 nm for shCTR; 136.8 ± 6.0 nm for shmDia1+3, p_shCTR + DMSO vs shmDia1+3 + DMSO_ < 0.0001) or 10 µM EHT 1864 (130.9 ± 4.5 nm for shCTR, p_shCTR + DMSO vs shCTR + EHT 1864_ < 0.001; 143.1 ± 4.9 nm for shmDia1+3, p_shCTR + DMSO vs shmDia1+3 + EHT 1864_ < 0.0001, Kruskal-Wallis test with Dunn’s post-test) for 2 h before chemical fixation. Data represent mean ± SEM from n_shCTR + DMSO_ = 77 invaginations, n_shmDia1+3 + DMSO_ = 141 invaginations, n_shCTR + EHT 1864_ = 176 invaginations, n_shmDia1+3 + EHT 1864_ = 189 invaginations. **(D)** Average invagination width in shCTR and shmDia1+3 boutons treated with 0.1 % DMSO (124.5 ± 5.6 nm for shCTR; 184.1 ± 6.6 nm for shmDia1+3, p_shCTR + DMSO vs shmDia1+3 + DMSO_ < 0.0001) or 10 µM EHT 1864 (179.0 ± 5.8 nm for shCTR, p_shCTR + DMSO vs shCTR + EHT 1864_ < 0.0001; 191.0 ± 5.4 nm for shmDia1+3, p_shCTR + DMSO vs shmDia1+3 + EHT 1864_ < 0.0001, Kruskal-Wallis test with Dunn’s post-test) for 2 h before chemical fixation. Data represent mean ± SEM from n_shCTR + DMSO_ = 77 invaginations, n_shmDia1+3 + DMSO_ = 141 invaginations, n_shCTR + EHT 1864_ = 176 invaginations, n_shmDia1+3 + EHT 1864_ = 189 invaginations.

## REFERENCES

Anes E, Kuhnel MP, Bos E, Moniz-Pereira J, Habermann A, Griffiths G (2003) Selected lipids activate phagosome actin assembly and maturation resulting in killing of pathogenic mycobacteria. Nat Cell Biol 5: 793–802

Bingham D, Jakobs CE, Wernert F, Boroni-Rueda F, Jullien N, Schentarra EM, Friedl K, Da Costa Moura J, van Bommel DM, Caillol G et al (2023) Presynapses contain distinct actin nanostructures. J Cell Biol 222

Bleckert A, Photowala H, Alford S (2012) Dual pools of actin at presynaptic terminals. J Neurophysiol 107: 3479–3492

Bloom O, Evergren E, Tomilin N, Kjaerulff O, Low P, Brodin L, Pieribone VA, Greengard P, Shupliakov O (2003) Colocalization of synapsin and actin during synaptic vesicle recycling. J Cell Biol 161: 737–747

Borst JG, Soria van Hoeve J (2012) The calyx of Held synapse: from model synapse to auditory relay. Annu Rev Physiol 74: 199–224

Boucrot E, Ferreira AP, Almeida-Souza L, Debard S, Vallis Y, Howard G, Bertot L, Sauvonnet N, McMahon HT (2015) Endophilin marks and controls a clathrin-independent endocytic pathway. Nature 517: 460–465

Boulant S, Kural C, Zeeh JC, Ubelmann F, Kirchhausen T (2011) Actin dynamics counteract membrane tension during clathrin-mediated endocytosis. Nat Cell Biol 13: 1124–1131

Chanaday NL, Cousin MA, Milosevic I, Watanabe S, Morgan JR (2019) The Synaptic Vesicle Cycle Revisited: New Insights into the Modes and Mechanisms. J Neurosci 39: 8209–8216

Chanaday NL, Kavalali ET (2018) Optical detection of three modes of endocytosis at hippocampal synapses. Elife 7

Chandrasekar I, Huettner JE, Turney SG, Bridgman PC (2013) Myosin II regulates activity dependent compensatory endocytosis at central synapses. J Neurosci 33: 16131–16145

Chauhan BK, Lou M, Zheng Y, Lang RA (2011) Balanced Rac1 and RhoA activities regulate cell shape and drive invagination morphogenesis in epithelia. Proc Natl Acad Sci U S A 108: 18289–18294

Chen X, Wu X, Wu H, Zhang M (2020) Phase separation at the synapse. Nat Neurosci 23: 301–310

Cingolani LA, Goda Y (2008) Actin in action: the interplay between the actin cytoskeleton and synaptic efficacy. Nat Rev Neurosci 9: 344–356

Clayton EL, Cousin MA (2009) The molecular physiology of activity-dependent bulk endocytosis of synaptic vesicles. J Neurochem 111: 901–914

Colgan LA, Yasuda R (2014) Plasticity of dendritic spines: subcompartmentalization of signaling. Annu Rev Physiol 76: 365–385

Dani A, Huang B, Bergan J, Dulac C, Zhuang X (2010) Superresolution imaging of chemical synapses in the brain. Neuron 68: 843–856

Daou P, Hasan S, Breitsprecher D, Baudelet E, Camoin L, Audebert S, Goode BL, Badache A (2014) Essential and nonredundant roles for Diaphanous formins in cortical microtubule capture and directed cell migration. Mol Biol Cell 25: 658–668

Deguchi Y, Harada M, Shinohara R, Lazarus M, Cherasse Y, Urade Y, Yamada D, Sekiguchi M, Watanabe D, Furuyashiki T et al (2016) mDia and ROCK Mediate Actin-Dependent Presynaptic Remodeling Regulating Synaptic Efficacy and Anxiety. Cell Rep 17: 2405–2417

Del Signore SJ, Kelley CF, Messelaar EM, Lemos T, Marchan MF, Ermanoska B, Mund M, Fai TG, Kaksonen M, Rodal AA (2021) An autoinhibitory clamp of actin assembly constrains and directs synaptic endocytosis. Elife 10

Delvendahl I, Vyleta NP, von Gersdorff H, Hallermann S (2016) Fast, Temperature-Sensitive and Clathrin-Independent Endocytosis at Central Synapses. Neuron 90: 492–498

Eisenmann KM, Peng J, Wallar BJ, Alberts AS (2005) Rho GTPase-formin pairs in cytoskeletal remodelling. Novartis Found Symp 269: 206–218; discussion 219-230

Engqvist-Goldstein AE, Drubin DG (2003) Actin assembly and endocytosis: from yeast to mammals. Annu Rev Cell Dev Biol 19: 287–332

Ferguson SM, Brasnjo G, Hayashi M, Wolfel M, Collesi C, Giovedi S, Raimondi A, Gong LW, Ariel P, Paradise S et al (2007) A selective activity-dependent requirement for dynamin 1 in synaptic vesicle endocytosis. Science 316: 570–574

Ferguson SM, Raimondi A, Paradise S, Shen H, Mesaki K, Ferguson A, Destaing O, Ko G, Takasaki J, Cremona O et al (2009) Coordinated actions of actin and BAR proteins upstream of dynamin at endocytic clathrin-coated pits. Dev Cell 17: 811–822

Gan Q, Watanabe S (2018) Synaptic Vesicle Endocytosis in Different Model Systems. Front Cell Neurosci 12: 171

Ganguly A, Tang Y, Wang L, Ladt K, Loi J, Dargent B, Leterrier C, Roy S (2015) A dynamic formin-dependent deep F-actin network in axons. J Cell Biol 210: 401–417

Gerth F, Japel M, Pechstein A, Kochlamazashvili G, Lehmann M, Puchkov D, Onofri F, Benfenati F, Nikonenko AG, Fredrich K et al (2017) Intersectin associates with synapsin and regulates its nanoscale localization and function. Proc Natl Acad Sci U S A 114: 12057–12062

Goode BL, Eck MJ (2007) Mechanism and function of formins in the control of actin assembly. Annu Rev Biochem 76: 593–627

Gormal R, Valmas N, Fath T, Meunier F (2017) A role for tropomyosins in activity-dependent bulk endocytosis? Mol Cell Neurosci 84: 112–118

Gu C, Chang J, Shchedrina VA, Pham VA, Hartwig JH, Suphamungmee W, Lehman W, Hyman BT, Bacskai BJ, Sever S (2014) Regulation of dynamin oligomerization in cells: the role of dynamin-actin interactions and its GTPase activity. Traffic 15: 819–838

Higashi T, Ikeda T, Shirakawa R, Kondo H, Kawato M, Horiguchi M, Okuda T, Okawa K, Fukai S, Nureki O et al (2008) Biochemical characterization of the Rho GTPase-regulated actin assembly by diaphanous-related formins, mDia1 and Daam1, in platelets. J Biol Chem 283: 8746–8755

Higashida C, Suetsugu S, Tsuji T, Monypenny J, Narumiya S, Watanabe N (2008) G-actin regulates rapid induction of actin nucleation by mDia1 to restore cellular actin polymers. J Cell Sci 121: 3403–3412

Hodge RG, Ridley AJ (2016) Regulating Rho GTPases and their regulators. Nature reviews Molecular cell biology 17: 496–510

Hori T, Eguchi K, Wang HY, Miyasaka T, Guillaud L, Taoufiq Z, Mahapatra S, Yamada H, Takei K, Takahashi T (2022) Microtubule assembly by tau impairs endocytosis and neurotransmission via dynamin sequestration in Alzheimer’s disease synapse model. Elife 11

Hua Y, Sinha R, Thiel CS, Schmidt R, Huve J, Martens H, Hell SW, Egner A, Klingauf J (2011) A readily retrievable pool of synaptic vesicles. Nat Neurosci 14: 833–839

Imoto Y, Raychaudhuri S, Ma Y, Fenske P, Sandoval E, Itoh K, Blumrich EM, Matsubayashi HT, Mamer L, Zarebidaki F et al (2022) Dynamin is primed at endocytic sites for ultrafast endocytosis. Neuron 110: 2815–2835 e2813

Keine C, Al-Yaari M, Radulovic T, Thomas CI, Valino Ramos P, Guerrero-Given D, Ranjan M, Taschenberger H, Kamasawa N, Young SM, Jr. (2022) Presynaptic Rac1 controls synaptic strength through the regulation of synaptic vesicle priming. Elife 11

Kessels MM, Qualmann B (2006) Syndapin oligomers interconnect the machineries for endocytic vesicle formation and actin polymerization. J Biol Chem 281: 13285–13299

Kitzing TM, Sahadevan AS, Brandt DT, Knieling H, Hannemann S, Fackler OT, Grosshans J, Grosse R (2007) Positive feedback between Dia1, LARG, and RhoA regulates cell morphology and invasion. Genes Dev 21: 1478–1483

Kononenko NL, Haucke V (2015) Molecular mechanisms of presynaptic membrane retrieval and synaptic vesicle reformation. Neuron 85: 484–496

Kononenko NL, Puchkov D, Classen GA, Walter AM, Pechstein A, Sawade L, Kaempf N, Trimbuch T, Lorenz D, Rosenmund C et al (2014) Clathrin/AP-2 mediate synaptic vesicle reformation from endosome-like vacuoles but are not essential for membrane retrieval at central synapses. Neuron 82: 981–988

Lash LL, Wallar BJ, Turner JD, Vroegop SM, Kilkuskie RE, Kitchen-Goosen SM, Xu HE, Alberts AS (2013) Small-molecule intramimics of formin autoinhibition: a new strategy to target the cytoskeletal remodeling machinery in cancer cells. Cancer Res 73: 6793–6803

Lawson CD, Ridley AJ (2018) Rho GTPase signaling complexes in cell migration and invasion. J Cell Biol 217: 447–457

Lopez-Hernandez T, Takenaka KI, Mori Y, Kongpracha P, Nagamori S, Haucke V, Takamori S (2022) Clathrin-independent endocytic retrieval of SV proteins mediated by the clathrin adaptor AP-2 at mammalian central synapses. Elife 11

Macia E, Ehrlich M, Massol R, Boucrot E, Brunner C, Kirchhausen T (2006) Dynasore, a cell-permeable inhibitor of dynamin. Dev Cell 10: 839–850

Marston DJ, Dickinson S, Nobes CD (2003) Rac-dependent trans-endocytosis of ephrinBs regulates Eph-ephrin contact repulsion. Nat Cell Biol 5: 879–888

McMahon HT, Boucrot E (2011) Molecular mechanism and physiological functions of clathrin-mediated endocytosis. Nature reviews Molecular cell biology 12: 517–533

Merrifield CJ, Perrais D, Zenisek D (2005) Coupling between clathrin-coated-pit invagination, cortactin recruitment, and membrane scission observed in live cells. Cell 121: 593–606

Miesenbock G, De Angelis DA, Rothman JE (1998) Visualizing secretion and synaptic transmission with pH-sensitive green fluorescent proteins. Nature 394: 192–195

Muller PM, Rademacher J, Bagshaw RD, Wortmann C, Barth C, van Unen J, Alp KM, Giudice G, Eccles RL, Heinrich LE et al (2020) Systems analysis of RhoGEF and RhoGAP regulatory proteins reveals spatially organized RAC1 signalling from integrin adhesions. Nat Cell Biol 22: 498–511

O’Neil SD, Racz B, Brown WE, Gao Y, Soderblom EJ, Yasuda R, Soderling SH (2021) Action potential-coupled Rho GTPase signaling drives presynaptic plasticity. Elife 10

Ogunmowo TH, Jing H, Raychaudhuri S, Kusick GF, Imoto Y, Li S, Itoh K, Ma Y, Jafri H, Dalva MB et al (2023) Membrane compression by synaptic vesicle exocytosis triggers ultrafast endocytosis. Nat Commun 14: 2888

Otomo T, Otomo C, Tomchick DR, Machius M, Rosen MK (2005) Structural basis of Rho GTPase-mediated activation of the formin mDia1. Mol Cell 18: 273–281

Peng GE, Wilson SR, Weiner OD (2011) A pharmacological cocktail for arresting actin dynamics in living cells. Mol Biol Cell 22: 3986–3994

Peng J, Kitchen SM, West RA, Sigler R, Eisenmann KM, Alberts AS (2007) Myeloproliferative defects following targeting of the Drf1 gene encoding the mammalian diaphanous related formin mDia1. Cancer Res 67: 7565–7571

Peng J, Wallar BJ, Flanders A, Swiatek PJ, Alberts AS (2003) Disruption of the Diaphanous-related formin Drf1 gene encoding mDia1 reveals a role for Drf3 as an effector for Cdc42. Curr Biol 13: 534–545

Piriya Ananda Babu L, Wang HY, Eguchi K, Guillaud L, Takahashi T (2020) Microtubule and Actin Differentially Regulate Synaptic Vesicle Cycling to Maintain High-Frequency Neurotransmission. J Neurosci 40: 131–142

Raimondi A, Ferguson SM, Lou X, Armbruster M, Paradise S, Giovedi S, Messa M, Kono N, Takasaki J, Cappello V et al (2011) Overlapping role of dynamin isoforms in synaptic vesicle endocytosis. Neuron 70: 1100–1114

Ramalingam N, Zhao H, Breitsprecher D, Lappalainen P, Faix J, Schleicher M (2010) Phospholipids regulate localization and activity of mDia1 formin. Eur J Cell Biol 89: 723–732

Renard HF, Simunovic M, Lemiere J, Boucrot E, Garcia-Castillo MD, Arumugam S, Chambon V, Lamaze C, Wunder C, Kenworthy AK et al (2015) Endophilin-A2 functions in membrane scission in clathrin-independent endocytosis. Nature 517: 493–496

Richards DA, Rizzoli SO, Betz WJ (2004) Effects of wortmannin and latrunculin A on slow endocytosis at the frog neuromuscular junction. J Physiol 557: 77–91

Ritter B, Ferguson SM, De Camilli P, McPherson PS (2017) A lentiviral system for efficient knockdown of proteins in neuronal cultures [version 1; referees: 2 approved]. MNI Open Res 1

Roux A, Uyhazi K, Frost A, De Camilli P (2006) GTP-dependent twisting of dynamin implicates constriction and tension in membrane fission. Nature 441: 528–531

Saffarian S, Cocucci E, Kirchhausen T (2009) Distinct dynamics of endocytic clathrin-coated pits and coated plaques. PLoS Biol 7: e1000191

Saheki Y, De Camilli P (2012) Synaptic vesicle endocytosis. Cold Spring Harbor perspectives in biology 4: a005645

Sakaba T, Kononenko NL, Bacetic J, Pechstein A, Schmoranzer J, Yao L, Barth H, Shupliakov O, Kobler O, Aktories K et al (2013) Fast neurotransmitter release regulated by the endocytic scaffold intersectin. Proc Natl Acad Sci U S A 110: 8266–8271

Sakaba T, Neher E (2003) Involvement of actin polymerization in vesicle recruitment at the calyx of Held synapse. J Neurosci 23: 837–846

Sankaranarayanan S, Atluri PP, Ryan TA (2003) Actin has a molecular scaffolding, not propulsive, role in presynaptic function. Nat Neurosci 6: 127–135

Schmied C, Soykan T, Bolz S, Haucke V, Lehmann M (2021) SynActJ: Easy-to-Use Automated Analysis of Synaptic Activity. Front Comp Sci-Switz 3

Shinohara R, Thumkeo D, Kamijo H, Kaneko N, Sawamoto K, Watanabe K, Takebayashi H, Kiyonari H, Ishizaki T, Furuyashiki T et al (2012) A role for mDia, a Rho-regulated actin nucleator, in tangential migration of interneuron precursors. Nat Neurosci 15: 373–380, S371-372

Shupliakov O, Bloom O, Gustafsson JS, Kjaerulff O, Low P, Tomilin N, Pieribone VA, Greengard P, Brodin L (2002) Impaired recycling of synaptic vesicles after acute perturbation of the presynaptic actin cytoskeleton. Proc Natl Acad Sci U S A 99: 14476–14481

Soykan T, Kaempf N, Sakaba T, Vollweiter D, Goerdeler F, Puchkov D, Kononenko NL, Haucke V (2017) Synaptic Vesicle Endocytosis Occurs on Multiple Timescales and Is Mediated by Formin-Dependent Actin Assembly. Neuron 93: 854–866 e854

Soykan T, Maritzen T, Haucke V (2016) Modes and mechanisms of synaptic vesicle recycling. Curr Opin Neurobiol 39: 17–23

Watanabe S, Boucrot E (2017) Fast and ultrafast endocytosis. Curr Opin Cell Biol 47: 64–71

Watanabe S, Rost BR, Camacho-Perez M, Davis MW, Sohl-Kielczynski B, Rosenmund C, Jorgensen EM (2013) Ultrafast endocytosis at mouse hippocampal synapses. Nature 504: 242–247

Watanabe S, Trimbuch T, Camacho-Perez M, Rost BR, Brokowski B, Sohl-Kielczynski B, Felies A, Davis MW, Rosenmund C, Jorgensen EM (2014) Clathrin regenerates synaptic vesicles from endosomes. Nature 515: 228–233

Willems J, de Jong APH, Scheefhals N, Mertens E, Catsburg LAE, Poorthuis RB, de Winter F, Verhaagen J, Meye FJ, MacGillavry HD (2020) ORANGE: A CRISPR/Cas9-based genome editing toolbox for epitope tagging of endogenous proteins in neurons. PLoS Biol 18: e3000665

Wu LG, Chan CY (2022) Multiple Roles of Actin in Exo- and Endocytosis. Front Synaptic Neurosci 14: 841704

Wu XS, Lee SH, Sheng J, Zhang Z, Zhao WD, Wang D, Jin Y, Charnay P, Ervasti JM, Wu LG (2016) Actin Is Crucial for All Kinetically Distinguishable Forms of Endocytosis at Synapses. Neuron

Wu Y, O’Toole ET, Girard M, Ritter B, Messa M, Liu X, McPherson PS, Ferguson SM, De Camilli P (2014) A dynamin 1-, dynamin 3- and clathrin-independent pathway of synaptic vesicle recycling mediated by bulk endocytosis. Elife 3: e01621

Xu Y, Moseley JB, Sagot I, Poy F, Pellman D, Goode BL, Eck MJ (2004) Crystal structures of a Formin Homology-2 domain reveal a tethered dimer architecture. Cell 116: 711–723

Zhou L, McInnes J, Wierda K, Holt M, Herrmann AG, Jackson RJ, Wang YC, Swerts J, Beyens J, Miskiewicz K et al (2017) Tau association with synaptic vesicles causes presynaptic dysfunction. Nat Commun 8: 15295

